# The essential role of connective-tissue cells during axolotl limb regeneration

**DOI:** 10.1101/2025.03.30.645595

**Authors:** Damián García-García, Dunja Knapp, Katelyn Jamwal, Riley E. Grindle, Heath Fuqua, Ryan P. Seaman, Markus Sujansky, Minjoo Kim, Sergej Nowoshilow, Maria Novatchkova, Fred W. Kolling, Joel H. Graber, Prayag Murawala

**Affiliations:** Mount Desert Island Biological Laboratory (MDIBL), Bar Harbor, Maine, USA; Hannover Biomedical Research School, Hannover Medical School. Hannover, Germany; Center for Regenerative Therapies (CRTD), Technische Universität Dresden, Dresden, Germany; Research Institute of Molecular Pathology, Vienna, Austria; Institute of Molecular Biotechnology (IMBA), Vienna, Austria; Dartmouth Cancer Center, Dartmouth Geisel School of Medicine, Lebanon, New Hampshire, USA; Department of Nephrology and Hypertension, Hannover Medical School, Hannover, Germany

## Abstract

Axolotls (*Ambystoma mexicanum*) are known for their remarkable limb-regeneration abilities, which involve the formation of the blastema, a specialized structure consisting of progenitor cells contributed by all major tissues of the limb. Lateral plate mesoderm (LPM)-derived connective tissue (CT) cells dedifferentiate and play a critical role in blastema formation and subsequent limb regeneration. However, the complexity of the blastema’s cellular composition and the extent of CT participation and necessity have not been rigorously explored. To address this gap, we conducted spatial transcriptomics using a select array of probes, revealing that CT cells constitute up to 75% of the blastema cells. Genetic ablation of CT cells significantly delays or truncates limb regeneration, underscoring their necessity during this process. Analysis of the molecular profile during blastema formation identifies a temporally ordered gene expression pattern reflecting sequential phases of CT cell dedifferentiation, proliferation, and redifferentiation. Finally, a cell-cell communication inference identifies a CT-cell-autonomous FGFR1/AKT1 signaling axis as a cell-intrinsic molecular requirement for limb regeneration. Together, our work reaffirms the central role of CT cells in axolotl limb regeneration and lays the foundation for identifying molecular mechanisms that govern blastema formation during the initial phases of limb regeneration.

## INTRODUCTION

The axolotl (*Ambystoma mexicanum*) is a salamander renowned for its remarkable regenerative abilities, which include the restoration of various tissues and organs, such as limbs, the heart, kidneys, and the central nervous system, among others ^1–4^. Limb amputation initiates a series of cellular and molecular processes that facilitate regeneration of the lost body part. Wound healing commences immediately after injury, and a new epidermis covers the amputation site within the first 24 hours ^5,6^. Subsequently, limb progenitors emerge through cell dedifferentiation and cellular activation from all major limb tissues that ultimately coalesce beneath the wound site, forming a heterogeneous pool of undifferentiated cells termed the blastema ^7–9^. As regeneration progresses, the limb blastema progenitors proliferate, redifferentiate, and reorganize into distinct tissues, ultimately forming a fully functional limb.

The cellular composition of the blastema evolves throughout the process of limb regeneration. In the initial stages of wound healing, a cytokine-rich microenvironment promotes the infiltration of leukocytes, including macrophages, neutrophils, B cells, and T cells, and this persists until the mid-bud blastema stage ^10–13^. At this time, leukocytes begin to leave the amputation site, and connective tissue (CT)-derived cells gradually dominate the blastema’s cellular identity. By the mid-bud blastema stage, CT-derived cells dedifferentiate, transforming into primitive, multipotent progenitor cells, which proliferate and, at the late-bud blastema stage, start to redifferentiate into cartilage, bone, and non-skeletal connective tissues ^8^. Thin fibers from the peripheral nervous system are detected in the blastema by the mid-bud stage and become more prominent during the palette and limb growth stages ^14^. Similarly, endothelial cells enter the blastema by the mid-bud stage, with visible blood vasculature appearing by the late-bud stage.

Finally, muscle progenitor cells also contribute to the blastema, beginning in the mid-bud blastema stages, differentiating into the muscle fibers in the regenerated limb ^9,15^.

While we have a general understanding of blastema composition during regeneration, the precise dynamics of different progenitor populations during blastema formation and growth remain unexplored. A quantitative analysis of cell identities at various stages of blastema development would provide a better understanding of the major events occurring during regeneration. This categorization would also facilitate future studies on the essential crosstalk taking place between cell types throughout these stages.

To address this gap, we performed spatial profiling of 100 transcripts at different stages during blastema development in axolotl. Overall, we identified distinct cell types and observed that immune cells are the predominant constituents of the early-bud blastema, whereas CT-derived cells dominate in the late-bud blastema stage (i.e., 14 dpa). Based on these results, we reasoned that CT cells might be key players in limb regeneration, including blastema formation. Accordingly, we quantified CT-derived cell abundance during blastema formation using *in viv*o lineage tracing and spatial transcriptomics. Both technologies confirmed that CT-derived cells account for approximately 75 % of the late-bud blastema cells. Next, we developed a genetic tool to eliminate CT-derived cells during limb regeneration, utilizing the nitroreductase (NTR)/metronidazole (MTZ) cell-ablation system. Use of this approach resulted in delayed or truncated limb regeneration, highlighting the necessity for these cells. Additionally, we complemented previously published single-cell RNA sequencing (scRNA-seq) data by performing expression profiling of CT-derived cells using microarray. This allowed us to gain novel insights into the molecular dynamics of CT cells at different stages of limb regeneration. We also performed cell-cell communication inference using CellChat ^16^, which identified a CT-cell autonomous FGFR1/AKT1 signaling axis as a key regulator of limb regeneration. To validate this, we performed functional interrogation via pharmacological inhibition, downstream pathway dissection, and dominant-negative overexpression and rescue with constitutive-active protein overexpression to demonstrate that FGFR1-AKT1 signaling is an essential cell-intrinsic molecular requirement for limb regeneration.

## RESULTS

### Limb blastema is composed primarily of CT-derived cells

CT-derived cells constitute the largest and most dynamically expanding cell population of the axolotl limb blastema ^8^. To characterize these dynamics quantitatively with spatial resolution, we employed the 10X Xenium platform to profile 100 selected genes across key stages of limb regeneration (**Figure 1).** Unlike previously published single-cell RNA sequencing studies in the regenerating blastema ^8,11,17^, Xenium preserves the spatial context, allowing resolution of cellular identities directly within the tissue architecture. Furthermore, Xenium offers a key technical advantage over previous Visium-based spatial transcriptomics in the regenerating blastema ^18^ by assigning the gene expression to individual cells detected by DAPI-based nuclear segmentation, rather than to large, spatially barcoded spots that can encompass multiple cells and obscure true single-cell resolution. The 100 selected genes include previously published tissue-specific markers ^8,11,18^ (**Full list in Supplementary Data 1**), and we analyzed their expression in multiple stages, including the mature limb (uninjured limb), Apical Ectodermal Cap (AEC) formation (3 days post-amputation [dpa]), early-bud (5 and 7 dpa), mid-bud (10 dpa), and late-bud (14 dpa) blastema stages, all sampled from 5-cm-long axolotls (**Figure 1a-b**). Using graph-based clustering analysis, we identified seven distinct cell clusters common across all the five stages corresponding to apical and basal skin keratinocytes, blood vessels, nerves, muscles, macrophages, and bone (**Figure 1b; Supplementary Figures 1-9**). We also detected clusters specific to limb regeneration and blastema development, which include the AEC cells cluster and the segmentation of CT cells into blastema fibroblasts and stump fibroblasts sub-clusters. Periskeletal cells were detected from the mature limb to 7 dpa and osteo/chondro-progenitor cells from 10 dpa onwards. Additionally, we detected a Langerhans cell cluster at the mature limb, 7 and 14 dpa, and goblet cells in the mature limb, and 3, 7, and 14 dpa. Interestingly, at a the 14 dpa timepoint, blastema fibroblasts could be further segmented into distal, proximal, medial, and posterior cells, along with the emergence of myogenic progenitor cells. Finally, we detected a cluster of general immune cells at the mature limb, and 3, 7, and 14 dpa. However, this cell cluster could not be precisely classified due to the limited number of immune-cell marker probes included in the analysis.

**Figure 1.**
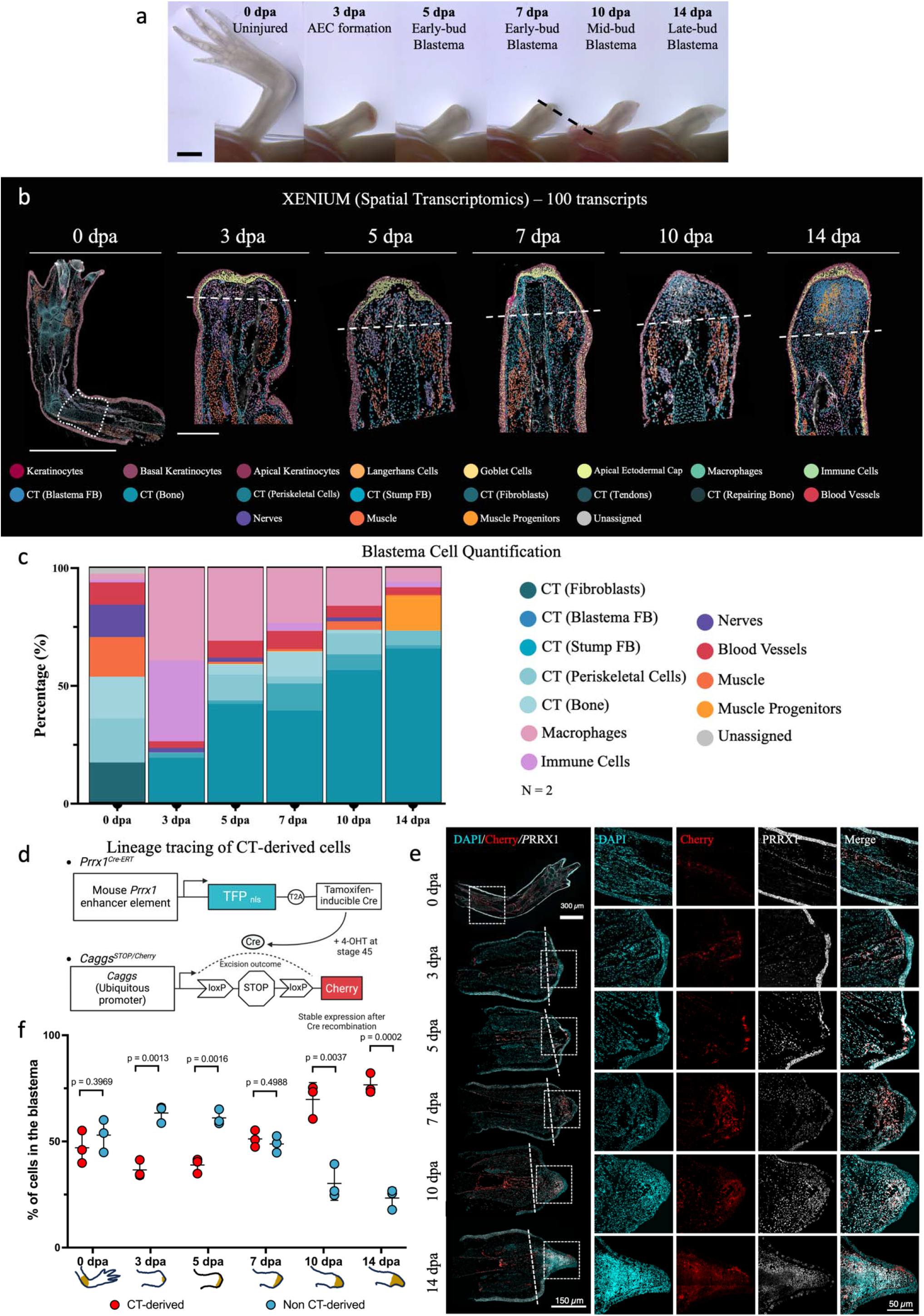
Connective tissue (CT)-derived cells are the major constituent of the blastema. **a,** Stereoscopic images illustrating the progressive stages of limb regeneration, visualized via widefield microscopy. **b,** Xenium analysis performed in an uninjured limb and at 3, 5, 7, 10, and 14 dpa. Cellular identities were determined by integrating nuclear segmentation with transcript presence, identifying distinct cell clusters, each one represented by a different color. Scale bars: 5 mm for the uninjured limb and 1 mm for the subsequent time points. The dotted white line marks the amputation plane. The amputation plane was determined based on the loss of continuity of muscle or nerve tissues. In the uninjured limb, the dotted line represents the area used for quantification in Fig. 1c. **c,** Quantification of the percentage of cell types within the blastema region (tissue distal to the amputation plane) excluding the epidermis. (n=1) **d**, Schematic representation of the experimental design detailing lineage tracing experiments conducted using converted *Prrx1^Cre-ERT^;Caggs^STOP/Cherry^*axolotls. **e**, Immunofluorescence analysis showing Cherry (red) or anti-PRRX1 (white) expression in longitudinal sections at key regenerative time points. Cells expressing Cherry, anti-PRRX1, or both were counted as CT-derived cells. Dotted lines delineate the amputation planes in each case. **f**, Quantification of CT-derived and non-CT-derived cells within the blastema region (n =3). The quantification region extends from the distal area following the amputation plane, excluding the limb epidermis. Statistical significance was determined using unpaired t-tests, with corresponding p-values annotated in the graph for each comparison.

Using gene enrichment analysis of the 100-probe Xenium set, we identified the genes most strongly expressed by each cell type (**Supplementary Figures 1-6, e in each case and Supplementary Data 2, and Supplementary Figure 8**). The goblet-cell cluster is characterized primarily by the expression of *Des* and *Trpv1*. Skin keratinocytes are defined by *Serpin1*, *Dll1*, and *Rbp7* expression, whereas *Mmp13*, *Sall4*, and *Wnt3a* are the predominant markers of the AEC. Blood vessels exhibit unique expressions of *Cygb*, *CD34*, and *Lorf2*, while nerve bundles are identified by *Sox10* and *Egr2* expression. Muscles are defined primarily by *Myf5* expression, whereas the macrophage cluster is characterized by *C1qa*, *C1qb*, and *Lamp3* expression, and the general immune-cell cluster is identified by *Pere* and *Arg1* expression. All CT-derived clusters are characterized by *Prrx1* expression. This includes the stump fibroblasts, which are distinguished by the expression of *Agra2* and *Dio3*, while blastema fibroblasts are defined by expression of *Kazald1*, *Grem1*, *Tgb1*, and *Crabp2*. Periskeletal cells are identified by *Dkk1* and *Scx* expression, distinguishing them from the *Mmp13*, *Sox9*, and *Grid1* expressing skeletal cells. Next, we quantified the cellular composition of the blastema, excluding the epidermal layer at each stage (**Figure 1b, the area distal to the dashed line at each stage**). Our analysis revealed that the CT-derived population exhibited the most significant increase in relative abundance over the five stages, comprising 21% of the cells at 3 dpa and gradually rising to a peak of 73% by 14 dpa. The immune-cell populations exhibited an inverse temporal pattern, collectively comprising 73% of the quantified cells at 3 dpa (39% were macrophages and 34% were other immune cells), and decreasing to a minimum of 8.6% by 14 dpa (6.2% were macrophages and 2.4% were other immune cells) (**Figure 1c**). This suggests that the role of macrophages and other types of immune cells is important primarily in the early stages of limb regeneration. In contrast to these dynamic changes, the proportion of blood vessels, nerves, and mature muscle cells remained relatively stable, each maintaining low levels (<8%) throughout the five stages (**Figure 1 b-c**).

Considering the importance of CT cells in limb identity and skeletal patterning, we next performed a validation experiment to determine the extent of CT cell participation during regeneration. To accomplish this, we performed an *in vivo* lineage tracing by crossing (I) an axolotl line expressing a tamoxifen-inducible CreER recombinase specifically in the limb CT cells driven by a mouse *Prrx1* enhancer element (*Prrx1^Cre-ERT^*) with (II) a *Caggs^STOP/Cherry^* axolotl line, where fluorescent Cherry protein is constitutively expressed upon tamoxifen-induced recombination. The resulting *Prrx1^Cre-ERT^Caggs^STOP/Cherry^*line tags CT-derived cells permanently with Cherry protein, which, in combination with anti-PRRX1 antibodies, enabled us to comprehensively identify CT-derived cells in the mature limb, stump, or blastema (**Figure 1d**). Quantification of CT cells in the uninjured limb revealed similar proportions of CT-derived and non-CT-derived cells. In addition, the analysis of the blastema cells confirmed a consistent enrichment of CT-derived cells over time, reaching 76.6 ± 4.8% of the total blastema cell population at 14 dpa (**Figure 1e-f**), showing a strong correlation with the results from Xenium spatial transcriptomics. Next, we analyzed cell dynamics within the source zone, defined as the 500-µm region proximal to the amputation plane at each time point. In this zone, we observed an increase in the abundance of CT-derived cells during regeneration, peaking at 7 dpa and remaining elevated through 14 dpa (**Supplementary Figure 10**), which correlates with previous reports ^19^.

Collectively, our spatial transcriptomics analysis and genetic lineage tracing experiments revealed distinct temporal dynamics of cellular participation during blastema development. Immune cells, particularly macrophages, are predominant during the early stages of regeneration, whereas muscle progenitors are notably present at the late-bud stage, the latest point analyzed, whereas endothelial cells and nerve components show only a minor participation. Importantly, PPRX1^+^-derived CT cells progressively accumulate and peak at 14 dpa, becoming the primary cellular component of the blastema.

### CT-derived cells are necessary for limb regeneration

Based on our demonstration of the progressive dominance of CT-derived cells in the blastema, we asked whether these cells are strictly required for regeneration and whether other cell types could compensate for their loss. To address this directly, we generated a *loxP*-dependent transgenic line with a conditional expression of the NTR2.0 gene (*CAGGS:Lp-nBFP-Lp-Cherry-T2A-NTR2.0*, hereinafter referred to as *Caggs^nBFP/Cherry-NTR2^)*. NTR2.0 catalyzes the reduction of the innocuous prodrug metronidazole (MTZ), into a cytotoxic product that induces a cell-autonomous cellular death ^20,21^.

To establish the optimal dosage of MTZ, we conducted survival assays on d/d axolotls at two different ages: stage 45 larvae, and 5-cm-long juveniles. We identified optimal working concentrations of 10 mM and 20 mM for the two groups, respectively, achieving a survival rate of over 90% after one day of treatment and more than 80% after two days for both groups (**Supplementary Figure 11**). A genetic cross between *Prrx1^Cre-ERT^* and *Caggs^nBFP/Cherry-NTR2^*axolotls allowed us to express NTR2.0 specifically in CT progenitors during embryonic limb development. We first examined the effect of CT-progenitor ablation during forelimb development by administering MTZ at larvae stage 47. All MTZ-treated animals (100%) displayed forelimb abnormalities, including truncated limb development or an incomplete zeugopod segment (**Supplementary Figure 12**).

Next, we assessed the impact of CT-derived-cell ablation on limb regeneration by performing an overnight MTZ treatment at 14 dpa, corresponding to the highest CT-derived-cell abundance, in 5-cm long *Prrx1^Cre-ERT^Caggs^nBFP/Cherry-NTR2^*axolotls (**Figure 2a-b**). Immunodetection of the cell-death marker Caspase-3 indicated abundant apoptosis in NTR2.0 (Cherry)-expressing blastema cells (**Figure 2c**), confirming that our NTR2.0/MTZ system functions efficiently in axolotls. While there was no overall effect on body size after MTZ treatment, we monitored limb regeneration from 0 dpa to 60 dpa and found that CT-ablated limbs exhibited a smaller limb-to-body-length ratio compared to controls (**Figure 2 d-e**). Phenotypic analysis of CT-ablated limbs revealed two distinct outcomes: 1) a delayed regeneration phenotype, characterized by a fully patterned limb but smaller than that of controls, and 2) a truncated phenotype, characterized by imperfect zeugopod development and formation of a maximum of two digits in the autopod (**Figure 2f**).

**Figure 2.**
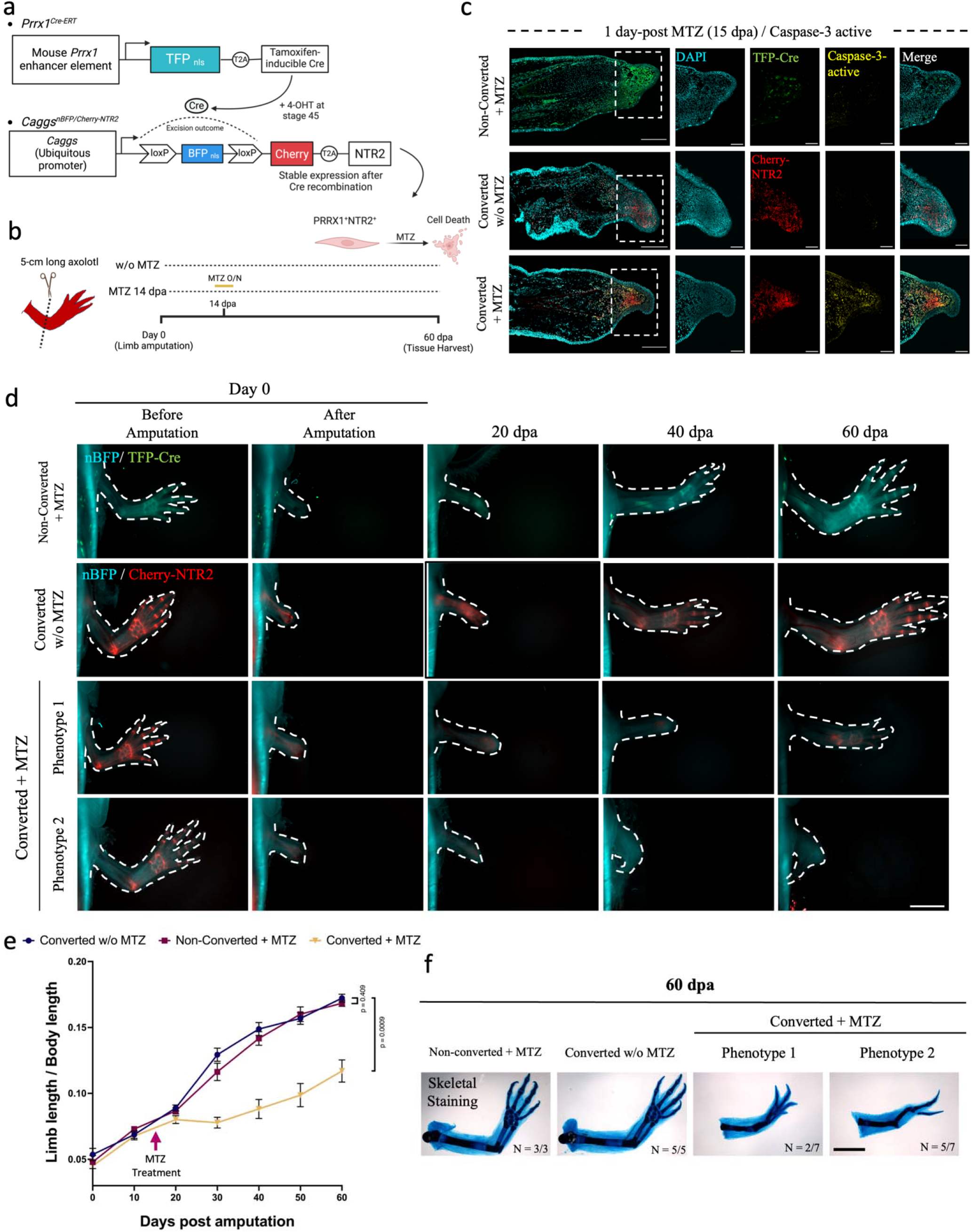
Connective tissue (CT)-derived cells are essential for limb regeneration. **a,** Schematic representation of the genetic cross between *Prrx1^Cre-ERT^* and *Caggs^nBFP/Cherry-NTR2^*to generate converted *Prrx1^Cre-ERT^;Caggs^nBFP/NTR2^* axolotls. Hatched larvae were treated with 4-OHT at stage 45 to induce gene recombination. **b**, Schematic of cell ablation experiment. At 14 dpa, animals were treated overnight with 20 mM metronidazole (MTZ) and allowed to regenerate until 60 dpa. **c,** Analysis of CT cell death at 15 dpa (1-day post-MTZ treatment) using cell death marker Caspase 3 (yellow). Longitudinal sections of 2 control conditions: 1) non-converted [Cherry^-^NTR2^-^] with MTZ treatment, and 2) converted [Cherry^+^NTR2^+^ in red] animals without MTZ treatment, as well as the experimental condition converted [Cherry^+^NTR2^+^ in red] with MTZ treatment, are shown. Scale bar = 500 µm (n = 2 each). The dotted lines indicate the magnified area. Scale bar in the magnified image= 200 µm. **d**, Stereoscopic images tracking limb regeneration over 60 days visualized via widefield microscopy. Fluorescence from non-converted *Prrx1^Cre-ERT^;Caggs^nBFP/NTR2^* (blue/green) and converted *Prrx1^Cre-ERT^;Caggs^nBFP/NTR2^* (blue/red) animals are displayed. Dotted lines mark the limb periphery. Scale bar = 2 mm. **e**, Quantitative assessment of limb length/body length ratio of the regenerated limb over time. The quantified region for limb length extends from the axolotl torso to the most distal part of the limb. The arrow indicates MTZ treatment at 14 dpa. Data are represented as mean ± SD (n = 3 for non-converted axolotls and n = 12 for each condition using converted axolotls). Statistical significance was determined at 60 dpa using two-way unpaired t-tests, with corresponding p-values annotated in the graph for each comparison. **f**, Phenotypic analysis of regenerated limb at 60 dpa using alcian blue/alizarin red staining for skeletal elements. The number of limbs analyzed per condition is mentioned in the figure. Two distinct phenotypes were observed followed CT cell ablation.

To determine whether the impact of CT ablation on limb regeneration is time-dependent, we applied single MTZ treatments at earlier time points (5, 7, or 10 dpa). We observed delayed limb regeneration, with smaller, fully patterned regenerated limbs at 60 dpa in all cases (**Supplementary Figure 13**). A more comprehensive analysis indicated that a later MTZ treatment (i.e., 14 dpa) resulted in a more severe phenotype (**Supplementary Figure 14**). Collectively, these experiments establish CT-derived cells as essential drivers of both axolotl limb development and regeneration, and reveal that their necessity scales with blastema maturity, motivating a deeper investigation of their molecular program.

### Transcriptional profiling of CT-derived cells reveals sequential, stage-specific gene-expression programs during blastema formation

Overall, CT is a major contributor to the blastema (**Figure 1**), necessary for blastema formation (**Figure 2**) and responsible for carrying positional and patterning information ^8,22^. Hence, a comprehensive understanding of the transcriptional transitions of CT cells through critical stages of blastema formation is essential for advancing our knowledge of limb regeneration. In previous work, we described the scRNA-seq profile of CT-derived cells during limb regeneration ^8^. While this scRNA-seq is informative on a global scale, the shallow coverage of reads per gene and limited number of genes detected in scRNA-seq data in general constrain its utility for analyzing individual gene expression in CT cells. In addition, the bulk-transcriptional profile of the blastema ^23^ also has limited value, as our findings (**Figure 1**) indicate that the complex cellular composition of the blastema is a constantly evolving phenomenon. To address these limitations and provide complementary insight into the CT cells transcriptome, we performed microarray expression profiling of sorted Cherry^+^ cells from *Prrx1^Cre-ERT^Caggs^STOP/Cherry^* animals at 1, 3, 5, 7, 10, and 14 dpa (**Figure 3a**).

**Figure 3.**
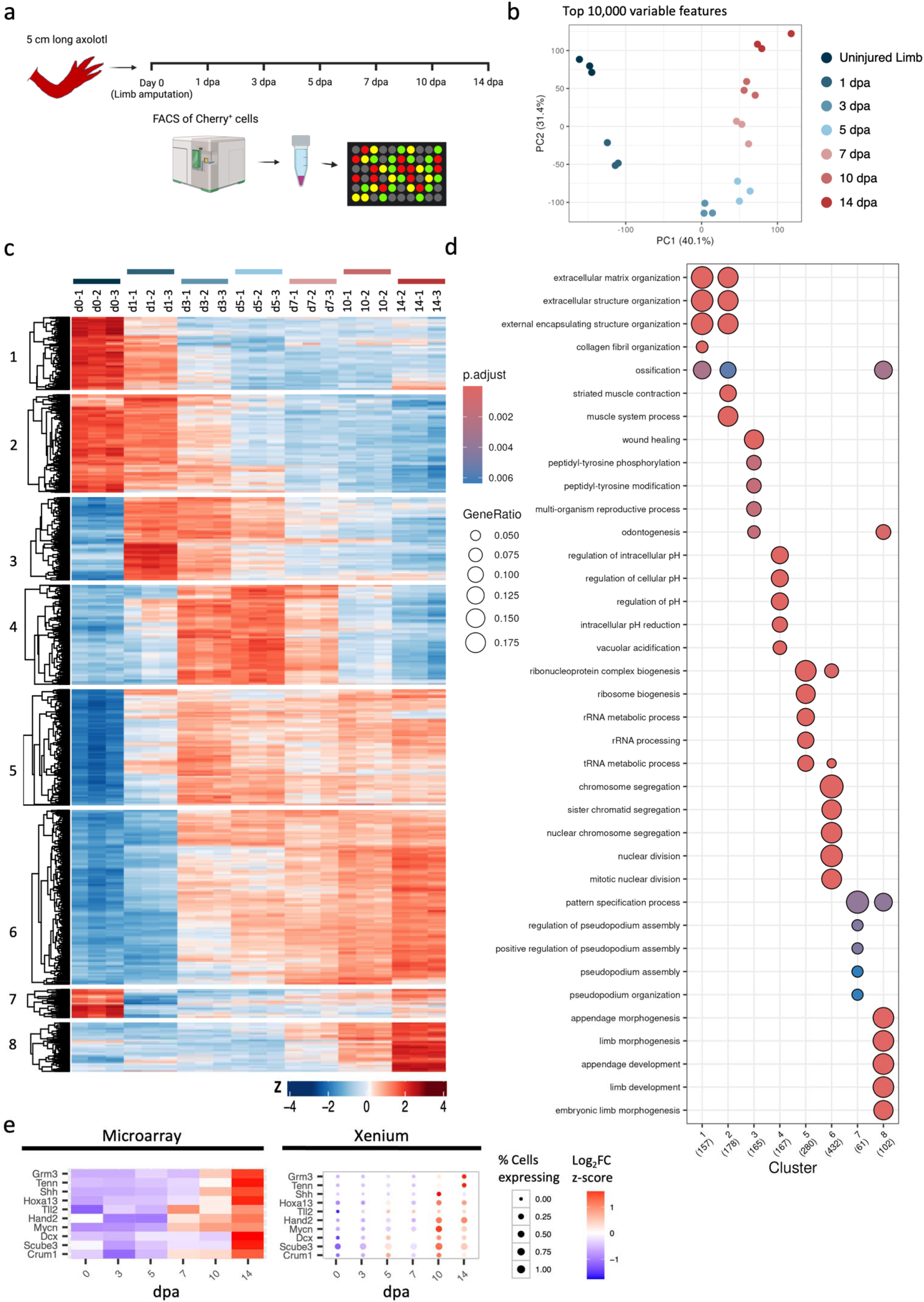
Microarray analysis of sorted CT cells reveals molecular dynamics during blastema formation. **a**, Schematic representation of the experimental design for the experiments conducted with converted *Prrx1^Cre-ERT^;Caggs^STOP/Cherry^*axolotls. **b**, Principal Component Analysis (PCA) of sorted CT cells, based on the top 10,000 most variable features of the microarray analysis (n = 3 per time point). **c**, Heatmap visualization of the microarray analysis, displaying all of the differentially expressed genes. Each row corresponds to a gene, while each column represents a biological sample. The color scale indicates fold-change in expression relative to the uninjured limb. Genes were categorized into eight major expression patterns, capturing both upregulated and downregulated trends over time. **d**, Gene ontology (GO) analysis for each of the eight clusters, identifying key biological processes associated with the observed expression patterns. **e**, Comparison of gene expression over time between microarray (left) dataset of FACS-sorted CT-derived cells and Xenium (right) of *in-silico*-sorted blastema fibroblast clusters. In both the microarray heatmap and Xenium dot plot, every row represents a gene, and every column a time point analyzed. Xenium’s dot size represents the percentage of blastema fibroblasts expressing the corresponding gene. In both microarray and Xenium datasets, the color scale indicates fold-change in expression relative to the uninjured limb (Time 0).

Principal Component Analysis (PCA) of the 10,000 most variable features demonstrated consistent clustering of the biological replicates at each analyzed time point, indicating strong reproducibility (**Figure 3b**). Clustering analysis of the microarray dataset revealed eight major dynamic expression clusters (**Figure 3c**), which, combined with a Gene Ontology (GO) enrichment analysis, revealed key biological features in the CT-derived cells across regenerative stages (**Figure 3d**). Clusters 1 and 2 are characterized by rapid early downregulation of genes associated with extracellular matrix structural and collagen organization, consistent with a simultaneous halt in tissue-maintenance programs and switch towards pro-regenerative states immediately after amputation. Cluster 3 showed a transient early peak in expression between 1 and 3 dpa, enriched for genes associated with wound healing. Cluster 4 showed a later transient peak of expression between 3 and 7 dpa, associated with intracellular pH regulation and vacuolar acidification; this may reflect an adaptive response to the acidic environment generated by macrophage-mediated ECM remodeling at the stump. Cluster 5 maintained relatively constant expression after limb amputation and was associated with ribosomal biogenesis, rRNA processing, and ncRNA processing, while cluster 6 showed a gradual upregulation of genes primarily associated with chromosomal segregation and DNA replication, reflecting the rapid proliferative expansion of CT-derived cells during blastema formation (**Figure 1**). Finally, clusters 7 and 8 showed a late-stage upregulation of transcripts involved in mesenchymal-cell differentiation, cell-fate commitment, embryonic limb development, and limb morphogenesis, suggesting that a defined temporal window exists between 10 and 14 dpa (late-bud blastema stage) during which CT-derived cells start the re-differentiation program.

To assess the robustness of the microarray dataset, we compared it with our previously published scRNA-seq dataset at the three time points in common to both platforms: uninjured limb, 5 dpa, and 10 dpa. For this comparison, we aggregated the scRNA-seq counts into a pseudobulk dataset. Of the total number of 21,881 genes detected by the two technologies, 10,847 (50%) were detected by both, while 2,184 (10%) were detected only by the microarray and 8,850 (40%) only by pseudobulk analysis (**Supplementary Figure 15**). Differential expression analysis revealed a systemic 2-fold higher log2FC values in the pseudobulk data compared to the microarray at both time points, consistent with the broader dynamic range of scRNA-seq technology (**Supplementary Figure 16 a-b**). Despite this difference in magnitude, the two platforms showed strong concordance in the detection of differentially expressed genes (DEG), with 38% of upregulated DEGs and 42% of downregulated DEGs identified by the two technologies at both 5 dpa and 10 dpa (**Supplementary Figure 16 c-f**). Notably, 55% of the upregulated DEGs at 5 dpa and 10 dpa were detected exclusively by the microarray, predominantly among lowly expressed transcripts, emphasizing the microarray’s advantage in detecting subtle yet biologically significant gene expression changes.

To test the accuracy of the microarray dataset, we compared the expression levels obtained from this technology with those detected in the blastema fibroblast cluster from the Xenium platform (**Figure 1c**). This comparison revealed a strong correlation in gene-expression dynamics over time (**Figure 3e, Supplementary Figure 17**). Individual Pearson correlation analyses for the 100 transcripts detected by both technologies showed that over 38% of the genes have a Pearson coefficient in the range of 0.7 to 1.0 indicating a strong correlation between the microarray and Xenium technologies, and 19% of the genes have a Pearson coefficient in the range of 0.3 to 0.7, indicating a moderate correlation (**Supplementary Figure 18**). These correlations support the microarray dataset as a reliable resource for future research studies.

To support the research community in exploring and analyzing individual transcripts from the microarray, we have developed an interactive web application (microarrayct-murawalalab.mdibl.org). This publicly available resource provides transcript IDs, sequences, and probes used for detection (**Supplementary Video 1**). By making this tool accessible, we aim to facilitate future studies and advance our collective understanding of the biological processes occurring in CT-derived cells after limb amputation. We anticipate that this application will be a valuable and widely used resource for researchers investigating limb regeneration and connective tissue biology.

### CellChat analysis reveals the FGFR1/AKT signaling axis as a major regulator of CT-cell function during limb regeneration

We next investigated the intercellular signaling landscape of CT-derived cells by using CellChat ^16^ to analyze the previously published scRNA-seq data. Our rationale for using this approach is twofold: we have established a strong concordance between the microarray and scRNA-seq datasets (see above), and the scRNA-seq data capture all blastema cell types and constitute the required input for cell-cell communication inference. We performed the CellChat analysis across the three stages between which we have previously compared them with the microarray dataset: the uninjured limb, the 5-dpa blastema, and the 10-dpa blastema. Global analysis of predicted intercellular signaling networks revealed a marked increase in the number and strength of interactions from the uninjured limb through 5 and 10 dpa (**Figure 4a**), reflecting the progressive amplification of cellular crosstalk during blastema formation. At 5 dpa, the strongest predicted communication involved the immune-cell cluster, which diminished by 10 dpa and shifted substantially towards CT cell-clusters, consistent with the progressive dominance of CT-derived cells at this stage (**Figure 1**). Given that CT-derived cells constitute the majority of the blastema cellular population and are necessary for regeneration (**Figures 1-2**), we focused our subsequent CellChat analysis on the autocrine and paracrine communication between various CT subgroups. Ranking CT-CT communication pathways by relative information flow identified fibronectin (Fn1), thrombospondin (THBS), periostin, and Ephrin B (EphB) signaling among the most prominently upregulated pathways at both 5 dpa and 10 dpa, and detected none of these three pathways in the uninjured limb (**Figure 4b**), suggesting their *de novo* activation of these pathways after amputation Among the pathways already active in the uninjured limb, Tenascin (Tnc), NCAM, JAM, and Col1a1 showed the greatest amplification at 5 dpa. Those signaling pathways sustained their high levels of communication at 10 dpa, while FGF and PDGF signaling emerged as additional highly ranked CT-CT pathways specifically at 10 dpa, suggesting a progressive layering of intercellular signals as the blastema forms.

**Figure 4.**
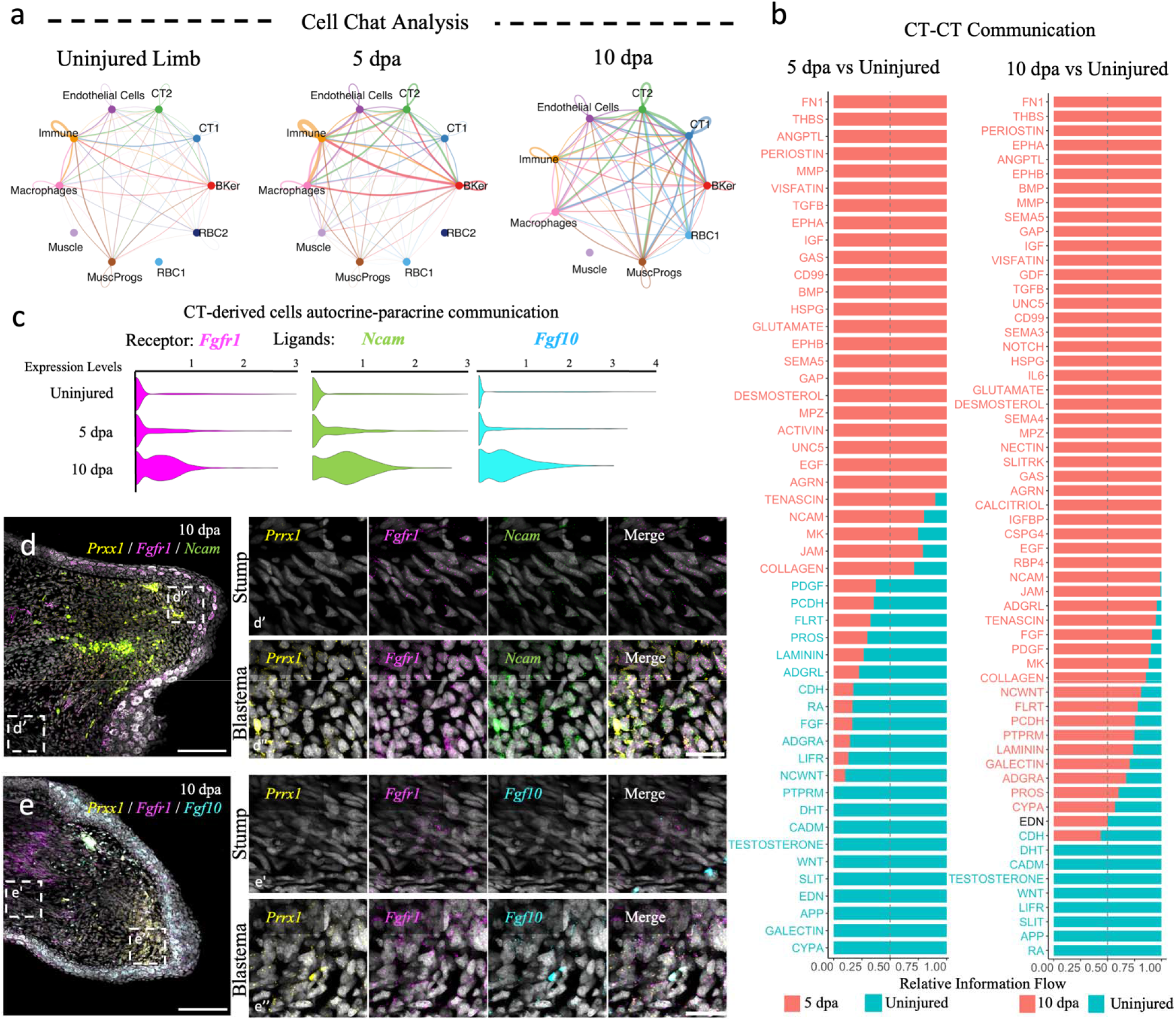
CellChat identifies FGFR1 as a key CT-cell signaling axis required for blastema formation and limb regeneration. **a**, Circle plots depicting the global intercellular communication networks inferred by CellChat analysis in the uninjured limb, at 5 dpa, and at 10 dpa. Line thickness and color indicate the strength and directionality of predicted signaling interactions between cell populations. CT, Connective Tissue; MusProgs, Muscle Progenitors; RBC, Red Blood Cells; BKer, Basal Keratinocytes. **b**, Information flow charts ranking the CT-CT cell autocrine and paracrine communication pathways by relative information flow at 5 dpa (left) and 10 dpa (right) compared to the uninjured limb. Orange bars indicate pathways active at 5 dpa or 10 dpa; cyan bars indicate pathways active in the uninjured limb. Pathways are organized in three groups from top to bottom: (1) pathways exclusively detected at 5 dpa or 10 dpa (orange only), reflecting their *de novo* activation after amputation; (2) pathways present in the uninjured limb with augmented information flow at 5 dpa or 10 dpa (both orange and cyan), reflecting amplification of pre-existing signals; and (3) pathways exclusively detected in the uninjured limb (cyan only), reflecting signals downregulated after amputation. **c**, Violin plots showing the expression of *Fgfr1*, *Ncam*, and *Fgf10* in CT-cell clusters across the uninjured limb, 5 dpa, and 10 dpa, as inferred from CellChat analysis. **d-e,** HCR validation of transcript co-expression of *Fgfr1* (magenta)/*Ncam* (d) or *Fgfr1/Fgf10* (e) in *Prrx1* (yellow) CT cells. The left column shows the blastema section; scale bar, 250 μm. Remaining columns show magnified views of the stump (d’, e’) and blastema regions (d’’, e’’); scale bar, 50 μm.

To functionally test our CellChat predictions, we first analyzed three highly ranked receptors by morpholino knockdown of *syndecan-1* (*Sdc1*) and pharmacological inhibition of Ephrin B and FGFR1. CellChat-inferred receptor *Syndecan-1* (*Sdc1*) and *Fn1*, *tenascin*, and *Col1a1* ligands were upregulated during blastema formation, as confirmed by HCR (**Supplementary Figure 19a),** finding the expression in the basal keratinocytes, the apical epidermal cap, and the *Prrx1^+^* CT cells (**Supplementary Figure 19b**). However, although Sdc-1 morpholino-treated axolotls exhibited delayed regeneration, a reverse-complement control morpholino limb produced a similar phenotype (**Supplementary Figure 19c-f**). This result may be attributed to a possible non-cell autonomous action of SDC1 or the lack of specificity of *Sdc1* knockdown. Additionally, pharmacological inhibition of Ephrin B signaling using ALW-II-41-27 resulted in 100% of animal mortality within 24 hours, precluding regeneration analysis (data not shown). These limitations prompted us to focus on alternative CellChat-predicted pathways.

Among the remaining candidates to functionally test our CellChat results, NCAM-FGFR1 signaling showed the strongest increase at 5 dpa among pathways already active in the uninjured limb, while FGFR1-FGF10 interactions emerged at 10 dpa (**Figure 4c**). HCR-ISH analysis confirmed the expression of both *Ncam* and *Fgf10* in *Prrx1*^+^ CT cells in the blastema (**Figure 4d-e**), identifying FGFR1 as a promising candidate for functional validation.

To test this, we treated axolotls with the FGFR1-specific inhibitor AZD4547 (via bath immersion) every other day for 14 days after amputation, corresponding to the late-bud blastema stage of development (**Figure 5a**). At 14 dpa, blastema formation was impaired in FGFR1-inhibited animals, achieving complete wound healing and the formation of a structure resembling the Apical Epidermal Cap (AEC), but blastema outgrowth was not observed (**Figure 5b**). By tracking regeneration of *Prrx1^Cre-ERT^Caggs^STOP/Cherry^* axolotls through 60 dpa, we confirmed that the 14-days FGFR1 inhibition window was sufficient to permanently abolish limb regeneration in all treated animals (**Figure 5c-d**), demonstrating that FGFR1 signaling during blastema formation is indispensable for limb regeneration.

**Figure 5.**
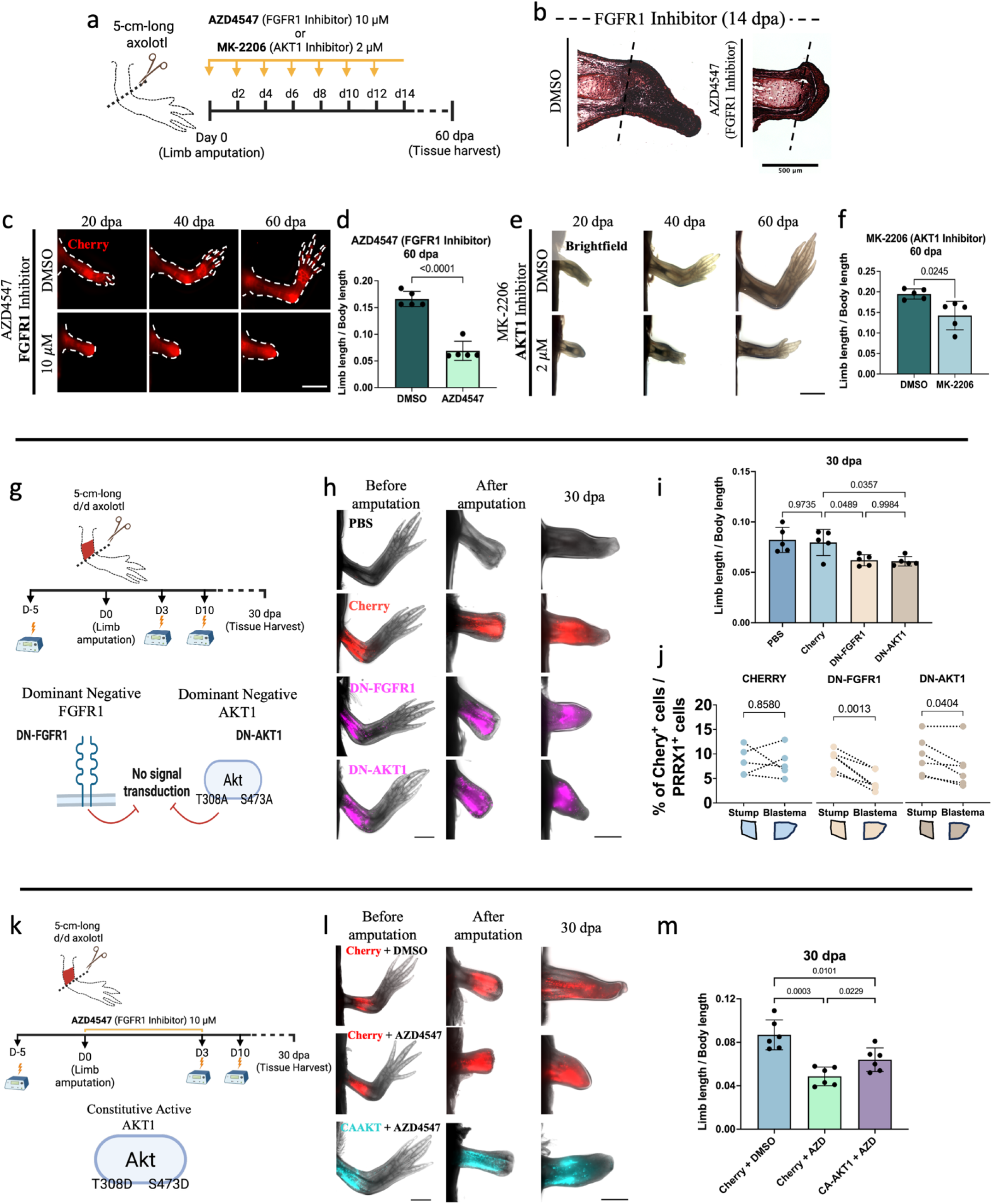
CellChat identifies FGFR1/AKT1 as a key CT-cell signaling axis required for blastema formation and limb regeneration. **a-f,** Pharmacological inhibition of FGFR1 and AKT1. **a,** Schematic of the experimental design for pharmacological inhibition studies using the FGFR1 inhibitor AZD4547 (10 µM) and the AKT inhibitor MK-2206 (2 µM). Each inhibitor was administered by bath immersion every other day from 0 to 14 dpa. **b**, Masson’s trichome-stained sections of control DMSO- and AZD4547-treated limbs (left and right, respectively) at 14 dpa. The dashed line indicates the amputation plane. Scale bar: 500 μm. **c, e**, Stereoscopic images tracking limb regeneration over 60 days in converted *Prrx1^Cre-ERT^;Caggs^STOPCherry^* axolotls treated with AZD4547 (c) or d/d axolotls treated with MK-2206 (e), with treatment in both (c) and (e) administered during the first 14 dpa. Scale bar, 2 mm. **d, f**, Quantitative assessment of limb-length/body-length ratio of the regenerated limb at 60 dpa after AZD4547 (d), or MK-2206 (f) treatment. Data are presented as mean ± SD (n=5, for each condition). Statistical significance was determined using an unpaired two-tailed t-test, with corresponding p-values annotated in the graph for each comparison. **g-j**, Delivery of Dominant Negative (DN) FGFR1 or AKT1 proteins during limb regeneration. **g,** Schematic of the experimental design for mosaic overexpression and the delivery of DN constructs. DN-FGFR1, DN-AKT1, or Cherry-backbone control plasmids were electroporated at the stylopod level five days prior to amputation, with repeated delivery at 3 and 10 dpa. Bottom panel illustrates the molecular features of each construct: DN-FGFR1 lacks the intracellular kinase domain, retaining only the extracellular and transmembrane domains; DN-AKT1 carries inactivating point mutations at the two key phosphorylation sites T308A and S473A. **h**, Stereoscopic images tracking limb regeneration over 30 days following electroporation of PBS, Cherry (*Caggs:Cherry-T2A-MCS*), DN-FGFR1 (*Caggs:Cherry-T2A-DNFGFR1*), or DN-AKT1 (*Caggs:Cherry-T2A-DNAKT1*) in d/d animals. Scale bar, 2 mm. **i**, Quantitative of limb-length/body-length ratio at 30 across all four electroporation conditions. Data are presented as mean ± SD (n=5, for each condition). Statistical significance was determined using a one-way ANOVA with Tukey’s post-hoc test, with corresponding p-values annotated on the graph for each comparison. **j**, Quantification of the percentage of Cherry^+^ cells among the total number of PRRX1^+^ CT cells in the blastema region and the source zone for Cherry-backbone control, DN-FGFR1, and DN-AKT1. Individual paired values are plotted with dotted lines connecting measurements from the same animal. Statistical significance was determined using a paired two-tailed t-test; p-values are indicated for each comparison. **k-m**, Rescue experiment by delivery of constitutively active (CA) AKT1 protein during FGFR1 pharmacological inhibition. **k,** Schematic of the experimental design for mosaic overexpression and the delivery of CA construct. CA-AKT1 or Cherry-backbone control plasmids were electroporated at the stylopod level five days prior to amputation, with repeated delivery at 3 and 10 dpa. In the same experiment, a pharmacological inhibitor AZD4547 was administered during the first three days after amputation. The bottom panel illustrates the molecular features of the CA-AKT1 construct, which carries activating point mutations at the two key phosphorylation sites T308D and S473D, mimicking the protein phosphorylation state. **l**, Stereoscopic images tracking limb regeneration over 30 days following electroporation of Cherry (*Caggs:Cherry-T2A-MCS*) + DMSO treatment, Cherry + AZD4547 10 µM, or CA-AKT1 (*Caggs:Cherry-T2A-CAAKT1*) + AZD4547 10 µM in d/d animals. Scale bar, 2 mm. **m**, Quantitative of limb-length/body-length ratio at 30 across all four electroporation conditions. Data are presented as mean ± SD (n=5, for each condition). Statistical significance was determined using a one-way ANOVA with Tukey’s post-hoc test, with corresponding p-values annotated on the graph for each comparison.

To determine whether FGFR1 inhibition reflected a disruption of early immune cell activation, we quantified Iba1^+^ macrophages in 2-dpa blastemas following AZD4547 treatment. No significant difference was found compared with the DMSO control condition (**Supplementary Figure 20a-c**), indicating that FGFR1 inhibition does not perturb initial macrophage recruitment. Instead, BrdU pulse-labeling at 14 dpa in axolotls treated with AZD4547 from 8 dpa onwards revealed a significant reduction in CT-cell proliferation within the blastema (**Supplementary Figure 20d-f**), demonstrating that FGFR1 signaling specifically supports CT-cell proliferation. FGFR1 transduces signals through three major downstream branches: the MAP/ERK, JAK/STAT, and PI3K/AKT/mTOR pathways (**Supplementary Figure 21a**) ^24^. To determine which of these pathways is active in blastema CT cells, we assessed pathway-specific downstream target genes by RT-qPCR and HCR in blastemas treated with AZD4547 from 8 to 14 dpa. The MAPK-ERK pathway target *Raf* was unaffected (with the MAPK-ERK pathway target *Braf* undetectable in CT cells) (**Supplementary Figure 21c**). Similarly, the JAK/STAT pathway targets *Egf* and *Vcam1* showed no significant reduction in expression compared to DMSO-treated controls and showed comparable spatial expression patterns between conditions (**Supplementary Figure 21d**). In contrast, the PI3K/AKT/mTOR target *Akt1* was significantly downregulated in AZD4547-treated blastemas, with HCR imaging confirming a marked reduction of *Akt1* transcript levels in *Prrx1^+^* CT cells (**Supplementary Figure 21e**). These findings suggest that FGFR1 preferentially activates the PI3K/AKT/mTOR axis in blastema CT cells, with AKT1 serving as a key intracellular effector.

To functionally validate the role of AKT1 during limb regeneration, we treated regenerating axolotls with the AKT1 inhibitor MK-2206 for the same length of time that we used in the FGFR1 inhibition studies. Fourteen days of AKT1 inhibition significantly impaired limb regeneration, with treated limbs showing a significantly reduced limb-length-to-body-length ratio from 20 dpa onwards (**Figure 5e**). Unlike FGFR1-inhibition, AKT1-inhibited axolotls retained the capacity to form a blastema and limbs that were fully patterned at 60 dpa, but significantly smaller than those of the controls (**Figure 5f**), suggesting that AKT1 is a key but partially redundant downstream mediator of FGFR1 signaling during regeneration.

To confirm these findings using a cell-autonomous approach and to address potential off-target effects of pharmacological inhibition, we designed and overexpressed dominant-negative (DN) constructs using mouse FGFR1 and AKT1, thereby creating a mosaic environment in which some cells express the wild-type proteins, and others express the DN versions. Sequence alignments between mouse (*Mm*) and axolotl (*Am*) FGFR1 and AKT1 proteins confirmed high overall conservation, including full preservation of the FGFR1 extracellular domain and the key AKT1 phosphorylation sites T308 and S473, validating the applicability of established mammalian DN strategies ^25,26^ to the axolotl system (**Supplementary Figure 22a-b**). DN-FGFR1 comprises a truncated receptor lacking intracellular kinase activity, acting as a decoy that sequesters ligands without transducing downstream signals ^26^. DN-AKT1 carries inactivating T308A and S473A substitutions and a myristoylation sequence that tethers the constitutively inactive kinase to the cellular membrane ^25^ (**Figure 5g**). Both constructs were cloned independently into a Cherry-expressing backbone (*Caggs:Cherry-T2A-MCS*) to enable tracking of transfected cells. DN plasmids or Cherry-backbone control were electroporated at the stylopod level 5 days prior to amputation, with repeated delivery at 3 dpa and 10 dpa. Both PBS- and Cherry-electroporated limbs progressed normally, reaching the digit-formation stage at 30 dpa, while DN-FGFR1 and DN-AKT1-electroporated limbs exhibited delayed regeneration, reaching the medium-to-late-bud blastema stage at 30 dpa (**Figure 5h-i**). Notably, Cherry expression was markedly reduced in the distal blastema of DN-FGFR1 and DN-AKT1 axolotls, suggesting that cells expressing the mutated proteins fail to migrate into the blastema region and accumulate near the amputation plane. To verify this, we sectioned 30 dpa limbs, stained the sections for PRRX1, identified double-positive cells (PRRX1^+^/Cherry^+^) (**Supplementary Figure 22d**). Analysis in the blastema and source zone confirmed that DN-FGFR and DN-AKT1-expressing CT cells were preferentially retained near the amputation plane and failed to accumulate in the regenerating tissue distal to it, with DN-FGFR1-expressing cells showing more pronounced retention than DN-AKT1-expressing cells. In contrast, Cherry control cells were distributed uniformly across both the stump and blastema regions (**Figure 5j)**.

To confirm that the activation of AKT1 is downstream of the FGFR1 signal pathway, we ought to rescue the ablated phenotype caused by the FGFR1 pharmacological inhibition by overexpressing a constitutively-active (CA) mouse AKT1 construct. CA-AKT1 carries two specific mutations, T308D and S473D, mimicking the protein’s phosphorylated state, and a myristoylation sequence that tethers the constitutively active kinase to the cellular membrane ^25^. CA-AKT1 or Cherry-backbone control were electroporated at the stylopod level 5 days prior to amputation, with repeated delivery at 3 dpa and 10 dpa. After amputation, FGFR1 pharmacological inhibitor was administered by bathing during the first three days after amputation (**Figure 5k**). At 30 dpa, Cherry-electroporated limbs with DMSO control treatment regenerated normally, reaching a digit formation stage, whereas Cherry-electroporated + AZD4547 FGFR1 inhibitor presented delayed regeneration by forming an early-bud stage blastema. Contrary, the CA-AKT1-electroporated limbs + AZD4547 FGFR inhibitor developed a longer blastema with mesenchymal condensations and digit formation (**Figure 5l-m**), confirming a rescued phenotype of FGF1R1 inhibition by CA-AKT1 delivery. Collectively, these data provide cell-autonomous evidence that corroborates that FGFR1/AKT1 is a cell-intrinsic molecular axis controlling CT-cell participation in limb regeneration.

## DISCUSSION

This study explores the role of CT cells in axolotl limb regeneration, demonstrating their importance in blastema formation and providing new insights into their cellular and molecular dynamics during this process. CT cells have been previously considered crucial for limb regeneration and remain a tissue of interest among regeneration biologists ^8,27–29^ due to their unique ability to encode positional and patterning information ^9,30,31^, as well as their prevalence in the blastema ^8^. Previous quantitative studies from 1986 ^32^ suggested that CT cells comprise up to ∼43% of blastema cells. Here, to analyze the dynamic blastema cellular composition, taking into account all major cell types, we applied a spatial transcriptomic approach showing that up to ∼75% of blastema cells are CT-derived, reaffirming their status as the major blastema cellular population. Moreover, our unique approach using Xenium technology deconstructs the blastema’s cellular identities across its developmental stages, revealing a sequential pattern of cellular participation. Macrophages and immune cells dominate the early stages after amputation, consistent with their established roles in wound healing and early regenerative signaling. While our selected probe set did not resolve specific immune cell subpopulations, recent work has shown that adaptive immune T and B lymphocytes are abundant at 1 and 2 days after amputation, despite their dispensability for limb regeneration ^13^. Other cell types - including muscle progenitors, endothelial cells, and neuronal components - contribute minimally to the early blastema and maintain low proportions throughout the late-bud blastema stage, suggesting that they repopulate the regenerating limb during subsequent stages. The apparent absence of neuronal components in our dataset is most likely due to Xenium’s limited resolution of thin axonal fibers that innervate the blastema, as innervation has been shown to be crucial for axolotl limb regeneration ^33,34^. Intriguingly, muscle progenitors showed limited presence in the blastema until 14 dpa, coinciding with the peak of CT-cell accumulation. This temporal overlap suggests a sequential pattern of cellular recruitment, from macrophages and immune cells to CT-derived progenitors, and then to muscle progenitors, raising the compelling hypothesis that each arriving population may provide signals to recruit the next.

Underscoring the central role of CT-derived cells in blastema development, we demonstrate that genetic ablation of CT cells severely impedes both limb development and regeneration. Interestingly, we find that ablation at early limb-regeneration stages leads to a less severe phenotype, whereas ablation of CT cells at later stages produces a more severe phenotype. A possible explanation is that partial ablation of CT cells in the early stages results in cellular compensation due to signals that sustain CT-cell dedifferentiation and proliferation, likely mediated by the low innervation detected, which provides crucial signals to maintain blastema growth. Similarly, other studies have focused on depleting specific cellular populations during limb regeneration. An important role for macrophages was also shown in a previous study in the axolotl, in which clodronate-induced macrophage depletion resulted in inhibition of axolotl limb regeneration ^10^. In contrast, muscle progenitor depletion in *Pax7-*knockout axolotls has shown that muscles play a limited role in limb regeneration, and even under severe deficiency of the muscle lineage, the axolotl limb develops and regenerates almost perfectly ^35,36^. Keratinocytes are not considered part of the blastema and were excluded from this study. However, it is known that keratinocyte-CT cellular crosstalk is essential for limb regeneration ^37^. In future studies, it will be important to ablate other cell types, including nerve cells, non-macrophage innate immune cells, and endothelial cells, both to understand the specific roles of these cells in limb regeneration and to gain a more comprehensive understanding of the cellular-crosstalk occurring with CT-cells, which allows them to induce, maintain, organize, and completely achieve axolotl limb regeneration.

CT-derived cells are primary contributors to the blastema by undergoing cell dedifferentiation and acquiring a state resembling mesenchymal embryonic limb bud cells ^8^. However, the signals, cellular pathways, and molecular changes that govern this cellular reprogramming are poorly understood. In this study, we characterized the molecular transitions of CT-derived cells during regeneration and found distinct waves of gene regulation. The earliest upregulated set of genes is associated with cellular chemotaxis, coinciding with the time of immune cell invasion, and the second wave peaking between 3 and 7 dpa involves genes that modulate the pH microenvironment. While the functional significance of this pH modulation in CT-derived cells remains unexplored, we hypothesize that it could be related to phagocytosis to enable extracellular-matrix remodeling at the stump tissue, and our findings establish a foundation for future investigations. Finally, at the late-bud blastema stage, CT cells exhibited activation of the differentiation program, characterized by embryonic limb development and morphogenesis, providing a defined temporal window to study the mechanisms controlling dedifferentiation and subsequent redifferentiation of CT-derived cells.

To accelerate research in regenerative biology, we have developed an open-access interactive web platform encompassing a comprehensive dataset of CT-derived cell expression dynamics during blastema development. This resource enables exploration of differentially regulated genes and pathways during the regenerative process, facilitating the discovery of novel mechanisms controlling limb regeneration.

Additionally, we introduced a versatile genetic ablation system for the axolotl model, featuring an inducible loxP-based NTR2.0 transgene compatible with any tissue-specific Cre-driver. This powerful tool enables precise temporal control of cell-type-specific ablation, allowing researchers to dissect the functional roles of distinct cell populations across various regenerative contexts. Two preliminary studies - a tBID-based conditional ablation system applied to CT cells ^38^ and a systematic comparison of NTR2.0 and inducible Caspase-9 (iCASP9) ^39^ - highlight the importance of several aspects of experimental design when using ablation systems in axolotls, including the efficiency of Cre-mediated recombination in the target tissue, prodrug delivery route, animal size, and the proliferative states of the target cell population. In the present work, NTR2.0/MTZ ablation was performed in 5- to 6-cm-long axolotls, and CT-cell ablation was confirmed by Caspase-3 immunostaining. Survival studies confirmed a high survival rate following overnight 20 mM MTZ treatment across a 10-day protocol. Most importantly, ablated phenotypes were consistent with the tBID-based approach ^38^, providing convergent evidence of the necessity of CT cells for limb regeneration, regardless of the ablation mechanism employed. The broad compatibility of our NTR2.0 system extends its applicability beyond limb regeneration, providing a valuable resource for investigating cellular mechanisms in both development and regeneration in other tissues and cellular populations in the axolotl ^40^.

Our identification of FGFR1/AKT1 signaling as a cell-autonomous requirement for CT-cell participation in blastema formation is consistent with previous work in model organisms. In zebrafish, early inhibition of FGFR1 blocks blastema formation without effects on wound healing ^41^ - a phenotype similar to what we observe in the axolotl following AZD4547 treatment. In previous work with the axolotl, broad FGFR inhibition with SU5402 during limb development produced smaller limbs with posterior digit loss ^42^, demonstrating a role for FGF signaling in CT-cell proliferative expansion. Similarly, FGF signaling has been shown to regulate SHH induction and subsequent patterning in the axolotl blastema ^43,44^. Our findings extend these previous findings in zebrafish and axolotl by suggesting that FGFR1 functions as an autocrine receptor specifically in CT cells, and that its inhibition impairs CT-cell proliferation and blastema formation without perturbing early macrophage recruitment. The complete and permanent abolition of limb regeneration that we observed following FGFR1 inhibition is one of the most severe pharmacological phenotypes reported in studies of axolotls, comparable in magnitude to the phenotypes resulting from TGF-β or Wnt/β-catenin inhibition during the first stages after limb amputation ^45,46^. This observation suggests that FGFR1 signaling in CT cells is a molecular requirement operating at the same hierarchical level as these two pathways with established roles during the first stages of limb regeneration.

Surprisingly, CellChat analysis inferred that the major ligand acting on the FGFR1 receptor was NCAM. A 1986 study in newts established that blockade of NCAM activity by the infusion of Fab fragments after limb amputation delays newt limb regeneration ^47^. We hypothesize that NCAM expressed in CT cells engages FGFR1 to sustain autocrine receptor activation, providing a molecular basis for the observation in this earlier functional study in newts and our experiments. Whether NCAM functions as a regulator of canonical FGFR1/FGF activation remains to be determined, as biochemical evidence indicates that NCAM acts as a competitive inhibitor of the interaction of FGF to FGFR receptors ^48^.

Our results showing selective activation of AKT1 and its downstream effector mTOR is consistent with previous work in axolotl identifying mTOR as a key upstream signal required for initial wound closure following limb amputation and for subsequent sustained limb regeneration ^49^. Our findings suggest that the FGFR1/AKT1 axis may represent a cell-type-specific upstream activator of the mTOR-dependent translational program essential for regeneration.

Together, our findings reframe the blastema’s cellular composition as an active, dynamic cellular community in which CT cells sustain their participation through cell-autonomous molecular programs. Furthermore, the tools and datasets introduced in this study establish a robust toolkit for future systematic and mechanistic investigations of regenerative processes and cellular plasticity.

## METHODS

### Axolotl husbandry

Axolotls (*Ambystoma mexicanum*) used in this study were bred and reared at the animal core facilities of the Mount Desert Island Biological Laboratory (MDIBL) in Maine, USA, and the Institute of Molecular Pathology (IMP) in Vienna, Austria. All animal handling and surgical procedures adhered to local ethics committee guidelines approved by the Institutional Animal Care and Use Committee (IACUC) of MDIBL and the Magistrate of Vienna. The experiments using animals and the generation of the transgenic axolotl line, were conducted using the axolotl *d/d* strain, characterized by leucistic skin resulting from a defect in melanocyte migration during embryonic development that is associated with a mutation in the *Edn3* gene ^50^.

Axolotl larvae were bred using 20% Holtfreter’s solution prepared with locally sourced tap water from Maine or Vienna. Experiments were conducted using axolotls ranging in length from 5 to 6 cm from snout to tail, corresponding to approximately 4-month-old post-egg laying. To prevent cannibalism, axolotls were individually housed in containers from the age of 1-month post-hatching onwards.

Axolotls were fed daily *ad libitum* with Artemia (*Artemia gracillis*) for the first 3 months after hatching, after which their diet was switched to 5 mm food pellets (Rangen, Inc.) once they reached a length of 4 cm. Complete water changes were performed three times each week to maintain water quality. Axolotl husbandry practices remained consistent throughout all experimental procedures.

### Axolotl transgenesis and DNA recombination induced with tamoxifen

The *Prrx1*^+^ Cre driver (tgSceI(Mmu.*Prrx1*:TFPnls-T2A-Cre-ER^T2^)^Etnka^) (hereafter, *Prrx1^Cre-ERT^*), and LoxP reporter tgSceI(*CAGGs*:LoxP-GFP-dead(STOP)-LoxP-Cherry)^Etnka^, (hereafter, *Caggs^STOP/Cherry^*) were previously reported ^8,31^. To generate LoxP driver transgenic axolotls capable of inducing expression of the nitroreductase enzyme (tgSceI(*CAGGs*:LoxP-BFPnls-LoxP-Cherry-T2A-NTR2.0)^Pmx^) (hereafter, *Caggs^nBFP/Cherry-NTR2^*), a *T2A* sequence followed by the *NfsB* gene sequence was cloned at the 3’ end of the *CAGGs:LoxP-BFPnls-LoxP-Cherry* cassette, flanked by I-SceI sites. The *NfsB* sequence was obtained from the plasmid 5xUAS:GAP-mCherry-P2A-NfsB_Vv F70A/F108Y;he:tagBFP2, which was a gift from Jeff Mumm (Addgene, 158653) ^21^. The resulting plasmid was co-injected along with I-SceI endonuclease into one-cell stage embryos as previously described ^51^. Larvae exhibiting nBFP fluorescence were screened and raised for one-and-a-half years until sexual maturity was reached; these animals served as founder breeders. Successful germ-line transmission was confirmed by observing constitutive nuclear BFP expression in the offspring larvae. The construct used for generating Lp-NTR2.0 axolotls has been deposited in (Addgene, 219785).

Cre-mediated recombination was induced with 4-hydroxytamoxifen (4-OHT) treatments. *Prrx1^Cre-ERT^;Caggs^STOP/Cherry^* and *Prrx1^Cre-ERT^;Caggs^nBFP/Cherry-NTR2^* larvae at stage 44 ^52^ were treated with 2µM 4-OHT (Sigma-Aldrich, H6278) three times on alternate days. Each treatment involved bathing the larvae for 6 hours in the dark at room temperature (RT). Ten days after the first 4-OHT treatment, converted axolotls (Cherry^+^) were screened and allowed to grow up to 5-6 cm in length for experiments. For limb-development studies, larvae were screened and used immediately at stage 47 ^53^.

### Limb amputation

Axolotls were anesthetized using 0.03% benzocaine (Sigma-Aldrich, E1501) diluted in 20% Holtfreter’s solution. Once the axolotls no longer responded to mechanical stimuli, they were transferred to a surgical plate under a stereoscope (Carl Zeiss Microscopy, Stemi DV4). The right forelimb was then amputated at the mid-zeugopod level, and any protruding bone was carefully trimmed using scissors to facilitate skin retraction and wound healing. Subsequently, axolotls were immediately transferred to fresh 20% Holtfreter’s solution to aid their recovery.

### Limb harvesting, histological processing, and immunofluorescence

Complete right forelimbs were harvested at various time points and fixed overnight (O/N) in 4% paraformaldehyde (Sigma-Aldrich, P6148) at 4°C with agitation. Subsequently, the samples were washed three times with PBS for 5 minutes each at RT and transferred to 30% sucrose (Sigma-Aldrich, S0389) diluted in PBS at 4°C with agitation until they sank. The samples were embedded in an O.C.T. Compound (Fisher Scientific, 23-730-571) and stored at -70°C. Sagittal and longitudinal cryosections of 20 µm of thickness were obtained for immunofluorescent protein detection.

For immunofluorescence staining, sections were washed three times with PBS for 5 minutes each RT, followed by permeabilization with 0.3% Tween-20 (Sigma-Aldrich, P1379) in PBS for 40 min at RT. Subsequently, the sections were blocked with PBS containing 0.3% Triton-X-100 (Millipore, T8787) and 2% Normal Horse Serum (Vector Laboratories, S-2000) for 1 hour at RT. Slides were then incubated O/N with the primary antibody. The primary antibodies used in this study were rabbit anti-PRRX1 (IMP, homemade, ratio 1:200), mouse anti-Caspase 3 active (Abcam, Ab13847, ratio 1:500), mouse anti-BrdU (DSHB, G3G4, concentration 5 μg/ml), and rabbit anti-Iba1 (Wako Chemicals, 019-19741, ratio 1:500) diluted in blocking buffer.

The following day, the primary antibody was washed thrice with PBS for 5 minutes each at RT. Subsequently, the sections were incubated with the secondary antibodies. We used goat anti-rabbit coupled with Alexa Fluor 647 fluorophore (Invitrogen, A21245) or donkey anti-mouse diluted coupled with Alexa Fluor 647 fluorophore (Jackson, 715607003) at a ratio of 1:250 in PBS for 3 hours at RT. Slides were washed thrice with PBS for 5 min each at RT and stained with DAPI (Sigma-Aldrich, 10236276001) at a concentration of 0.5 µg/µl for 15 min. Finally, slides were mounted with Vectashield (Vector Laboratories, H-1900-10) for subsequent microscopy visualization.

For BrdU detection in limb sections, we treated the sections with 2 M HCl for 20 min at 37°C to denature DNA before the blocking step, as previously described ^54^.

### Lineage-tracing experiments and cell counting in blastemas

We previously demonstrated that the *Prrx1^Cre-ERT^* driver and anti-PRRX1 antibodies both reliably label CT cells ^8^. However, labeling through the Cre driver is not absolute. Similarly, while anti-PRRX1 antibodies produce a strong signal in the blastema, they do not detect CT cells well in the mature limb, particularly those embedded within muscle fibers. To address these limitations, we used a combined approach: lineage tracing with *Prrx1^Cre-ERT^;Caggs^STOP/Cherry^* animals and immunohistochemistry with anti-PRRX1 antibodies to identify CT cells across the mature limb, blastema, and stump regions. For this assay, cells positive for Cherry, anti-PRRX1, or both were classified as CT-derived cells.

Converted *Prrx1^Cre-ERT^;Caggs^STOP/Cherry^* axolotls ranging from 5 to 7 cm in length, were used in these experiments and randomly assigned to each experimental condition. Right forelimbs were amputated, and complete limbs were harvested at 3, 5, 7, 10, and 14 days post-amputation (dpa). Longitudinal cryosections were obtained, and immunofluorescence staining for PRRX1 was conducted. Mosaic images of the sections were captured using a Zeiss Axio Observer.Z1 microscope equipped with a 20X apochromatic lens (0.8 Numerical Aperture (NA)) and Axiovision or Zen2 software (Zeiss). The resulting files were saved in .zvi format and analyzed using the Cell Counter plugin in FIJI software. The counted cells were categorized into CT-derived cells in the source zone, non-CT-derived cells in the source zone, CT-derived cells in the blastema, and non-CT-derived cells in the blastema. The “source zone” was defined as the 500 µm preceding the amputation plane. Notably, the epidermis and apical epidermal cap were excluded from the analysis.

For each time point, three sections (technical replicate) were analyzed for each biological replicate. Statistical analysis was performed using PRISM (version 10.2.0). Student’s t-tests were performed at each time point, comparing non-CT cells with CT cells and comparing each group of cells at every time point to the uninjured condition.

### Specific cell ablation with metronidazole treatments and image acquisition

Converted *Prrx1^Cre-ERT^;Caggs^nBFP/Cherry-NTR2^*axolotls, measuring from 5 to 6 cm in length, were used in these experiments and randomly assigned to each experimental condition. The right forelimbs were amputated, and regeneration was monitored before amputation and every 10 days until the completion of regeneration at 60 dpa. Metronidazole (MTZ) treatments were administered at 5, 7, 10, and 14 dpa, as required. Metronidazole (Millipore Sigma, M3761) was diluted in 20 % Holtfreter’s solution at 10 mM for larvae experiments and 20 mM for 5-to-6-cm axolotl experiments as a final concentration. A single MTZ treatment was performed by overnight bathing for approximately 18 hours in the dark at RT. Larvae were treated together in 1L solution, whereas 5-to-6-cm-long axolotls were treated individually in containers with 200 mL each. The following day, axolotls were transferred to 20% of Holtfreter’s solution.

At each time point analyzed, axolotls were anesthetized with 0.03% benzocaine diluted in 20% Holtfreter’s solution until unresponsive to mechanical stimuli. Subsequently, they were transferred to a surgical plate where the body length and stretched length of the amputated and non-injured contralateral forelimbs were measured with a ruler underneath. Following measurement, axolotls were transferred to a petri dish containing 1% solidified agarose (Sigma-Aldrich, A9539) diluted with PBS to serve as a background for imaging.

Images were acquired using a Zeiss Axio Zoom.V16 (Carl Zeiss Microscopy, 435080-9031-000) equipped with a Zeiss Plan Z 1.0x/0.25 objective lens (Carl Zeiss Microscopy, 435282-9100-000). Samples were illuminated with a 120W Metal halide lamp (Carl Zeiss Microscopy, HXP 120C). BFPnls fluorescence was excited and collected by using a filter set 01filter cube (Carl Zeiss Microscopy, 488001-9901) consisting of an excitation filter: Zeiss BP 365/12 nm + dichromic mirror Zeiss FT 395 nm + emission filter: Zeiss LP 397 nm. mCherry fluorescence was excited and collected by using filter set 63 HE (Carl Zeiss Microscopy, 486093-0000) consisting of an excitation filter Zeiss BP 572/25 nm (HE) + dichromic mirror Zeiss FT 590 nm (HE) + emission filter: Zeiss BP 629/62 nm (HE). Z-stacks images were acquired with a Zeiss Axiocam 506 color camera (Carl Zeiss Microscopy, 426556-0000) controlled with a Zen 3.1 (Carl Zeiss Microscopy) at zoom 15x, binning 1x1, at a resolution of 2752 x 2208 pixels, in 14 bit and saved in CZI format. Maximal z-stack projections and automatic intensity values of every acquired fluorophore were obtained using FIJI software. Background removal was performed manually in some images without altering limbs. Scale bars were added to each image using FIJI software, and the metadata was read for size dimensionality.

Statistical analysis for limb lengths was done using PRISM software (version 10.2.0). Two-tailed Student’s t-tests were performed for each time point, comparing non-MTZ-treated conditions with MTZ treatments. A significance level of 0.05 was used for all tests, with each p-value annotated accordingly in the graphs.

### Microarray design for PRRX1^+^-CT cells

A custom Agilent 400K microarray (Agilent Technologies) was designed using the axolotl transcriptome assembly release 25 (https://www.axolotl-omics.org/assemblies, version Am_2.2). Three sense probes and one antisense probe were designed for each annotated transcript with known strandedness. For transcripts with unknown strandedness, which includes non-coding, partial, or novel sequences lacking homology to known proteins, three sense, and three antisense probes were generated, utilizing RNA-seq strandedness data to guide orientation. Additionally, 20,839 probes from a previously established EST and cDNA-based microarray ^55^ were incorporated to ensure backward compatibility. Overall, the microarray used in this project comprises a total of 415,996 probes.

### Tissue collection for microarray and library preparation

Converted *Prrx1^Cre-ERT^;Caggs^STOP/Cherry^* axolotls were amputated at the mid-zeugopod level at the size of 4.5-5 cm from nose to cloaca (corresponding to 9-10 cm from nose to tail), ignoring their sex. Tissue from the intact lower arm or from the blastema at different stages of regeneration (1, 3, 5, 7, 10, and 14 dpa), including tissue 2 mm behind the amputation plane, was collected and washed in 0.8x PBS buffer. Blastemas from 4-6 axolotls from the same clutch were pooled to collect a sufficient number of cells. Three biological replicates represent independently collected and processed samples from three separate animal batches.

Tissue was transferred into 1 ml of dissociation solution containing 0.35/ml of Liberase TM (Roche, 05401119001) and 100 U/ml of DNase1 (Roche, 04716728001) in 0.8x PBS. Tissue was dissected into small pieces using forceps and then rocked or pipetted for approximately 30 minutes at RT. The cell suspension was cooled on ice and filtered through a 70-µm filter before FACS-sorting. Cherry^+^ cells were sorted directly into the lysis buffer RLT (Qiagen, 79216). Total RNA from 150,000 to 250,000 cells per sample was purified using RNeasy mini kit (Qiagen, 74104) according to the manufacturer’s protocol.

Cy3-labeled cRNA probes were generated from 45 ng of purified total RNA using Agilent Low Input Quick Amp Gene Expression Labeling Kit (Agilent, 5190-2307). Custom-designed 400K Agilent arrays (Name:Am400k_v2, Design ID: 084163, Design Format: 2x 400 K, Control Grid: IS-420288-2-v2_400kBY2_gx_eqc_20100210) were probed with 3.75 ug of cRNA per array according to the Agilent protocol and scanned using the AgilentG3_GX_1Color Raw scan protocol.

Raw Agilent microarray data were imported and processed using limma v3.58.1 in R v4.3.2, applying background correction using the normexp method, quantile between array normalization, filtering out control and low-expressed probes, as well as probes assigned to multiple genes, and averaging intensities of multiple probes per gene. Differential expression at an absolute log2Fold Change of 1 with an adjusted p-value < 0.01 of d1 – d14 timepoints versus d0 was determined using limma-trend and functions lmfit, treat and topTreat. Gene set over-representation analysis per cluster was performed with clusterprofile v4.10.01 and GO.db v3.18.0.

### Axolotl transcriptome assembly for microarray and pseudobulk analysis

For microarray and pseudobulk analysis, the axolotl transcriptome assembly was built by improving AMEX_v47 using private RNAseq data graciously shared by Dr. James Godwin. Axolotl genome version AMEX_v6.0-DD and associated transcriptome assembly AMEXT_v47 were downloaded from axolotl-omics.org. RNA-seq fastq files (from Dr. Godwin) were analyzed with the Nextflow (https://nextflow.io/) NF-core (https://nf-co.re/) rnaseq workflow, version 3.8.1 (https://nf-co.re/rnaseq/3.8.1), using the Amazon Genomics Command-line (AGC) interface, version 1.5. The analysis was carried out in a two-stages. In the first stage, the sequences were run through the pipeline with default options except for the following: “--skip_bbsplit --remove_ribo_rna --igenomes_ignore --save_reference --stringtie_ignore_gtf -- pseudo_aligner salmon --aligner star_salmon”. The sequences were aligned to the draft genome with STAR ^56^ version 2.6.1d. With the –stringtie_ignore_gtf flag set, each sample file’s alignments were passed through stringtie version 2.1.7 to create a sample-specific GTF file describing the inferred transcript structures. In the second stage, stringtie ^57^ version 2.1.4 was run in “--merge” mode, in which all sample-specific GTF files were merged with the GTF from the draft genome to create an updated GTF that included evidence of the current data files. The resulting annotation GTF and transcriptome fasta-formatted sequence files were used as targets for all subsequent analyses described here. GTF (**Supplementary Data 4**) file is included as supplementary material.

### Comparison between microarray analysis and single-cell data

Single-cell data of sorted PRRX1^+^ cells and unsorted cells from different time points of limb regeneration were obtained from previously generated data ^8^. Raw reads were processed using nf-core/scrnaseq (v2.3.0) ^58^ from the nf-core collection of workflows ^59^. Reads were aligned to the axolotl transcriptome using STAR ^56^ as implemented in 10x Genomics Cell Ranger (v7.1.0). We used R studio to run custom R scripts for analysis. Genes expressed in fewer than three cells were excluded from the analysis. Cell quality control was conducted on the data by eliminating the bottom and top 10% of cells based on read count, along with cells containing a mitochondrial percentage greater than 10%. The *Seurat* package (v4.1.3) ^60^ was used for clustering cell identity classification and pseudo-bulk generation. Expression levels were normalized through the “NormalizedData” function with the LogNormalize method. For visualization, the “ScaleData” function was used to regress out differences in the number of molecules, number of genes, percent mitochondrial genes, cell cycle effects, and the expression of fluorescent labeling proteins (eGFP and mCherry). Principal component analysis (PCA) was conducted, and the first 31 PCs were used to identify clusters and generate both t-distributed stochastic neighbor embedding (t-SNE) and uniform manifold approximation and projection (UMAP) plots. To extract exclusively connective tissue cells, the dataset was reduced by excluding clusters containing only cells from the unsorted samples. The pseudobulk counts were then generated using the “AgregateExpression” function grouping by sample.

The pseudobulk counts were passed into the RNfuzzy web application ^61^, and low-count genes were filtered out using a threshold of 10. Differential expression analysis was then conducted using the edgeR analysis method, the TMM normalization method, and a 0.001 FRD cutoff.

Normalized, filtered, and replicate-averaged microarray expression data were converted to counts and analyzed for differential expression using BioTEA software (v.1.1.0), comparing 5 dpa vs uninjured condition and 10 dpa vs. uninjured condition. Each expression dataset was then independently merged with a probe-to-gene table to label each probe with its corresponding axolotl gene annotation. Probes aligning with multiple genes were excluded, and gene-level transcript counts were tabulated. Mean-per-gene expression, fold-change, and adjusted P-values were calculated to account for cases of multiple probes aligning with a given name.

### 10X Xenium spatial imaging

Xenium panel design, limb histology, and tissue processing were performed with the manufacturer’s instructions (10X Genomics, Xenium in Situ). The list of all transcripts detected can be found in **Supplementary Data 1**. The custom panel design targeted 100 genes relevant to limb regeneration and cell type classification. Each gene was given between 6 and 8 probes and was developed based on the provided gene sequence. The final genes and number of probes per gene selected were validated by cross-referencing with the unfiltered single-cell expression data. This ensured adequate cell type utilization, specifically preventing overcrowding of any singular cell type, which could lead to an inability to detect distinct fluorescent signals. A list of all probes used is provided as **Supplementary Data 3**.

Xenium slides were prepared following manufacturer’s instructions from fresh frozen tissue sections (Protocol: CG000581) followed by probe hybridization, ligation, and amplification (Protocol: CG000582). Slides were run on a Xenium Analyzer instrument running Xenium software (v.2.0.1.0) and Onboard Analysis software (v.2.0.0.10) to produce the output data bundle used for downstream analysis. Following the Xenium run, slides were H&E stained in a Sakura Tissue-Tek Prism stainer, and whole slide imaging was conducted at 40x magnification using an Aperio GT450 instrument (Leica).

Secondary analysis of Xenium experiments was performed by the Xenium Onboard Analysis software (v2.0.0.10). The analysis includes PCA, K-means and graph-based clustering, differential expression, and UMAP dimensionality reduction.

### Comparison between microarray analysis and Xenium data

Xenium data was post-processed using the Seurat ^62^ package (v.5.1.0) to facilitate streamlined data manipulation. Microarray expression data was reduced to include only probes with maximum match scores that targeted the positive strand of corresponding transcripts. This subset was further refined to include only probes that matched those in the Xenium experiment, followed by the averaging of biological replicates. Each Xenium sample was annotated using graph-based clustering algorithms, and subsequent analyses were focused on proposed blastema clusters. Differential expression analysis was performed independently for each time point (Day 3, 5, 7, 10, and 14) compared to Day 0. Z-score normalization was applied to each probe’s temporal expression profile to eliminate scaling inconsistencies between datasets. Probes were evaluated for cross-experimental correlation over time using Pearson’s correlation coefficients to assess consistency between technologies.

### Inference of cellular communication (CellChat)

Intercellular communication was inferred from single-cell transcriptomic data using the CellChat package ^16^ (v2.2.0). As no axolotl-specific CellChat database is currently available, we constructed a custom axolotl signaling database by mapping axolotl gene identifiers onto the human CellChatDB reference database. Axolotl feature annotations were preprocessed to extract canonical gene symbols by removing species-specific suffixes, locus identifiers, and isoform notation. Human ligand and receptor gene symbols in the interaction table were systematically replaced with their corresponding axolotl identifiers, and multi-subunit protein complexes were renumbered to reflect the species-specific gene-symbol mappings. To minimize the risk of falling to detect biologically relevant interactions, non-one-to-one mappings were retained in the translated database by duplicating interaction entries for each axolotl gene mapped to a single human gene. Normalized counts were extracted from the preprocessed single-cell dataset, and intercellular communication was inferred independently for each condition across all annotated cell types. Cross-condition comparisons were performed by merging individual CellChat objects. All visualizations and downstream analysis were performed using CellChat and ggplot2 functions.

### Pharmacological treatments

All pharmacological treatments were administered by bath immersion. Inhibitors were diluted from stock solutions to the indicated working concentrations in 20% Holtfreter’s solution. Each axolotl was treated individually in 150 mL of solution in a separate container, and the treatment solution was replaced every other day throughout the duration of each experiment. The following inhibitors were used: the FGFR1 inhibitor AZD4547 (Fexagratinib, MedChemExpress, HY-13330, 10 μM and 20 μM working concentrations, stock 50 mM in DMSO), the AKT1 inhibitor MK-2206 (MedChemExpress, HY-10832, 2 μM working concentration, stock 10 mM in DMSO), and the EphrinB2 inhibitor ALW-11-41-27 (MedChemExpress, HY-18007, stock 50 mM in DMSO). All stock solutions were aliquoted and stored at -20 °C. DMSO at the equivalent dilution was used as a vehicle control in all experiments.

### Hybridization Chain Reaction (HCR) and image analysis

HCR probes targeting axolotl *Prrx1, Fgfr1, Ncam, Fgf10, Sdc1,Col1a1, Raf, Egf,* and *Akt1* were designed against sequences from the axolotl transcriptome assembly Amex.T_v47 ^63^, using a custom probe-generator tool developed at MDI Biological Laboratory. Each probe set consisted of a pool of 37 split-initiator probe pairs targeting multiple sites across the mRNA sequence (5’UTR, ORF, and 3’UTR) for use with the distinct HCR amplifier system ^64^. All probe sequences are provided in **Supplementary Data 5.** Lyophilized probes (Integrated DNA Technologies) were resuspended in nuclease-free water to a stock concentration of 100 mM. Single-, double-, or triple-plex HCR was performed on 50-um cryosections following the manufacturer’s instructions (Molecular Instruments). Fluorescence images were acquired using a Nikon Ti-E inverted confocal microscope coupled with a Yokogawa CSU-W1 25µm/50 µm spinning disk unit equipped with a Nikon CFI Plan Apochromat Lambda D 10X/0.45 air objective, a Nikon Plan Apochromat Lambda 20X/0.75 air objective, and a Nikon CFI Apochromat Lambda S 60X/1.40 oil objective. Samples were illuminated with a LUNF-XL laser unit combiner (405 nm - 50 mW, 448 nm - 60 mW, 561 nm - 40 mW, 594 nm - 40 mW, and 640 nm - 40 mW). DAPI fluorescence was excited and collected using a DM 405/488/561/640 dichromic beam splitter (Nikon, MHE46420) + Chroma ET 436/20X filter. GFP and Alexa 488 fluorescence were excited and collected using a filter Semrock FF01-525/60 filter. mCherry, and Alexa 541fluorescence were excited and collected using a Chroma ET605/52m bandpass filter. Alexa 647 was excited and collected using a Chroma ET705/72m single-band filter. Z-stack images were acquired with a CMOS Zyla 4.2 camera, binning 1x1, at a resolution of 2048 x 2048 pixels, in 16-bit and saved in ND2 format. Images were later denoised using Nikon NIS-Elements software. Maximal z-stack projections were obtained using FIJI software. Dot detection and quantification were performed using the RS-FISH plugin ^65^. Dots’ statistics were exported, and data visualization was performed using ggplot2 functions.

### RNA Extraction, cDNA synthesis, and RT-qPCR

Tissues from three regenerating limbs were pooled per biological replicate, snap-frozen in liquid nitrogen, and stored at -80 °C until processing. Total RNA was extracted using the NucleoSpin RNA kit (Macherey-Nagel, 740955) according to the manufacturer’s instructions. RNA integrity was assessed via agarose gel electrophoresis. First-strand cDNA synthesis was performed from 1 ug of total RNA using the M-MLV reverse transcriptase (Promega, M170A) following the manufacturer’s instructions. The resulting cDNA was quantified and diluted to a concentration of 0.5 μg/μl. Quantitative PCR was performed using SYBR Green PCR Master Mix (Thermo Fisher Scientific, 439155) with 1 ng of cDNA per reaction, following the manufacturer’s instructions. Each reaction was run in technical duplicate. Gene-expression levels were normalized to the *ribosomal protein 13* (*Rpl13*) reference gene, and relative mRNA abundance was calculated using the 2^−ΔΔCq^ method. Statistical significance between groups was assessed using an unpaired two-tailed Student’s t-test, with the DMSO-treated condition as the reference. All oligonucleotide primer sequences are provided in **Supplementary Data 6.**

### Dominant-negative DN-FGFR1, DN-AKT1, and constitutive active (CA) CA-AKT1 plasmid construction

Dominant-negative (DN) and constitutive active (CA) sequences were cloned into a Cherry-expression backbone plasmid (*CAGGS: Cherry_NLS_-T2A-MCS*), linearized by digestion with MluI (New England BioLabs, R3198) and NotI (New England BioLabs, R3189), followed by column purification using a PCR purification kit (Qiagen, 28106). A mouse DN-FGFR1 insert was amplified from plasmid pLenti-CMV-Blast-DNFGFR1-HA, which was a gift from Gian-Paolo Dotto (Addgene, Plasmid #130887) ^66^, which encodes a truncated FGFR1 comprising the extracellular and transmembrane domains but lacking the intracellular kinase domain. A mouse DN-AKT1 insert was amplified from plasmid pBSFI-myr-akt-T308A/S473A, which was a gift from Peter Vogt (Addgene, Plasmid #49191) ^67^, which encodes a myristoylated, constitutively membrane-tethered AKT1 carrying inactivating T308A and S473A mutations at both phosphorylation sites. A mouse CA-AKT1 insert was amplified from plasmid pBSFI-myr-akt-T308D/S473D, which was a gift from Peter Vogt (Addgene, Plasmid #49192) ^67^, which encodes a myristoylated, constitutively membrane-tethered AKT1 carrying activating T308D and S473D mutations at both phosphorylation sites. DN-FGFR1 (1078 bp), DN-AKT1 (1509 bp), and CA- AKT1 (1509 bp) amplicons were gel-purified using a gel extraction kit (Qiagen, 28706). Inserts were assembled into the linearized vector using Gibson assembly (New England BioLabs) according to the manufacturer’s instructions. All resulting constructs were verified by whole-plasmid sequencing (Plasmidsaurus Inc.). Oligonucleotide primer sequences used for cloning are provided in **Supplementary Data 6**.

### Morpholino and plasmid limb electroporation

*Syndecan-1* morpholino (5’ CTGGAAGTTAAGAAAGGCTGCTGCC 3’) and a control morpholino comprising the reverse complement sequence (5’ CCGTCGTCGGAAAGAATTGAAGGTC 3’) were synthesized by GeneTools LLC. Both morpholinos were synthesized with a 3’-Carboxifluorescein modification to enable visualization of transfected cells. Morpholinos were reconstituted to a stock concentration of 1 mM in 2X PBS and diluted to a working concentration of 500 µM in 1X PBS prior to electroporation.

Morpholino oligonucleotides and DN plasmid constructs were delivered into the limb as previously described ^68^. Briefly, the right stylopod or blastema was injected with 0.5 - 1.0 μl solution containing PBS, morpholino or DN constructs supplemented with Fast Green dye (0.1 %) to visualize injection, using a pneumatic air microinjector (World Precision Instruments, PV830). Immediately after injection, axolotls were immersed in cold 1X PBS and electroporated using a NepaGene electroporator (NEPA21, type II) equipped with 5-mm-diameter electrodes (NepaGene, CUY650P5). The poring pulse parameters consisted of three pulses at 150 volts with a pulse length of 5 milliseconds, a pulse interval of 10 milliseconds, a decay rate of 0%, and positive (+) polarity. The transfer pulse parameters were 5 pulses at 50 volts with a pulse length of 50 milliseconds, a pulse interval of 950 milliseconds, a decay rate of 0%, and a positive (+) polarity. For DN constructs, bidirectional polarity (±) was applied during the transfer pulses to maximize transfection efficiency. After electroporation, axolotls were transferred to fresh 20% Holtfreter’s solution for recovery.

## DATA AVAILABILITY

The 10X Xenium spatial transcriptomic dataset and the microarray reported in this paper are deposited in the Gene Expression Omnibus (GEO) (Superseries data accession number: GSE294346 [under review process]). GTF file used for microarray and pseudobulk analysis is included as **Supplementary Data 4**.

## ACKNOWLEDGMENTS

We acknowledge the imaging support provided by MDIBL’s Light Microscopy Facility (LFM: RRID:SCR_019166) and the microscopy facility at IMP. We also thank the invaluable support of the animal core facilities at MDIBL and IMP and Dr. Stephen Sampson for proofreading this manuscript.

This research was funded by grants from the National Institute of Health (NIH) ORIP-R21OD031971, NIGMS-R35GM161492, and the DFG (527098031) awarded to PM. Research reported in this publication was supported by an Institutional Development Award (IDeA) from the National Institute of General Medical Sciences of the National Institutes of Health under grant numbers P20GM103423 and P20GM104318 to MDIBL. The 10X Xenium data acquisition was carried out in the Genomics and Molecular Biology Shared Resource (RRID: SCR_021293) at Dartmouth, which is supported by NCI Cancer Center Support Grant 5P30CA023108 and NIGMS COBRE P20GM130454 awards.

## AUTHOR CONTRIBUTIONS

Conceptualization: D.G.G., D.K. & P.M.

Methodology: D.G.G., D.K., M.K., K.W., H.F., R.P.S., R.E.G., S.N., K.E., M.N., F.W.K., J.H.G. & P.M.

Investigation: D.G.G., D.K., H.F., R.P.S., R.E.G., M.N., J.H.G. & P.M.

Writing original draft: D.G.G. & P.M.

Review and editing of the original draft: D.G.G., D.K., M.K., K.W., H.F., R.P.S., R.E.G., S.N., K.E., M.N., F.W.K., J.H.G. & P.M.

