## Supplementary Figures 1-22 for "The essential role of connective-tissue cells during axolotl limb regeneration"

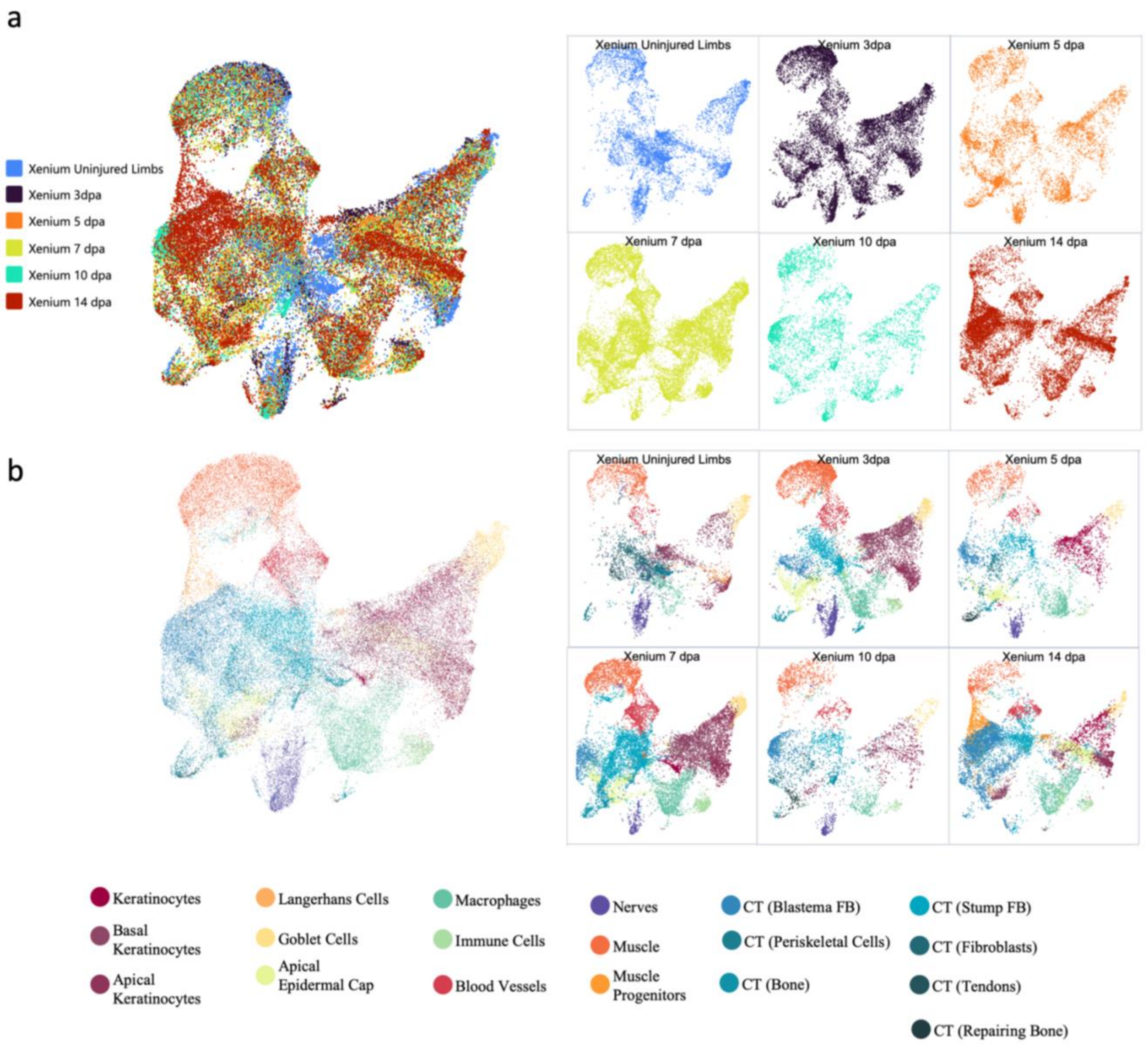
a
Xenium Uninjured Limbs
Xenium 3dpa
Xenium 5 dpa
Xenium 7 dpa
Xenium 10 dpa
Xenium 14 dpa
b
Xenium Uninjured Limbs
Xenium 3dpa
Xenium 5 dpa
Xenium 7 dpa
Xenium 10 dpa
Xenium 14 dpa
Keratinocytes
Basal Keratinocytes
Apical Keratinocytes
Langerhans Cells
Goblet Cells
Apical Epidermal Cap
Macrophages
Immune Cells
Blood Vessels
Nerves
Muscle
Muscle Progenitors
CT (Blastema FB)
CT (Periskeletal Cells)
CT (Bone)
CT (Stump FB)
CT (Fibroblasts)
CT (Tendons)
CT (Repairing Bone)

**Supplementary Figure 1. Detection of cellular identity of axolotl forearm regeneration using spatial transcriptomics**. **a**, Uniform manifold approximation and projection of all Xenium cells detected at all time points analyzed during blastema formation (left panel), and split by time point (right panel). **b**, Cell cluster detection and their distribution in all Xenium time points analyzed.

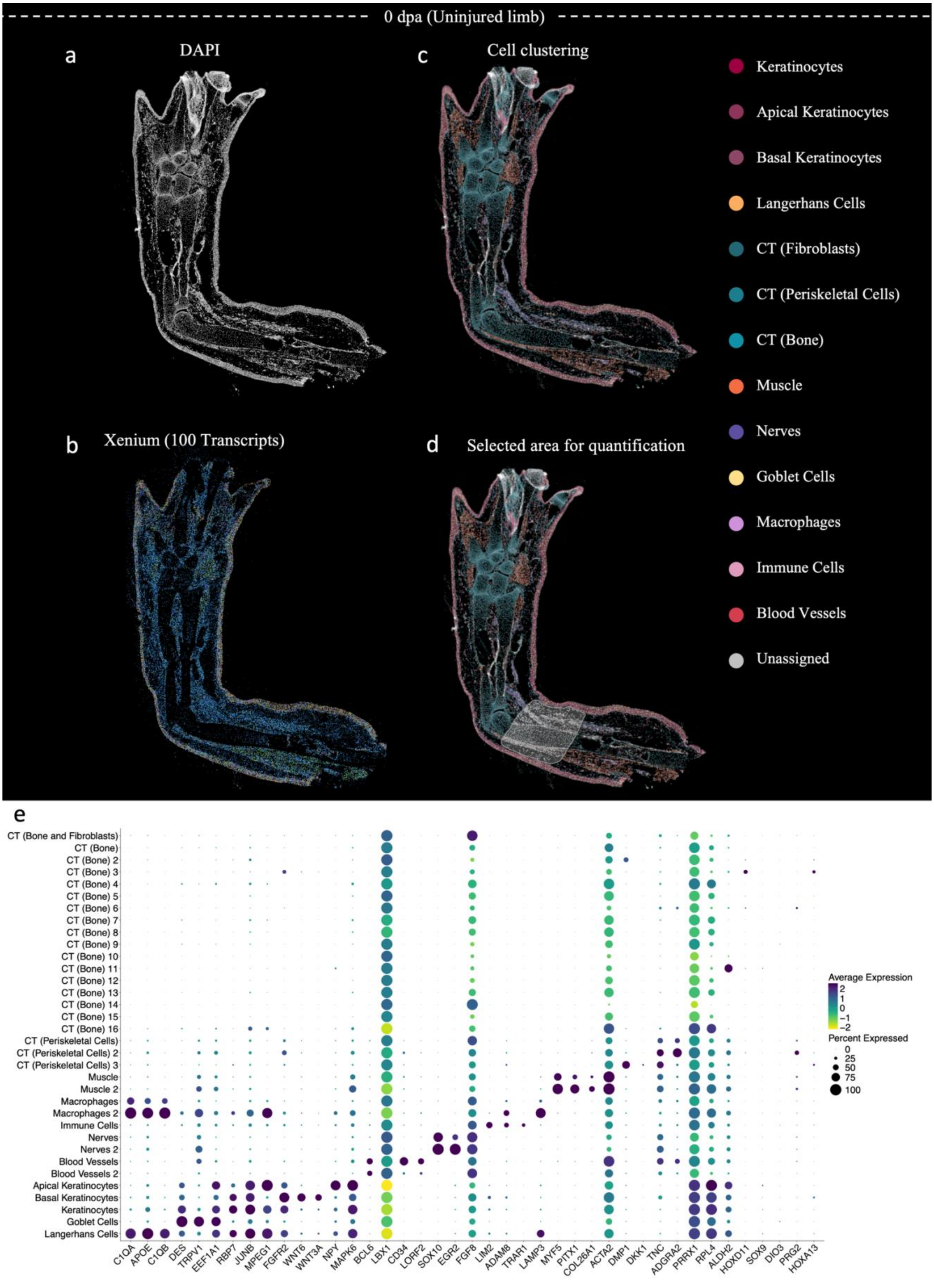

0 dpa (Uninjured limb)
a
DAPI
b
Xenium (100 Transcripts)
c
Cell clustering
d
Selected area for quantification
Keratinocytes
Apical Keratinocytes
Basal Keratinocytes
Langerhans Cells
CT (Fibroblasts)
CT (Periskeletal Cells)
CT (Bone)
Muscle
Nerves
Goblet Cells
Macrophages
Immune Cells
Blood Vessels
Unassigned
e
CT (Bone and Fibroblasts)
CT (Bone)
CT (Bone) 2
CT (Bone) 3
CT (Bone) 4
CT (Bone) 5
CT (Bone) 6
CT (Bone) 7
CT (Bone) 8
CT (Bone) 9
CT (Bone) 10
CT (Bone) 11
CT (Bone) 12
CT (Bone) 13
CT (Bone) 14
CT (Bone) 15
CT (Bone) 16
CT (Periskeletal Cells)
CT (Periskeletal Cells) 2
CT (Periskeletal Cells) 3
Muscle
Muscle 2
Macrophages
Macrophages 2
Immune Cells
Nerves
Nerves 2
Blood Vessels
Blood Vessels 2
Apical Keratinocytes
Basal Keratinocytes
Keratinocytes
Goblet Cells
Langerhans Cells
C1QA
APOE
C1QB
DES
TRPV1
EEF1A1
RBP7
JUNB
MPEG1
FGFR2
WNT6
WNT3A
NPY
MAPK6
BCL6
LBX1
CD34
LORF2
SOX10
EGR2
FGF8
LIM2
ADAM8
TRAR1
LAMP3
MYF5
PITX1
COL26A1
ACTA2
DMP1
DKK1
TNC
ADGRA2
PRRX1
RPL4
ALDH2
HOXD11
SOX9
DIO3
PRG2
HOXA13
Average Expression
2
1
0
−1
−2
Percent Expressed
0
25
50
75
100

**Supplementary Figure 2. Characterization of cellular identity using Xenium analysis in an uninjured axolotl forelimb**. **a-d**, Xenium analysis of a mature axolotl forelimb (uninjured limb) using 100 probes. Cell identities were determined through a combination of DAPI-based nuclear segmentation (a) and detected transcript profiles (b), resulting in the identification of 20 distinct cell clusters (c). Shadowed area (d) corresponds to the selected area used for the quantification of cellular identities for Fig 1d. **e**, Dot plot illustrating the top 3 genes defining each cluster, with color-coded average expression levels (ranging from yellow to dark blue), and dot size representing the percentage of cells within the corresponding cluster (shown on the y-axis) that express the corresponding gene (shown on the x-axis).

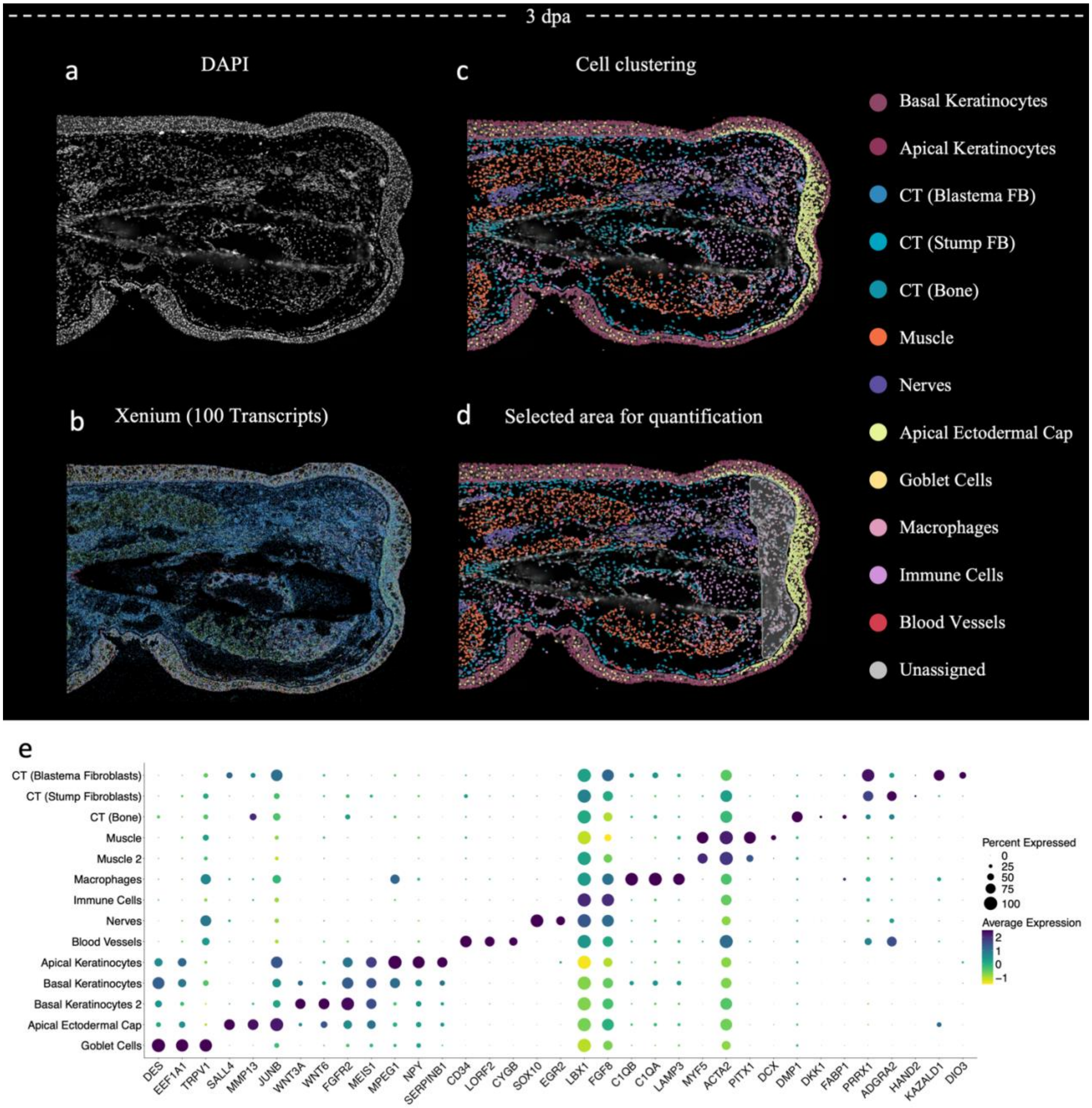
3 dpa
a
DAPI
b
Xenium (100 Transcripts)
c
Cell clustering
d
Selected area for quantification
Basal Keratinocytes
Apical Keratinocytes
CT (Blastema FB)
CT (Stump FB)
CT (Bone)
Muscle
Nerves
Apical Ectodermal Cap
Goblet Cells
Macrophages
Immune Cells
Blood Vessels
Unassigned
e
CT (Blastema Fibroblasts)
CT (Stump Fibroblasts)
CT (Bone)
Muscle
Muscle 2
Macrophages
Immune Cells
Nerves
Blood Vessels
Apical Keratinocytes
Basal Keratinocytes
Basal Keratinocytes 2
Apical Ectodermal Cap
Goblet Cells
DES
EEF1A1
TRPV1
SALL4
MMP13
JUNB
WNT3A
WNT6
FGFR2
MEIS1
MPEG1
NPY
SERPINB1
CD34
LORF2
CYGB
SOX10
EGR2
LBX1
FGF8
C1QB
C1QA
LAMP3
MYF5
ACTA2
PITX1
DCX
DMP1
DKK1
FABP1
PRRX1
ADGRA2
HAND2
KAZALD1
DIO3
Percent Expressed
0
25
50
75
100
Average Expression
2
1
0
−1

**Supplementary Figure 3. Characterization of cellular identity using Xenium analysis in a 3-dpa axolotl forelimb**. **a-c**, Xenium analysis of 3-dpa forelimb using 100 probes. Cell identities were determined through a combination of DAPI-based nuclear segmentation (a) and detected transcript profiles (b), resulting in the identification of 15 distinct cell clusters (c). Shadowed area (d) corresponds to the selected area used for the quantification of cellular identities for Fig 1d. **e**, Dot plot illustrating the top 3 genes defining each cluster, with color-coded average expression levels (ranging from yellow to dark blue), and dot size representing the percentage of cells within the corresponding cluster (shown on the y-axis) that express the corresponding gene (shown on the x-axis).

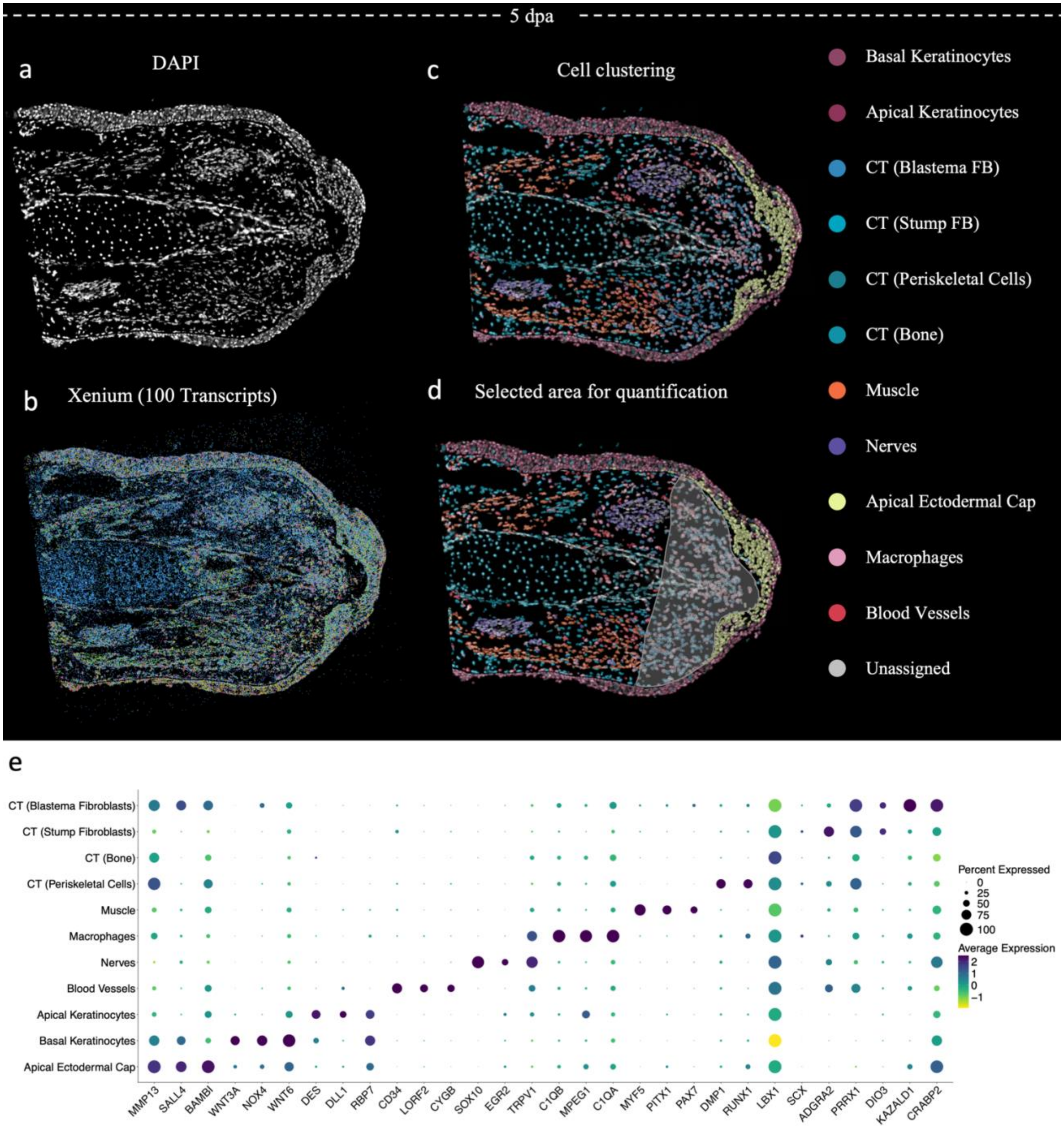
5 dpa
a
DAPI
b
Xenium (100 Transcripts)
c
Cell clustering
d
Selected area for quantification
Basal Keratinocytes
Apical Keratinocytes
CT (Blastema FB)
CT (Stump FB)
CT (Periskeletal Cells)
CT (Bone)
Muscle
Nerves
Apical Ectodermal Cap
Macrophages
Blood Vessels
Unassigned
e
CT (Blastema Fibroblasts)
CT (Stump Fibroblasts)
CT (Bone)
CT (Periskeletal Cells)
Muscle
Macrophages
Nerves
Blood Vessels
Apical Keratinocytes
Basal Keratinocytes
Apical Ectodermal Cap
MMP13
SALL4
BAMBI
WNT3A
NOX4
WNT6
DES
DLL1
RBP7
CD34
LORF2
CYGB
SOX10
EGR2
TRPV1
C1QB
MPEG1
C1QA
MYF5
PITX1
PAX7
DMP1
RUNX1
LBX1
SCX
ADGRA2
PRRX1
DIO3
KAZALD1
CRABP2
Percent Expressed
0
25
50
75
100
Average Expression
2
1
0
−1

**Supplementary Figure 4. Characterization of cellular identity using Xenium analysis in a 5-dpa axolotl forelimb**. **a-c**, Xenium analysis of 5-dpa forelimb using 100 probes. Cell identities were determined through a combination of DAPI-based nuclear segmentation (a) and detected transcript profiles (b), resulting in the identification of 13 distinct cell clusters (c). Shadowed area (d) corresponds to the selected area used for the quantification of cellular identities for Fig 1d. **e**, Dot plot illustrating the top 3 genes defining each cluster, with color-coded average expression levels (ranging from yellow to dark blue), and dot size representing the percentage of cells within the corresponding cluster (shown on the y-axis) that express the corresponding gene (shown on the x-axis).

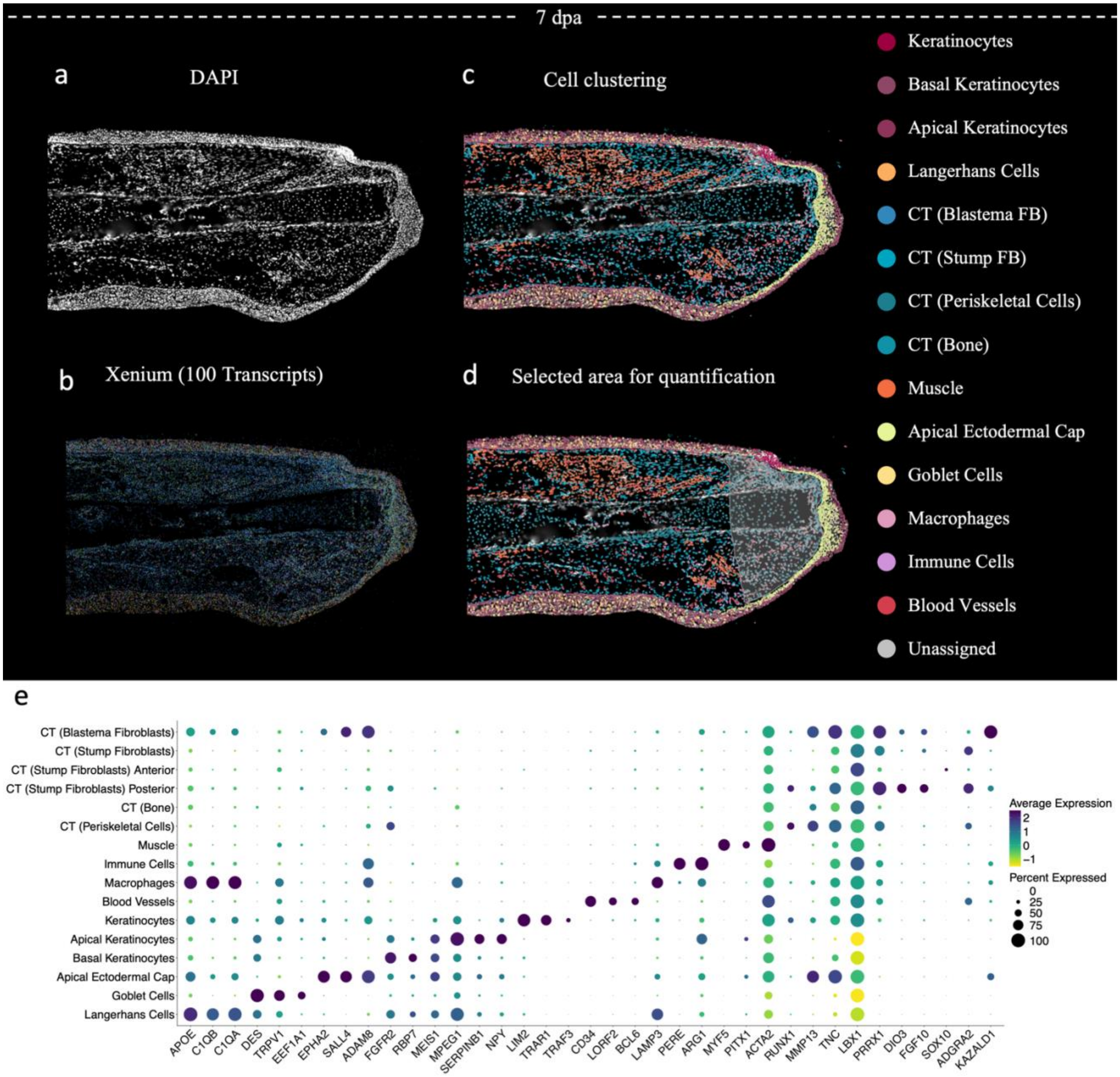
7 dpa
a
DAPI
b
Xenium (100 Transcripts)
c
Cell clustering
d
Selected area for quantification
Keratinocytes
Basal Keratinocytes
Apical Keratinocytes
Langerhans Cells
CT (Blastema FB)
CT (Stump FB)
CT (Periskeletal Cells)
CT (Bone)
Muscle
Apical Ectodermal Cap
Goblet Cells
Macrophages
Immune Cells
Blood Vessels
Unassigned
e
CT (Blastema Fibroblasts)
CT (Stump Fibroblasts)
CT (Stump Fibroblasts) Anterior
CT (Stump Fibroblasts) Posterior
CT (Bone)
CT (Periskeletal Cells)
Muscle
Immune Cells
Macrophages
Blood Vessels
Keratinocytes
Apical Keratinocytes
Basal Keratinocytes
Apical Ectodermal Cap
Goblet Cells
Langerhans Cells
APOE
C1QB
C1QA
DES
TRPV1
EEF1A1
EPHA2
SALL4
ADAM8
FGFR2
RBP7
MEIS1
MPEG1
SERPINB1
NPY
LIM2
TRAR1
TRAF3
CD34
LORF2
BCL6
LAMP3
PERE
ARG1
MYF5
PITX1
ACTA2
RUNX1
MMP13
TNC
LBX1
PRRX1
DIO3
FGF10
SOX10
ADGRA2
KAZALD1
Average Expression
2
1
0
−1
Percent Expressed
0
25
50
75
100

**Supplementary Figure 5. Characterization of cellular identity using Xenium analysis in a 7-dpa axolotl forelimb**. **a-c**, Xenium analysis of 7-dpa forelimb using 100 probes. Cell identities were determined through a combination of DAPI-based nuclear segmentation (a) and detected transcript profiles (b), resulting in the identification of 15 distinct cell clusters (c). Shadowed area (d) corresponds to the selected area used for the quantification of cellular identities for Fig 1d. **e**, Dot plot illustrating the top 3 genes defining each cluster, with color-coded average expression levels (ranging from yellow to dark blue), and dot size representing the percentage of cells within the corresponding cluster (shown on the y-axis) that express the corresponding gene (shown on the x-axis).

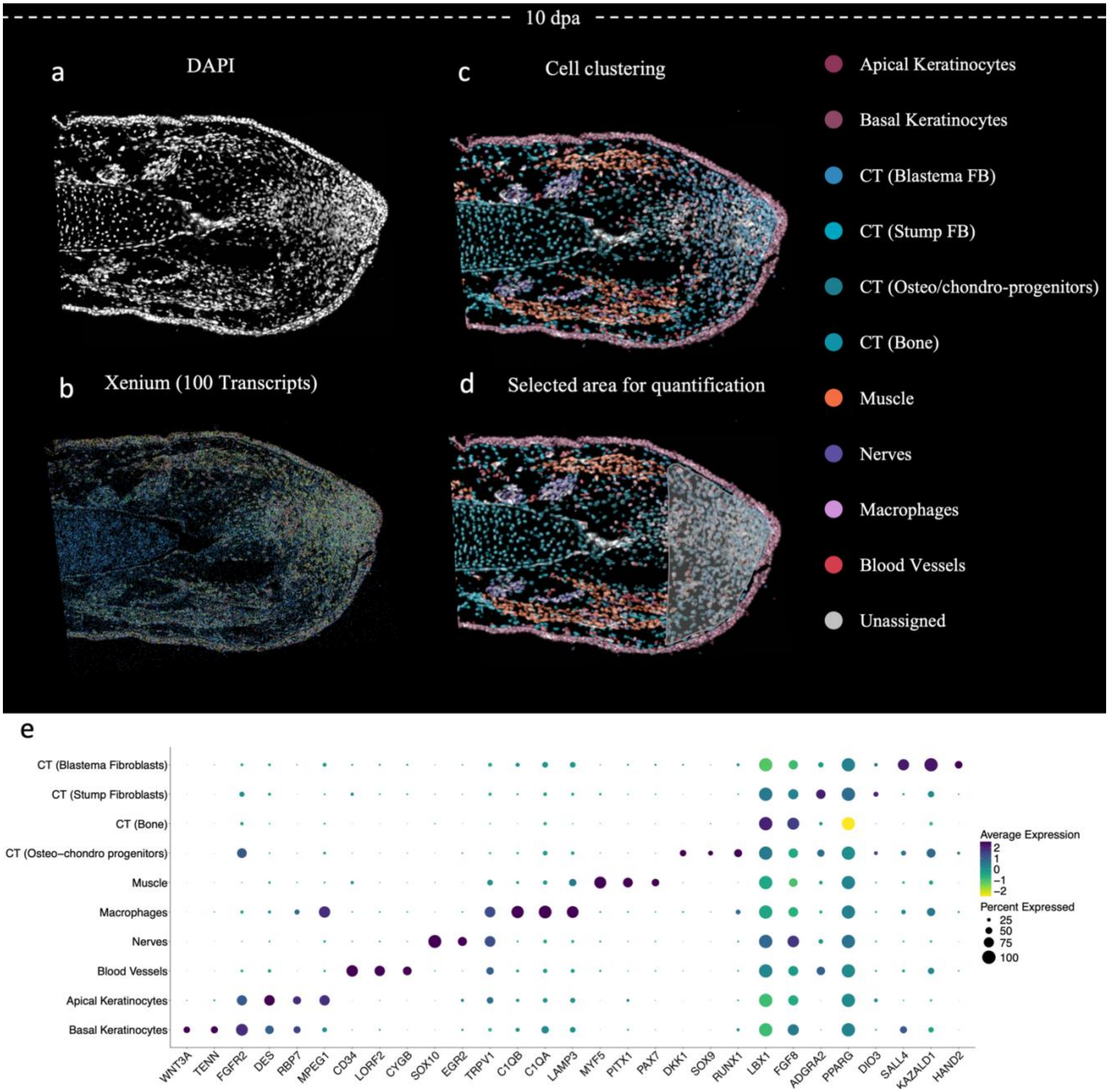

10 dpa
a
DAPI
c
Cell clustering
b
Xenium (100 Transcripts)
d
Selected area for quantification
Apical Keratinocytes
Basal Keratinocytes
CT (Blastema FB)
CT (Stump FB)
CT (Osteo/chondro-progenitors)
CT (Bone)
Muscle
Nerves
Macrophages
Blood Vessels
Unassigned
e
CT (Blastema Fibroblasts)
CT (Stump Fibroblasts)
CT (Bone)
CT (Osteo−chondro progenitors)
Muscle
Macrophages
Nerves
Blood Vessels
Apical Keratinocytes
Basal Keratinocytes
WNT3A
TENN
FGFR2
DES
RBP7
MPEG1
CD34
LORF2
CYGB
SOX10
EGR2
TRPV1
C1QB
C1QA
LAMP3
MYF5
PITX1
PAX7
DKK1
SOX9
RUNX1
LBX1
FGF8
ADGRA2
PPARG
DIO3
SALL4
KAZALD1
HAND2
Average Expression
2
1
0
−1
−2
Percent Expressed
25
50
75
100

**Supplementary Figure 6. Characterization of cellular identity using Xenium analysis in a 10-dpa axolotl forelimb**. **a-c**, Xenium analysis of 10-dpa forelimb using 100 probes. Cell identities were determined through a combination of DAPI-based nuclear segmentation (a) and detected transcript profiles (b), resulting in identification of 15 distinct cell clusters (c). Shadowed area (d) corresponds to the selected area used for the quantification of cellular identities for Fig 1d. **e**, Dot plot illustrating the top 3 genes defining each cluster, with color-coded average expression levels (ranging from yellow to dark blue), and dot size representing the percentage of cells within the corresponding cluster (shown on the y-axis) that express the corresponding gene (shown on the x-axis).

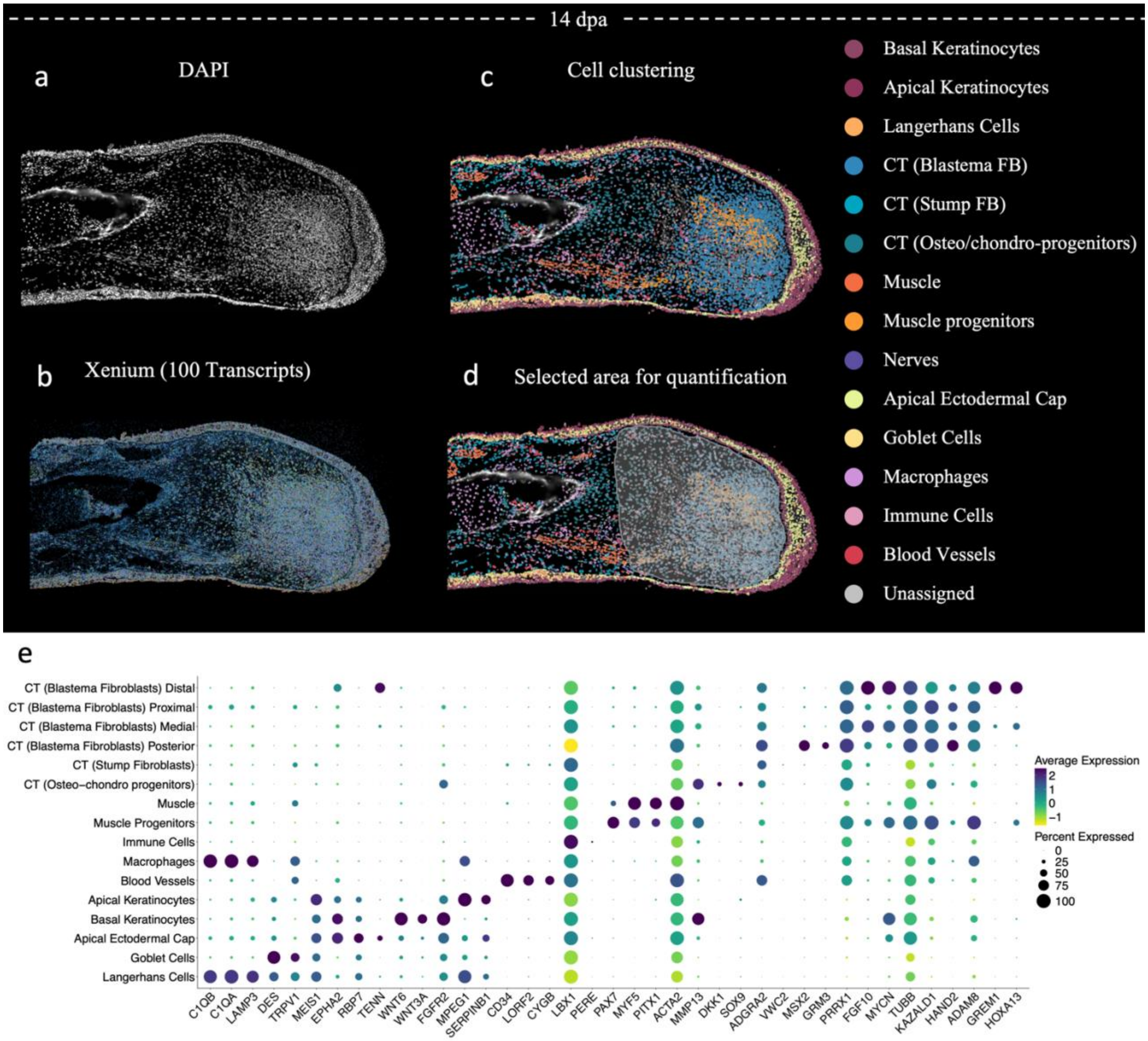
14 dpa
a
DAPI
b
Xenium (100 Transcripts)
c
Cell clustering
d
Selected area for quantification
Basal Keratinocytes
Apical Keratinocytes
Langerhans Cells
CT (Blastema FB)
CT (Stump FB)
CT (Osteo/chondro-progenitors)
Muscle
Muscle progenitors
Nerves
Apical Ectodermal Cap
Goblet Cells
Macrophages
Immune Cells
Blood Vessels
Unassigned
e
CT (Blastema Fibroblasts) Distal
CT (Blastema Fibroblasts) Proximal
CT (Blastema Fibroblasts) Medial
CT (Blastema Fibroblasts) Posterior
CT (Stump Fibroblasts)
CT (Osteo−chondro progenitors)
Muscle
Muscle Progenitors
Immune Cells
Macrophages
Blood Vessels
Apical Keratinocytes
Basal Keratinocytes
Apical Ectodermal Cap
Goblet Cells
Langerhans Cells
C1QB
C1QA
LAMP3
DES
TRPV1
MEIS1
EPHA2
RBP7
TENN
WNT6
WNT3A
FGFR2
MPEG1
SERPINB1
CD34
LORF2
CYGB
LBX1
PERE
PAX7
MYF5
PITX1
ACTA2
MMP13
DKK1
SOX9
ADGRA2
VWC2
MSX2
GRM3
PRRX1
FGF10
MYCN
TUBB
KAZALD1
HAND2
ADAM8
GREM1
HOXA13
Average Expression
2
1
0
−1
Percent Expressed
0
25
50
75
100

**Supplementary Figure 7. Characterization of cellular identity using Xenium analysis in a 14-dpa axolotl forelimb**. **a-c**, Xenium analysis of 14-dpa forelimb using 100 probes. Cell identities were determined through a combination of DAPI-based nuclear segmentation (a) and detected transcript profiles (b), resulting in the identification of 18 distinct cell clusters (c). Shadowed area (d) corresponds to the selected area used for the quantification of cellular identities for Fig 1d. **e**, Dot plot illustrating the top 3 genes defining each cluster, with color-coded average expression levels (ranging from yellow to dark blue), and dot size representing the percentage of cells within the corresponding cluster (shown on the y-axis) that express the corresponding gene (shown on the x-axis).

a

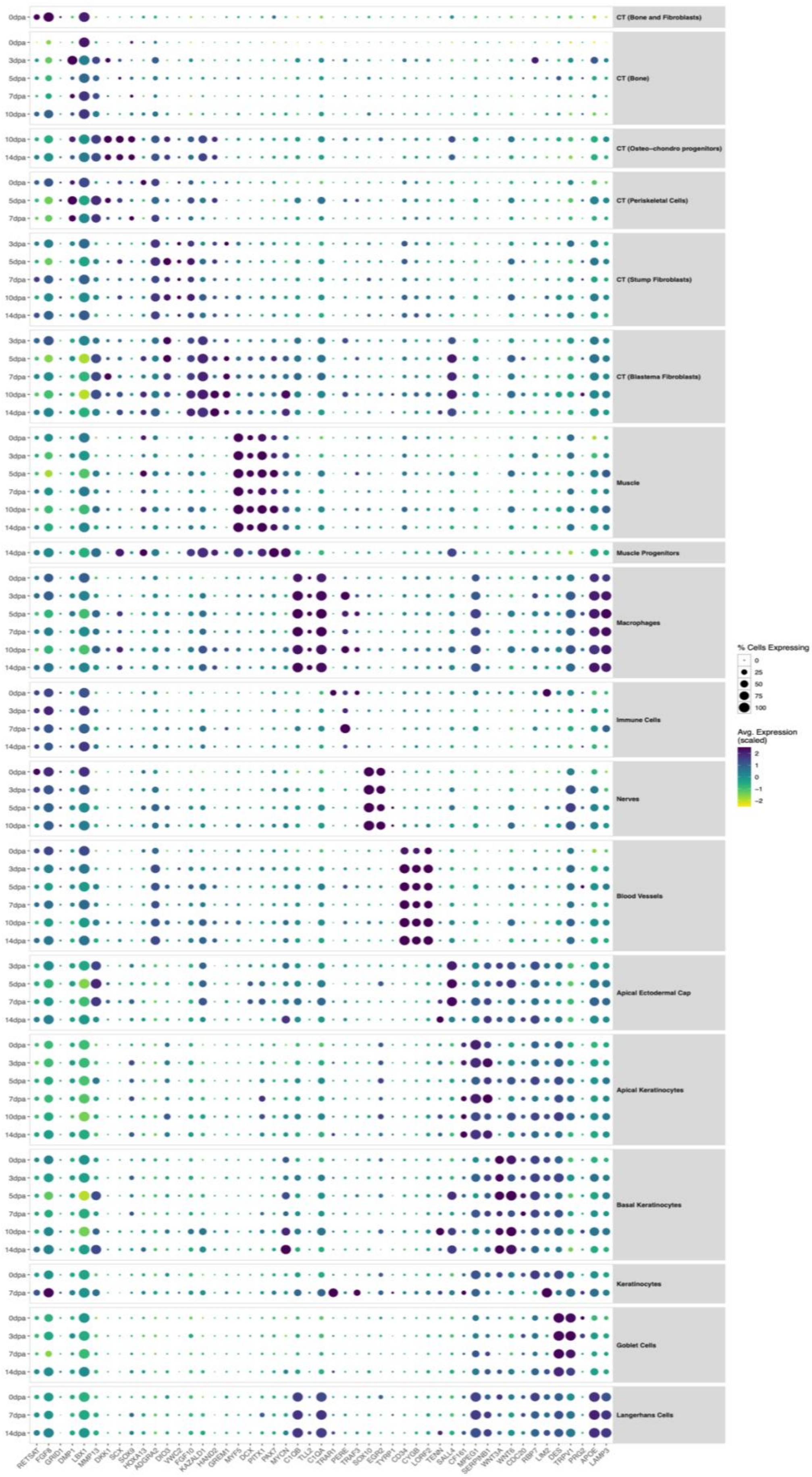
CT (Bone and Fibroblasts)
CT (Bone)
CT (Osteo–chondro progenitors)
CT (Periskeletal Cells)
CT (Stump Fibroblasts)
CT (Blastema Fibroblasts)
Muscle
Muscle Progenitors
Macrophages
Immune Cells
Nerves
Blood Vessels
Apical Ectodermal Cap
Apical Keratinocytes
Basal Keratinocytes
Keratinocytes
Goblet Cells
Langerhans Cells
0dpa
3dpa
5dpa
7dpa
10dpa
14dpa
RETSAT FGF8 GRID1 DMP1 LBX1 MMP13 DKK1 SCX SOX9 HOXA13 ADGRA2 DIO3 VWC2 FGF10 KAZALD1 HAND2 GREM1 MYF5 DCX PITX1 PAX7 MYCN C1QB TLL2 C1QA TRAP1 PERE TRAF3 SOX10 EGR2 TYRP1 CD34 CYGB LORF2 TENN SALL4 CF161 MPEG1 SERPINB1 WNT3A WNT6 CDC20 RBP7 LIM2 DES TRPV1 PRG2 APOE LAMP3
% Cells Expressing
0
25
50
75
100
Avg. Expression (scaled)
2
1
0
−1
−2

**Supplementary Figure 8. Molecular conservation of Xenium clusters over time**. **a,** Dot plot illustrating the top 3 genes defining each cluster and their variation over each time point in the Xenium analysis.

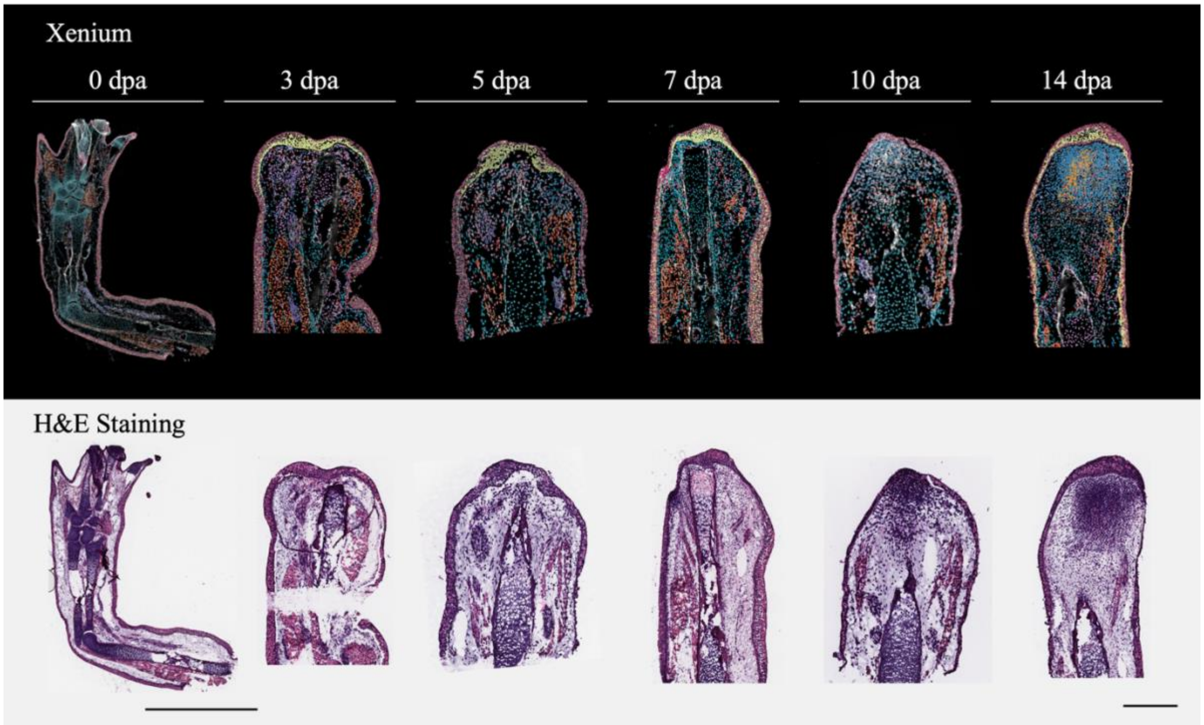


**Supplementary Figure 9. H&E staining from Xenium experiments**. Xenium analysis performed in an uninjured limb and at 3, 5, 7, 10, and 14 dpa (top panel). Hematoxylin and eosin (H&E) staining of sections after the Xenium experiment (bottom panel). Scale bars: 5 mm for the uninjured limb and 1 mm for the subsequent time points.

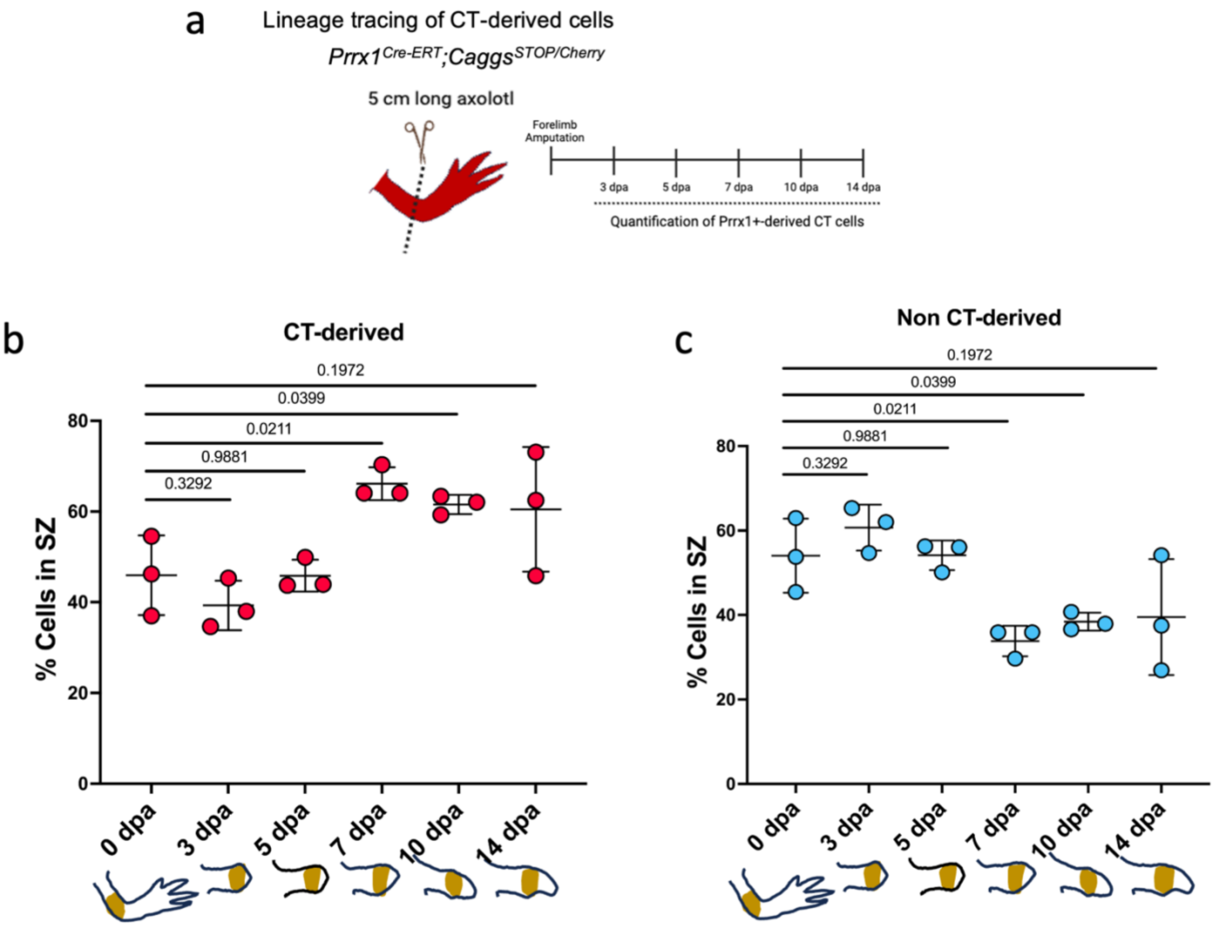


**Supplementary Figure 10. The percentage of CT-derived cells increases in the source zone during blastema formation**. **a**, Schematic representation of the experimental design detailing lineage tracing experiments conducted with converted *Prrx1*$^{Cre\text{-}ERT}$;*Caggs*$^{STOP/Cherry}$ axolotls as detailed in Figure 1d. **b-c**, Quantitative analysis of CT-derived cells (b) and non-CT-derived cells (c) within the source zone. The source zone is defined as the 500-micrometers region proximal to the amputation plane, excluding the limb epidermis (brown shadowed area) (n = 3). Statistical significance was determined using unpaired t-tests, with corresponding p-values annotated in the graph for each comparison.

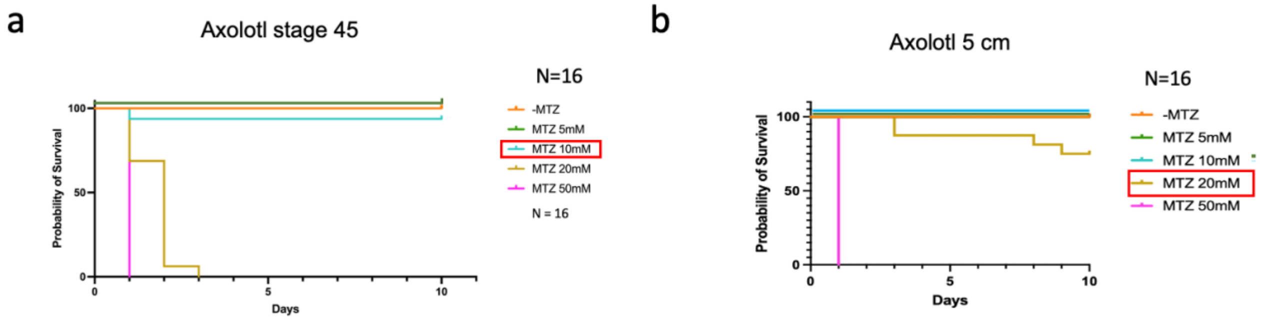


**Supplementary Figure 11. Optimization of metronidazole (MTZ) concentrations for axolotl larvae and juveniles. a, b,** Survival analysis of axolotls exposed to different MTZ concentrations at different developmental stages. Survival plots for d/d axolotl larvae at stage 45 (a), and for 5-cm-long juvenile axolotls (b). For every experiment, independent treatments were conducted using MTZ at 5 mM, 10 mM, 20 mM, and 50 mM. (n = 16 for each condition). The red box indicates the preferred MTZ concentration for each axolotl age.

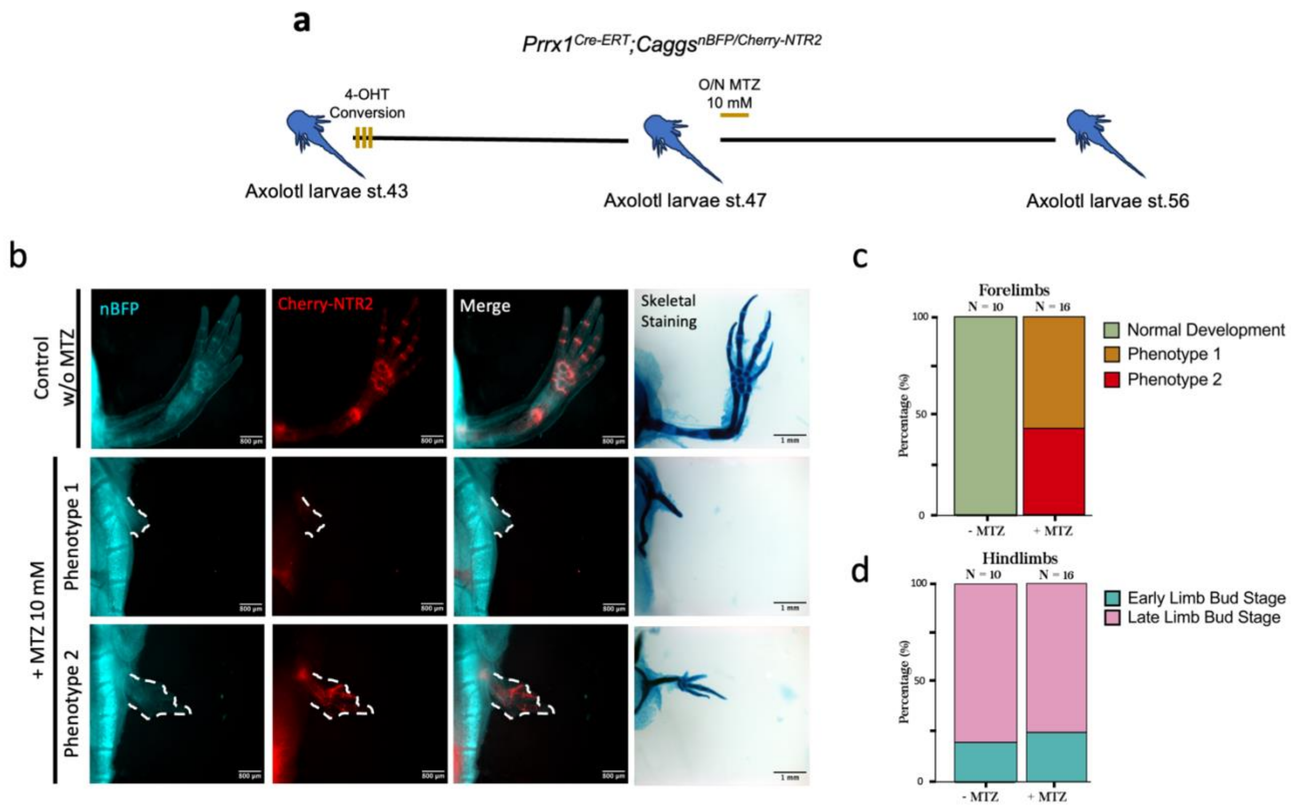


**Supplementary Figure 12. CT-derived cells are essential for limb development**. **a**, Schematic representation of the experimental design detailing experiments conducted using converted *Prrx1*$^{Cre-ERT}$;*Caggs*$^{nBFP/NTR2}$ axolotls. To induce conversion, three treatments of 4-hydroxytamoxifen (4-OHT) were performed at stage 43, spaced every other day. One overnight treatment of 10 mM MTZ was done 10 days post-first 4-OHT treatment (axolotl larvae stage 47), and limbs were imaged and harvested at stage 56. **b**, Stereoscopic images of the axolotl limb at stage 56 illustrating the final limb phenotype. Converted *Prrx1*$^{Cre-ERT}$;*Caggs*$^{\mathrm{nBFP/Cherry\text{-}NTR2}}$ (red) fluorescence is shown. Scale bar = 500 µm. The right-hand column shows the skeletal elements of every condition via alcian blue/alizarin staining. Scale bar = 1 mm. **c-d**, Phenotypic analysis with alcian blue/alizarin red of developed forelimbs (c) and hindlimbs (d). Hindlimbs are used as a control as they develop later and are not exposed to 4-OHT treatment at stage 43. Normal phenotype refers to proper development of the stylopod, zeugopod, and autopod segments. Phenotype 1 refers to a spike-like formation, and phenotype 2 refers to the development of stylopod and digit rays (autopod) but lacking the intermediate zeugopod segment.

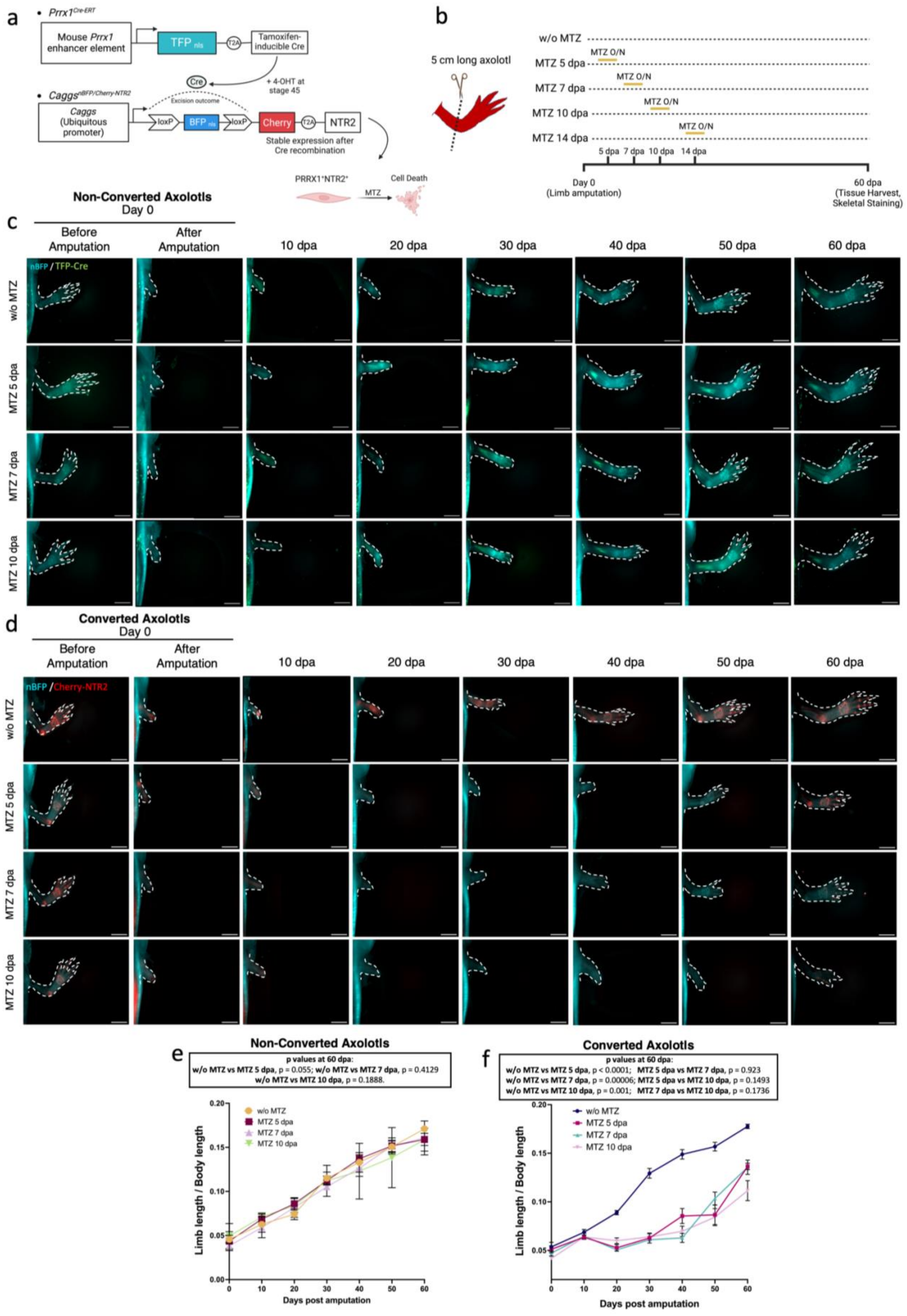
a
Prrx1Cre-ERT
Mouse Prrx1 enhancer element
TFP nls
T2A
Tamoxifen-inducible Cre
Cre
+ 4-OHT at stage 45
CaggsnBFP/Cherry-NTR2
Excision outcome
Caggs (Ubiquitous promoter)
loxP
BFP nls
loxP
Cherry
T2A
NTR2
Stable expression after Cre recombination
PRRX1+NTR2+
MTZ
Cell Death
b
5 cm long axolotl
w/o MTZ
MTZ 5 dpa
MTZ 7 dpa
MTZ 10 dpa
MTZ 14 dpa
MTZ O/N
5 dpa
7 dpa
10 dpa
14 dpa
Day 0 (Limb amputation)
60 dpa (Tissue Harvest, Skeletal Staining)
c
Non-Converted Axolotls
Day 0
Before Amputation
After Amputation
10 dpa
20 dpa
30 dpa
40 dpa
50 dpa
60 dpa
nBFP / TFP-Cre
w/o MTZ
MTZ 5 dpa
MTZ 7 dpa
MTZ 10 dpa
d
Converted Axolotls
Day 0
Before Amputation
After Amputation
10 dpa
20 dpa
30 dpa
40 dpa
50 dpa
60 dpa
nBFP / Cherry-NTR2
w/o MTZ
MTZ 5 dpa
MTZ 7 dpa
MTZ 10 dpa
e
Non-Converted Axolotls
p values at 60 dpa:
w/o MTZ vs MTZ 5 dpa, p = 0.055; w/o MTZ vs MTZ 7 dpa, p = 0.4129
w/o MTZ vs MTZ 10 dpa, p = 0.1888.
w/o MTZ
MTZ 5 dpa
MTZ 7 dpa
MTZ 10 dpa
Limb length / Body length
Days post amputation
f
Converted Axolotls
p values at 60 dpa:
w/o MTZ vs MTZ 5 dpa, p < 0.0001; MTZ 5 dpa vs MTZ 7 dpa, p = 0.923
w/o MTZ vs MTZ 7 dpa, p = 0.00006; MTZ 5 dpa vs MTZ 10 dpa, p = 0.1493
w/o MTZ vs MTZ 10 dpa, p = 0.001; MTZ 7 dpa vs MTZ 10 dpa, p = 0.1736
w/o MTZ
MTZ 5 dpa
MTZ 7 dpa
MTZ 10 dpa
Limb length / Body length
Days post amputation

**Supplementary Figure 13. Genetic ablation of CT-derived cells at early time points delays limb regeneration.** **a-b** Schematic representation of the cell-ablation experiments in converted $Prrx1^{Cre\text{-}ERT};Caggs^{nBFP/Cherry\text{-}NTR2}$ axolotls. To assess the impact of CT cell depletion, a single overnight 20 mM MTZ treatment was performed at 5, 7, or 10 dpa, and limb regeneration was monitored until 60 dpa (b) **c-d**, Stereoscopic images capturing the regeneration progress over the 60-day regeneration period. The limbs of both non-converted $Prrx1^{Cre\text{-}ERT};Caggs^{nBFP/Cherry\text{-}NTR2}$ (nBFP expression in cyan, and TFP-Cre expression in green) (c), and converted (nBFP expression in cyan, and Cherry-NTR2.0 expression in red) axolotls (d) are shown. Dotted lines outline the limb periphery. Scale bar = 1 mm (n = 3 for each condition using non-converted axolotls, and n = 12 for each condition using converted axolotls). **e-f**, Quantitative assessment of limb length/body length ratio of the regenerated limb of non-converted (e) and converted (f) axolotls over the course of the entire experiment (0 dpa to 60 dpa). The quantified limb region extends from the axolotl's torso to the most distal part of the limb. Values are shown as mean ± SD (n = 3 for non-converted axolotls and n = 12 for converted axolotls). Statistical significance of values at 60 dpa was determined using two-way unpaired t-tests, with corresponding p-values annotated at the top of the graph for each comparison.

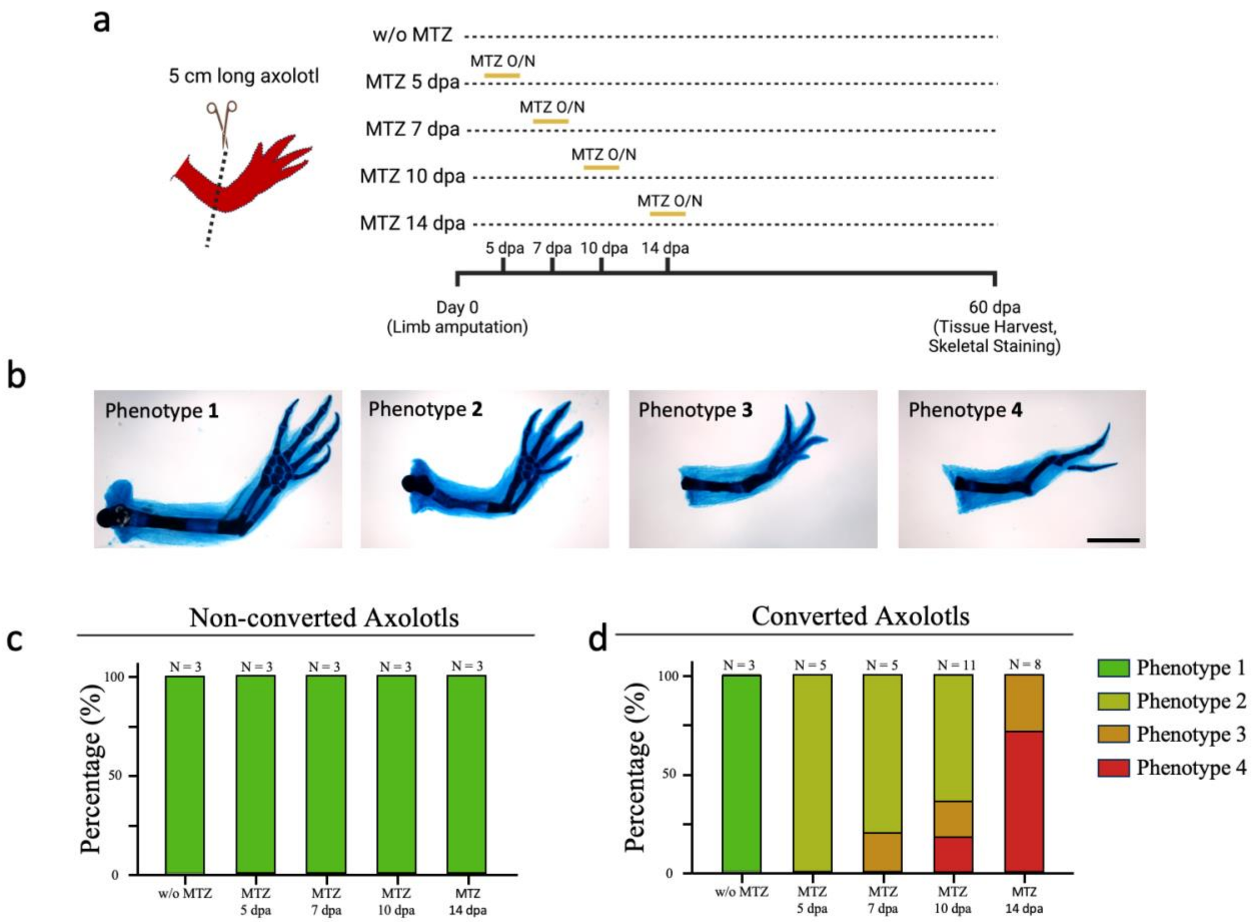

a
5 cm long axolotl
w/o MTZ
MTZ 5 dpa
MTZ 7 dpa
MTZ 10 dpa
MTZ 14 dpa
MTZ O/N
5 dpa
7 dpa
10 dpa
14 dpa
Day 0
(Limb amputation)
60 dpa
(Tissue Harvest,
Skeletal Staining)
b
Phenotype 1
Phenotype 2
Phenotype 3
Phenotype 4
c
Non-converted Axolotls
Percentage (%)
N = 3
w/o MTZ
MTZ 5 dpa
MTZ 7 dpa
MTZ 10 dpa
MTZ 14 dpa
d
Converted Axolotls
N = 3
N = 5
N = 5
N = 11
N = 8
Phenotype 1
Phenotype 2
Phenotype 3
Phenotype 4

**Supplementary Figure 14. Limb phenotype severity increases when CT-derived cells are ablated at later stages of limb regeneration. a,** Schematic representation of the experimental design and limb phenotype analysis following genetic CT-cell ablation in $Prrx1^{Cre\text{-}ERT};Caggs^{nBFP/Cherry\text{-}NTR2}$ axolotls. Animals underwent a single overnight 20mM MTZ treatment at 5, 7, 10, or 14 dpa, followed by regeneration assessment at 60 dpa. **b**, Widefield microscopy images of alcian blue/alizarin red staining illustrating the skeletal phenotype at 60 dpa. Phenotype 1 corresponds to a properly segmented limb with a normal size for all skeletal elements, serving as a control. Phenotype 2 corresponds to a properly segmented limb but with smaller skeletal elements than phenotype 1. Phenotype 3 presents loss of posterior carpal bones and posterior phalanges, along with reduced ulna and posterior digit bones. Phenotype 4 corresponds to a complete loss of elbow articulation, along with an absence of ulna, carpal bones, and posterior digits. Scale bar = 2mm. **c-d**, Percentages of the different phenotypes in each of the five experimental conditions for non-converted (c) and converted (d) axolotls. n for each condition is mentioned in the graph.

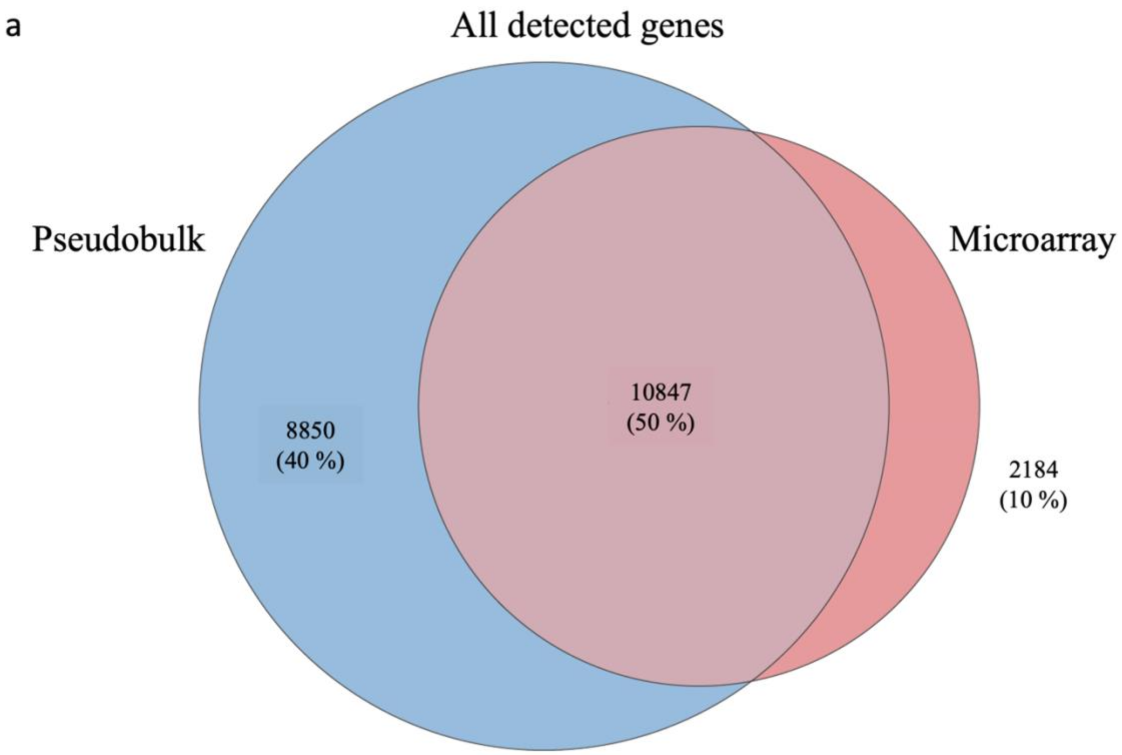


**Supplementary Figure 15. Total of genes detected by microarray and pseudo-bulk single-cell (sc) datasets from flow-sorted CT-derived cells. a,** Venn diagram comparing the numbers of genes detected by pseudobulk analysis (blue), and microarray (red).

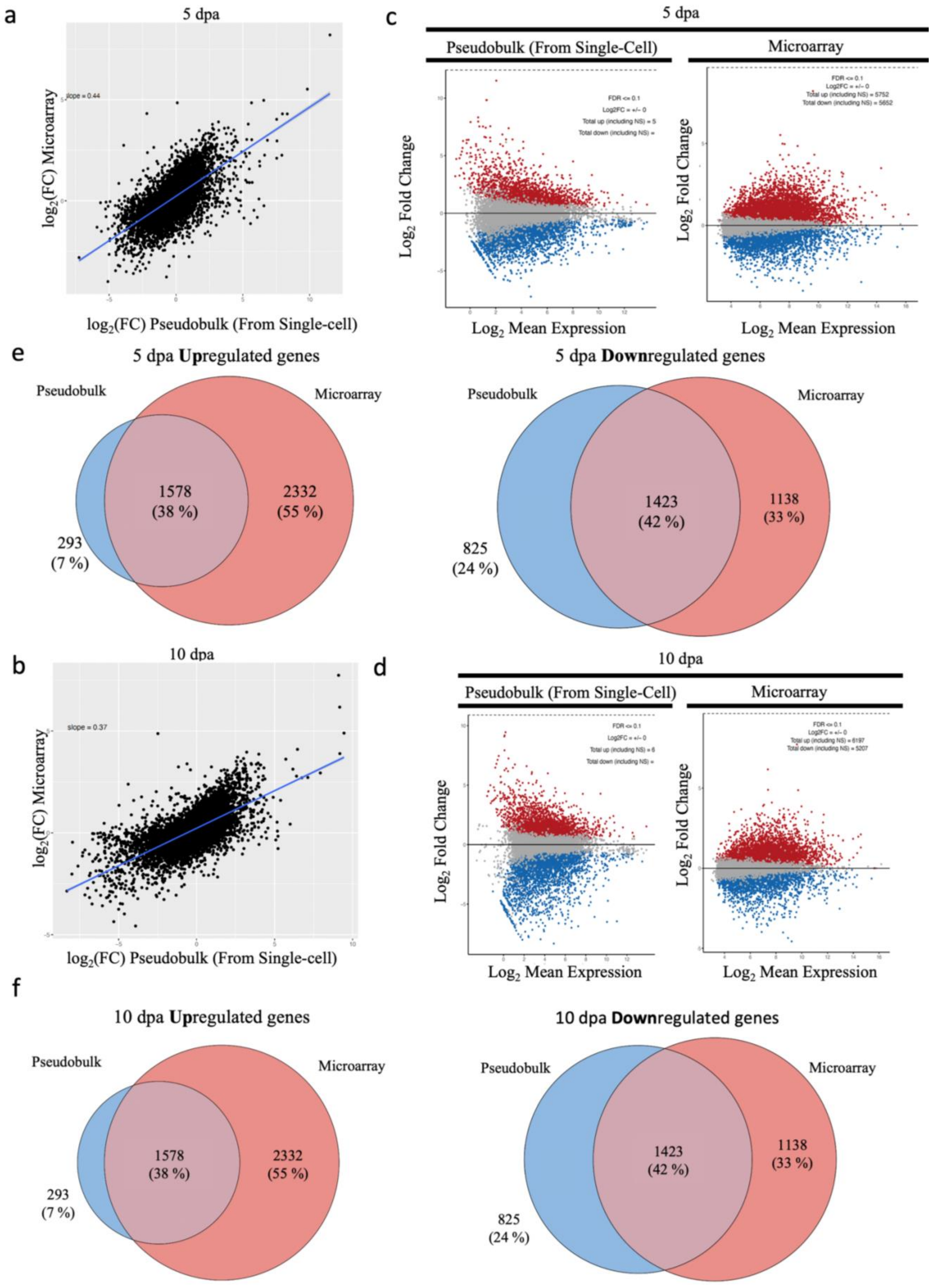
a
5 dpa
slope = 0.44
log2(FC) Microarray
log2(FC) Pseudobulk (From Single-cell)
c
5 dpa
Pseudobulk (From Single-Cell)
Microarray
FDR <= 0.1
Log2FC = +/- 0
Total up (including NS) = 5
Total down (including NS) =
Total up (including NS) = 5752
Total down (including NS) = 5652
Log2 Fold Change
Log2 Mean Expression
e
5 dpa Upregulated genes
Pseudobulk
Microarray
1578
(38 %)
2332
(55 %)
293
(7 %)
5 dpa Downregulated genes
Pseudobulk
Microarray
1423
(42 %)
1138
(33 %)
825
(24 %)
b
10 dpa
slope = 0.37
log2(FC) Microarray
log2(FC) Pseudobulk (From Single-cell)
d
10 dpa
Pseudobulk (From Single-Cell)
Microarray
FDR <= 0.1
Log2FC = +/- 0
Total up (including NS) = 6
Total down (including NS) =
Total up (including NS) = 6197
Total down (including NS) = 5207
Log2 Fold Change
Log2 Mean Expression
f
10 dpa Upregulated genes
Pseudobulk
Microarray
1578
(38 %)
2332
(55 %)
293
(7 %)
10 dpa Downregulated genes
Pseudobulk
Microarray
1423
(42 %)
1138
(33 %)
825
(24 %)

**Supplementary Figure 16. Comparative analysis of microarray and pseudo-bulk single-cell (sc) datasets from flow-sorted CT-derived cells.** Transcriptomic data comparison between the microarray analysis and the scRNAseq data previously published [8] (considered as a pseudobulk RNAseq) at 5 and 10 dpa. (**a,b**) Correlation plots of log 2 fold-change ($log_2FC$) estimates for each gene (black dots) between microarray and pseudobulk RNAseq at 5 dpa (a) and 10 dpa (b). (**c,d**) MA plots displaying gene expression differences for pseudobulk RNA-seq (left) and microarray (right) for 5 dpa (c) and 10 dpa (d). Upregulated Differential Expressed Genes (DEGs) are highlighted. (**e,f**) Venn diagrams comparing the numbers of upregulated (top row) and downregulated (bottom row) DEGs detected with both technologies at 5 dpa (e) and 10 dpa (f).

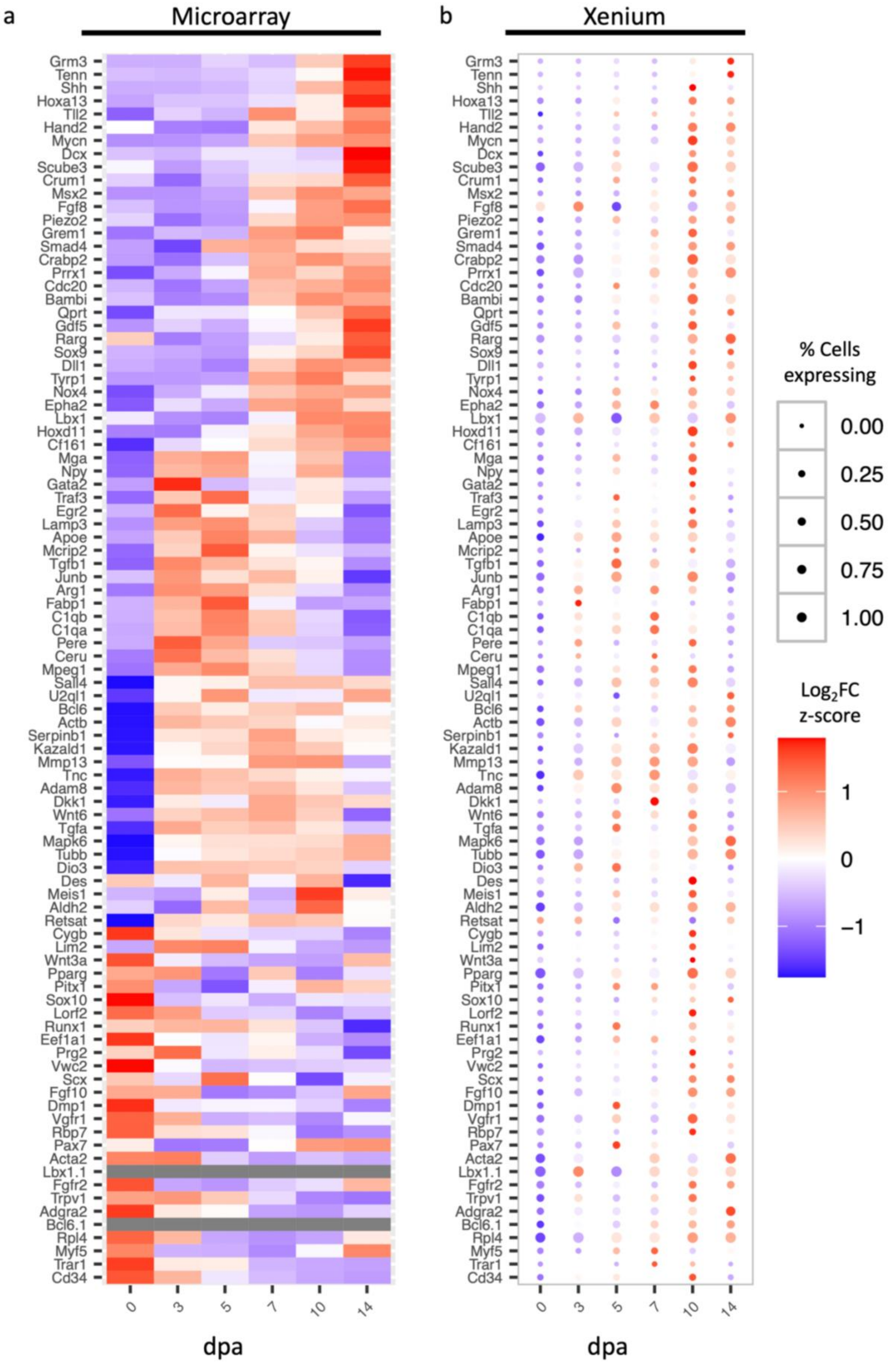
a
Microarray
b
Xenium
Grm3
Tenn
Shh
Hoxa13
Tll2
Hand2
Mycn
Dcx
Scube3
Crum1
Msx2
Fgf8
Piezo2
Grem1
Smad4
Crabp2
Prrx1
Cdc20
Bambi
Qprt
Gdf5
Rarg
Sox9
Dll1
Tyrp1
Nox4
Epha2
Lbx1
Hoxd11
Cf161
Mga
Npy
Gata2
Traf3
Egr2
Lamp3
Apoe
Mcrip2
Tgfb1
Junb
Arg1
Fabp1
C1qb
C1qa
Pere
Ceru
Mpeg1
Sall4
U2ql1
Bcl6
Actb
Serpinb1
Kazald1
Mmp13
Tnc
Adam8
Dkk1
Wnt6
Tgfa
Mapk6
Tubb
Dio3
Des
Meis1
Aldh2
Retsat
Cygb
Lim2
Wnt3a
Pparg
Pitx1
Sox10
Lorf2
Runx1
Eef1a1
Prg2
Vwc2
Scx
Fgf10
Dmp1
Vgfr1
Rbp7
Pax7
Acta2
Lbx1.1
Fgfr2
Trpv1
Adgra2
Bcl6.1
Rpl4
Myf5
Trar1
Cd34
0
3
5
7
10
14
dpa
% Cells expressing
0.00
0.25
0.50
0.75
1.00
Log2FC z-score
1
0
−1

**Supplementary Figure 17. Comparative analysis of microarray from flow-sorted CT-derived cells and Xenium blastema-fibroblasts cluster.** Complete comparison of gene expression over time between microarray (**a**) dataset of flow-sorted CT-derived cells, and Xenium (**b**) *in-silico*-sorted blastema fibroblast clusters. In both the microarray heatmap and the Xenium dot plot, every row represents a gene, and every column represents a time point analyzed. Xenium's dot-plot size represents the percentage of blastema fibroblasts expressing the corresponding gene. In both the microarray and Xenium datasets, the color scale indicates fold-change in expression relative to the uninjured limb.

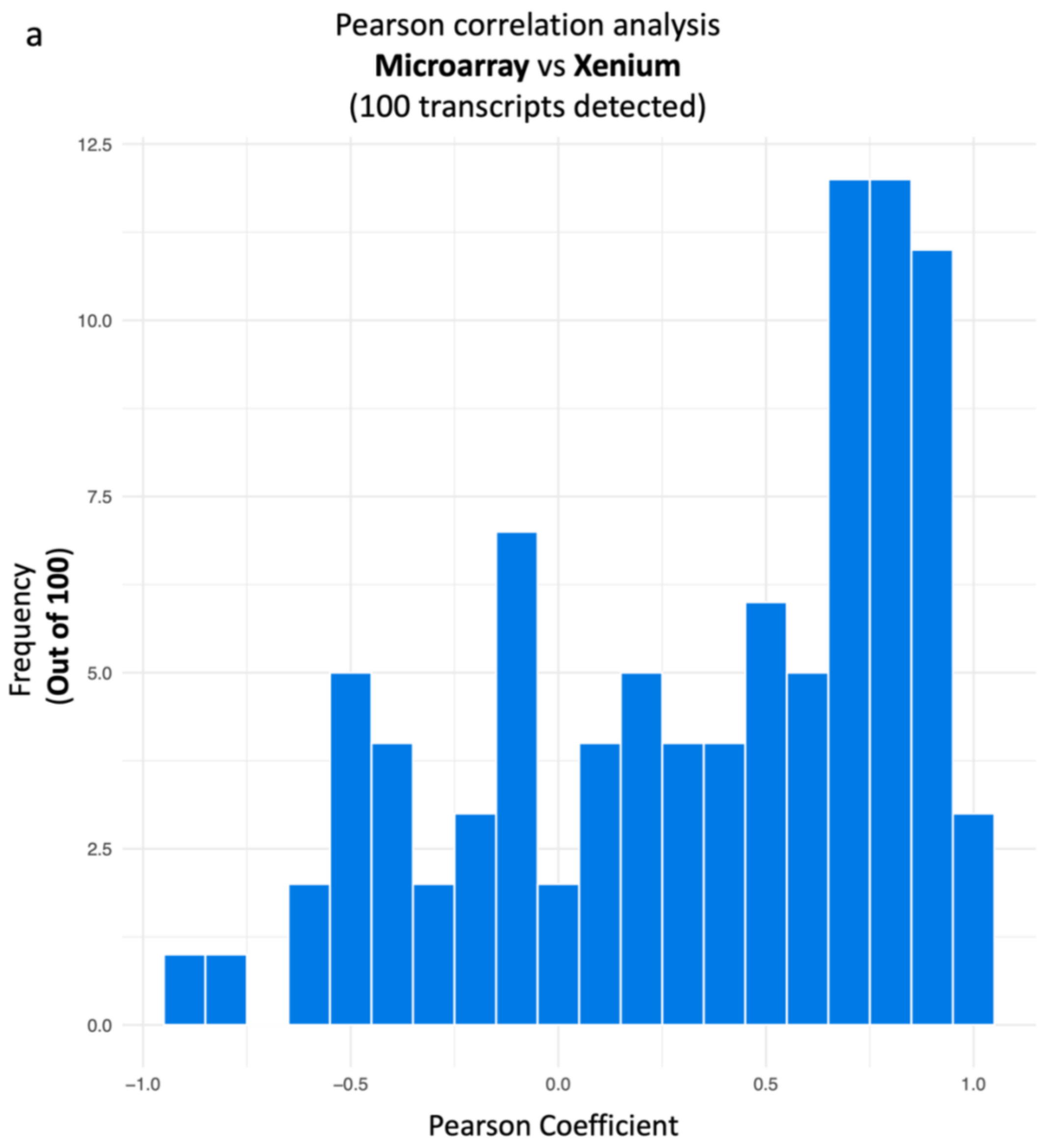


**Supplementary Figure 18. Pearson correlation analysis between microarray and Xenium technologies. a** Histogram depicting the frequency of Pearson correlation coefficients over the 100 gene comparisons performed between microarray and Xenium technologies. Negative correlation coefficients indicate that the detected gene tends to move in opposite directions.

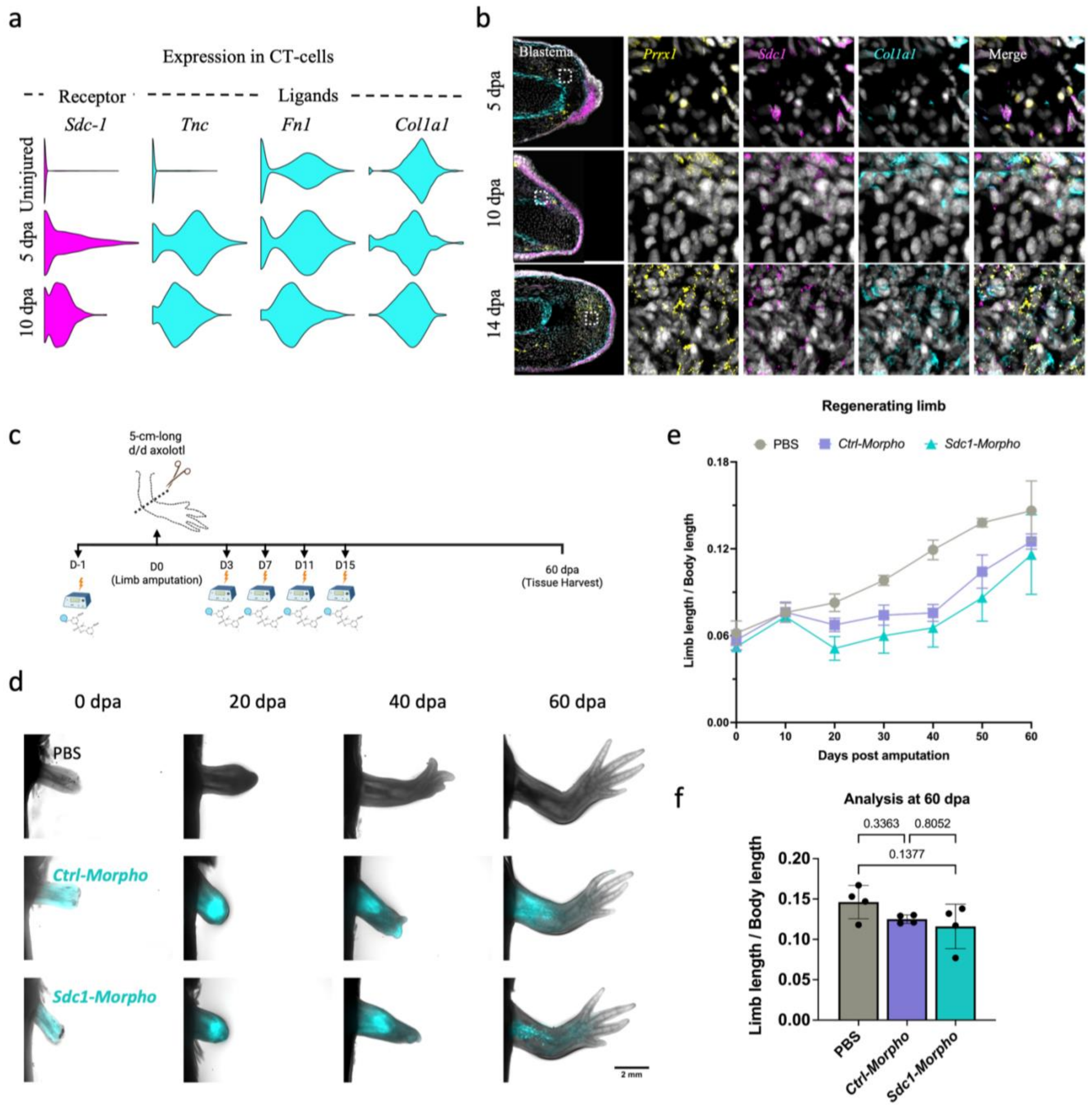


**Supplementary Figure 19. Morpholino electroporation delays limb regeneration. a**, Violin plots showing the expression levels of the *Sdc1* receptor and its associated ligands *Tnc*, *Fn1*, and *Col1a1* in CT-cell clusters across the uninjured limb, and 5-dpa, and 10-dpa blastema stages, as inferred by CellChat analysis. **b**, HCR validation of *Sdc1* and *Col1a1* transcript expression within *Prrx1*$^{+}$ CT cells at 5, 10, and 14 dpa. Left column shows longitudinal sections of the full blastema; scale bar, 1 mm. Remaining columns show magnified views of the boxed regions indicated by the white dotted line; scale bar, 100 μm. **c**, Schematic representation of the experimental design for the repeated electroporation of PBS, Sdc1-morpholino, and Control(Ctrl)-morpholino. **d**, Stereoscopic images of regenerating limbs at 0, 20, 40, and 60 dpa for each condition. Cyan fluorescence corresponds to the fluorescein tag present in all morpholino oligonucleotides. Scale bar = 2 mm. **e-f**, Quantitative assessment of the limb-length/body-length ratio over 60 days post amputation (e). The limb length was measured from the torso to the most distal point of the limb. Data are represented as mean ± SD (n = 4 for each group). Statistical comparisons at 60 dpa (f) were performed using a one-way ANOVA test, with p-values annotated above each comparison. Note that control morpholino produces a delay in regeneration similar to that of the Sdc1-morpholino, indicating that the observed phenotype reflects a response to the morpholino delivery rather than a Sdc1-specific knockdown.

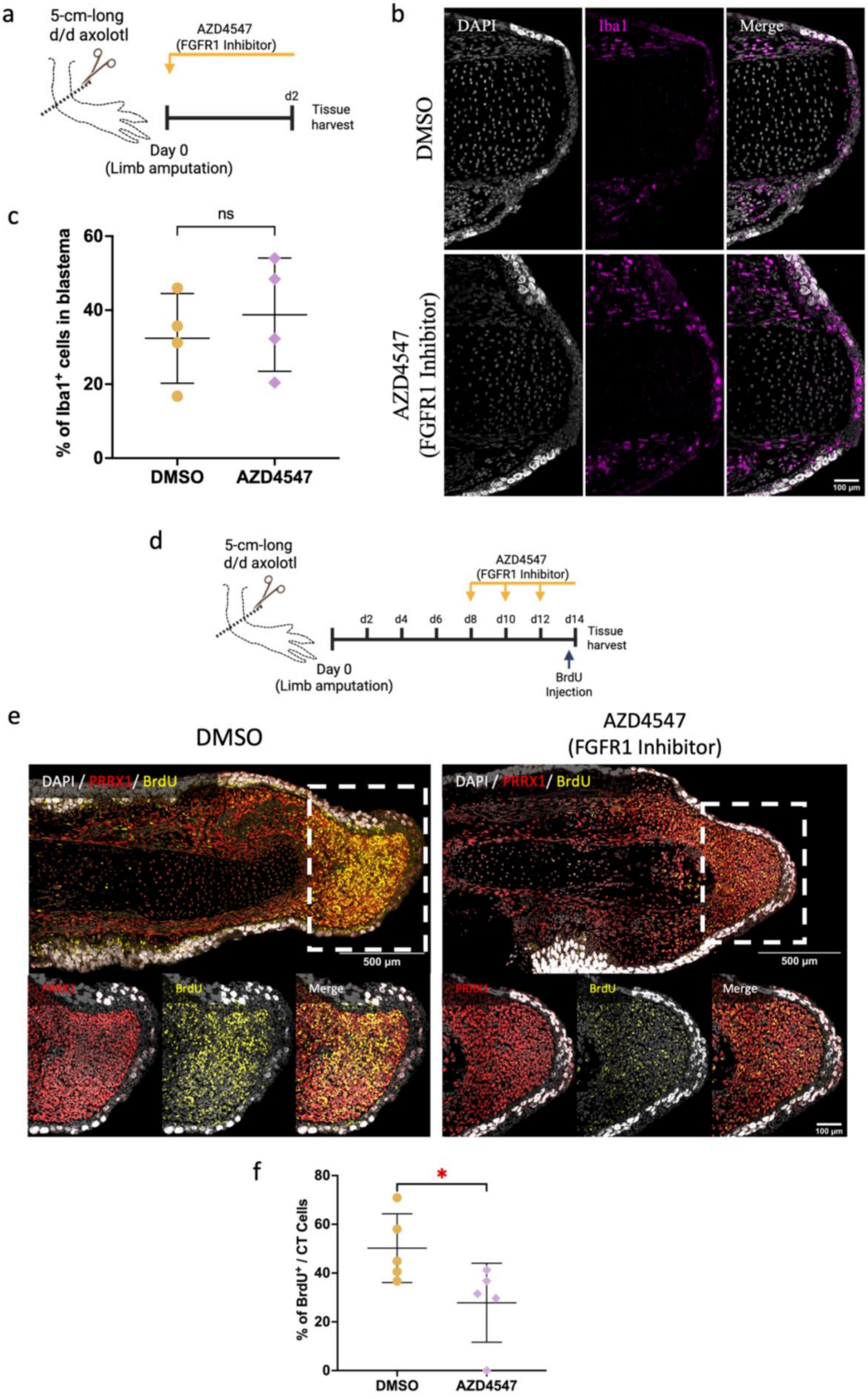
a
5-cm-long
d/d axolotl
AZD4547
(FGFR1 Inhibitor)
d2
Tissue
harvest
Day 0
(Limb amputation)
b
DAPI
Iba1
Merge
DMSO
AZD4547
(FGFR1 Inhibitor)
100 μm
c
ns
% of Iba1+ cells in blastema
60
40
20
0
DMSO
AZD4547
d
5-cm-long
d/d axolotl
AZD4547
(FGFR1 Inhibitor)
d2
d4
d6
d8
d10
d12
d14
Tissue
harvest
Day 0
(Limb amputation)
BrdU
Injection
e
DMSO
AZD4547
(FGFR1 Inhibitor)
DAPI / PRRX1/ BrdU
500 μm
PRRX1
BrdU
Merge
100 μm
f
*
% of BrdU+ / CT Cells
80
60
40
20
0
DMSO
AZD4547

**Supplementary Figure 20. FGFR1 inhibition does not affect early macrophage recruitment but reduces CT-cell proliferation in the blastema. a**, Schematic representation of the experimental design for assessment of early macrophage recruitment. AZD4547 treatment was performed by bathing axolotls for 2 days after limb amputation, at which point tissue was harvested. **b**, Confocal microscope images of longitudinal sections from 2-dpa blastemas in control DMSO and AZD4547-treated axolotls. DAPI (gray) marks all nuclei; Iba1 (magenta) marks macrophages. Quantification was performed within the most distal 500 μm region of the tissue, excluding bone and epidermis. **c**, Quantification of macrophage abundance, expressed as the percentage of Iba1$^{+}$ cells among all DAPI$^{+}$ cells within the quantified region. Data are represented as mean ± SD (n = 4 per condition). Statistical significance was determined using a two-way unpaired t-test; ns, not significant. **d**, Schematic representation of the experimental design for the assessment of CT-cell proliferation following FGFR1 inhibition. To avoid the complete block of blastema that results when AZD4547 treatment begins from day 0 (Figure 4f-g), we administered AZD4547 from 8 dpa (corresponding to the mid-bud blastema stage) through 14 dpa (coinciding with the late-bud blastema stage). BrdU was administered by intraperitoneal injection 6 hours before tissue harvest at 14 dpa. **e**, Confocal images of longitudinal blastema sections stained for PRRX1 (red) and BrdU (yellow) in control DMSO and AZD4547-treated axolotls. Top panels represent the full limb section; scale bar, 500 μm. Bottom panels show magnified views of the boxed blastema regions indicated in white dashed boxes in the top panels; scale bar, 100 μm. **f**, Quantification of CT-cell proliferation, expressed as the percentage of BrdU+ cells among the total number of PRRX1$^{+}$ CT cells within the blastema region, measured from the amputation plane to the distal tip of the blastema, excluding the epidermis. Data are shown as mean ± SD (n = 5 per condition). Statistical significance was determined using an unpaired two-tailed t-test.

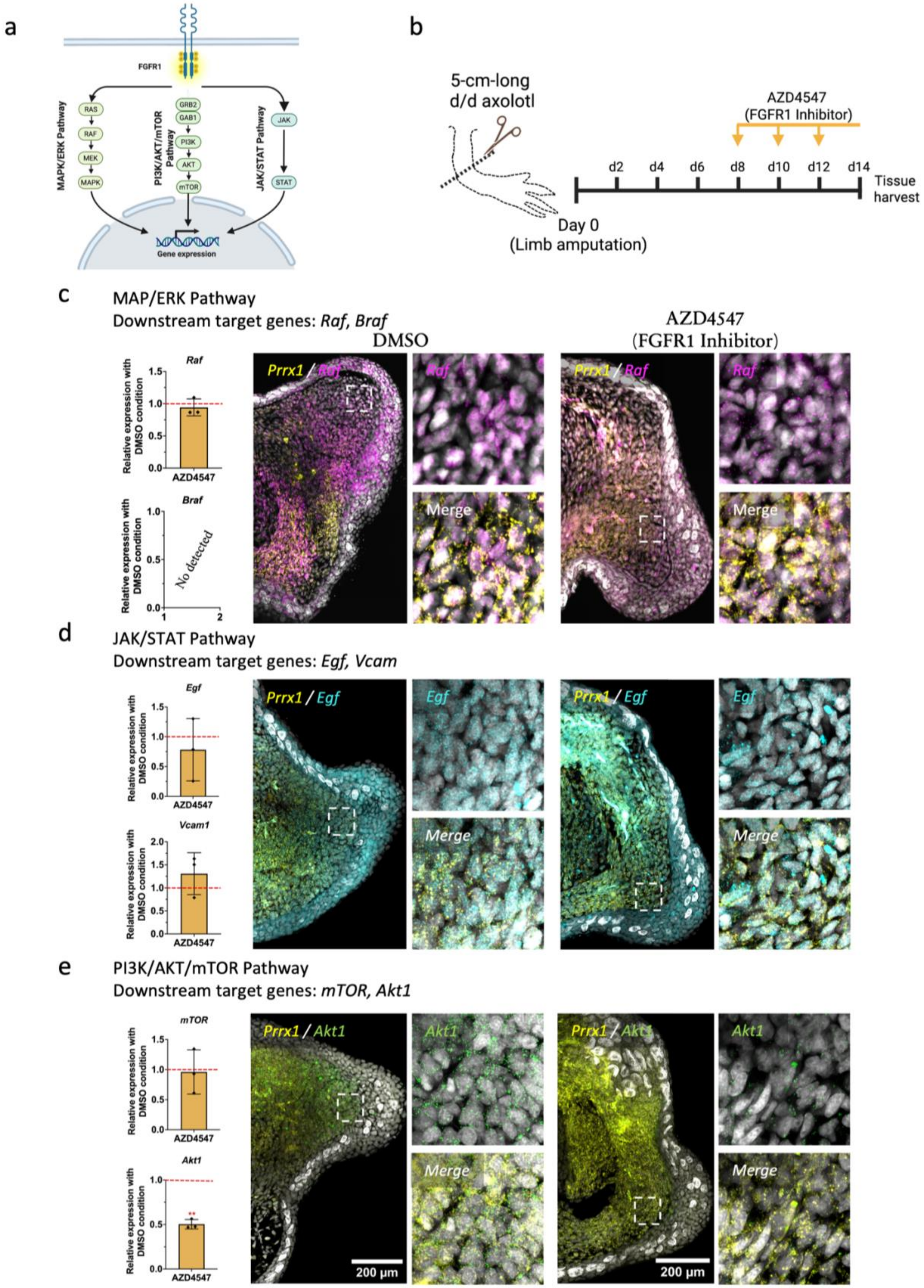
a
FGFR1
MAPK/ERK Pathway
RAS
RAF
MEK
MAPK
PI3K/AKT/mTOR Pathway
GRB2
GAB1
PI3K
AKT
mTOR
JAK/STAT Pathway
JAK
STAT
Gene expression
b
5-cm-long
d/d axolotl
AZD4547
(FGFR1 Inhibitor)
d2
d4
d6
d8
d10
d12
d14
Tissue harvest
Day 0
(Limb amputation)
c
MAP/ERK Pathway
Downstream target genes: Raf, Braf
DMSO
AZD4547
(FGFR1 Inhibitor)
Raf
Relative expression with DMSO condition
AZD4547
Braf
No detected
Prrx1 / Raf
Raf
Merge
d
JAK/STAT Pathway
Downstream target genes: Egf, Vcam
Egf
Vcam1
Prrx1 / Egf
Egf
Merge
e
PI3K/AKT/mTOR Pathway
Downstream target genes: mTOR, Akt1
mTOR
Akt1
Prrx1 / Akt1
Akt1
Merge
200 μm

315

**Supplementary Figure 21. FGFR1 inhibition downregulates *Akt1* expression in blastema CT cells, identifying the PI3K/AKT/mTOR pathway as the primary downstream effector. a**, Schematic representation of the three major downstream signaling branches activated by FGFR1: the MAPK/ERK, PI3K/AKT/mTOR, and JAK/STAT pathways. **b**, Schematic representation of the experimental design for assessing downstream pathway activation following FGFR1 inhibition. To avoid the complete block of blastema formation that results when AZD4547 treatment begins from day 0 (Figure 4f-g), we administered AZD4547 from 8 dpa (corresponding to a mid-bud blastema stage) through 14 dpa (coinciding with the late-bud blastema stage). **c-e**, Assessment of the pathway-specific downstream target-gene expression following FGFR1 inhibition, including the MAPK/ERK pathway targets *Raf* and *Braf* (c), the JAK-STAT pathway targets *Egf* and *Vcam1* (d), and the PI3K/AKT/mTOR pathway targets *mTOR* and *Akt1* (e). Left panels show RR-qPCR quantification of individual transcript levels normalized to the DMSO control condition. The red dashes indicate the control expression level. Data are represented as mean ± SD (n = 3 per condition). Statistical significance was determined using an unpaired two-tailed t-test; **p< 0.01. The middle and right panels show representative HCR images of blastema sections from control DMSO and AZD4547-treated axolotls, respectively, with double detection of *Prrx1* (yellow), and the pathway target of interest: *Raf* (magenta, c), *Egf* (cyan, d), and *Akt1* (green, e). Scale bar, 200 μm. Insets show magnified views of the region indicated by the white dashed box in each blastema section. Note that *Braf* transcripts were undetectable via RT-qPCR.

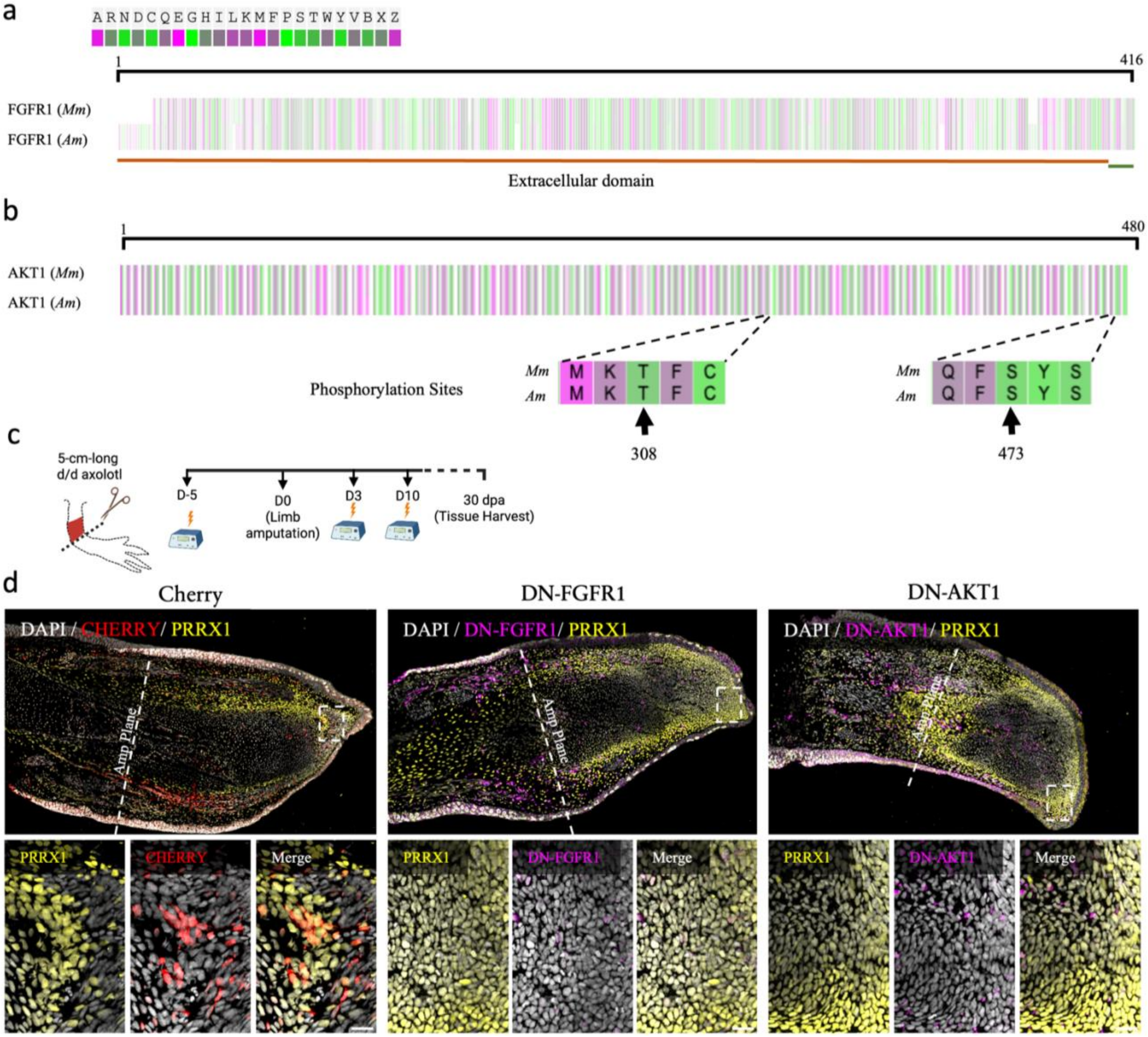
a
A R N D C Q E G H I L K M F P S T W Y V B X Z
1
416
FGFR1 (Mm)
FGFR1 (Am)
Extracellular domain
b
1
480
AKT1 (Mm)
AKT1 (Am)
Phosphorylation Sites
Mm M K T F C
Am M K T F C
308
Mm Q F S Y S
Am Q F S Y S
473
c
5-cm-long d/d axolotl
D-5
D0 (Limb amputation)
D3
D10
30 dpa (Tissue Harvest)
d
Cherry
DN-FGFR1
DN-AKT1
DAPI / CHERRY/ PRRX1
DAPI / DN-FGFR1/ PRRX1
DAPI / DN-AKT1/ PRRX1
Amp Plane
Amp Plane
PRRX1
CHERRY
Merge
PRRX1
DN-FGFR1
Merge
PRRX1
DN-AKT1
Merge

**Supplementary Figure 22. Dominant-negative (DN) FGFR1/AKT1-CT cells are preferentially retained near the amputation plane. a-b,** Amino acid sequence alignment between mouse (*Ms*) and axolotl (*Am*) FGFR1 (**a**) and AKT1 (**b**) proteins. In **a**, only the extracellular domain (bottom orange line) and transmembrane domain (bottom green line) of FGFR1 were used for the sequence alignment; the intracellular kinase domain is excluded as it is removed in the DN-FGFR1 construct. In **b**, the amplification (dotted lines) focuses on the regions flanking the two phosphorylation sites T308 and S473, which are mutated to T308A and S473A, respectively, in the DN-AKT1 construct. **c**, Schematic representation of the experimental design. DN-FGFR1, DN-AKT1, or Cherry-backbone control constructs were electroporated at the stylopod level five days prior to amputation to establish mosaic expression within the CT-cell population, with repeated electroporation at 3 and 10 dpa. Tissue was harvested at 30 dpa. **d**, Confocal images of longitudinal sections from electroporated limbs at 30 dpa, stained for PRRX1 (yellow), Cherry (red in Cherry-backbone control; magenta in DN constructs). Top panels show the full limb section; scale bar, 500 μm. Bottom panels show magnified views of the boxed region at the distal tip of the blastema; scale bar, 100 μm.
